# Identifying Modules of Cooperating Cancer Drivers

**DOI:** 10.1101/2020.06.29.168229

**Authors:** Michael I. Klein, Vincent L. Cannataro, Jeffrey P. Townsend, Scott Newman, David F. Stern, Hongyu Zhao

## Abstract

Identifying cooperating modules of driver alterations can provide biological insights to cancer causation and would advance the development of effective personalized treatments. We present Cancer Rule-Set Optimization (CRSO) for inferring the combinations of alterations that cooperate to drive tumor formation in individual patients. Application to 19 TCGA cancer types found a mean of 11 core driver combinations per cancer, comprising 2-6 alterations per combination, and accounting for a mean of 70% of samples per cancer. CRSO departs from methods based on statistical cooccurrence, which we demonstrate is a suboptimal criterion for investigating driver cooperation. CRSO identified well-studied driver combinations that were not detected by other approaches and nominated novel combinations that correlate with clinical outcomes in multiple cancer types. Novel synergies were identified in *NRAS*-mutant melanomas that may be therapeutically relevant. Core driver combinations involving *NFE2L2* mutations were identified in four cancer types, supporting the therapeutic potential of NRF2 pathway inhibition. CRSO is available at https://github.com/mikekleinsgit/CRSO/.

## 1 Introduction

Cells must deregulate multiple genetic pathways in in order to become cancerous. Most recent estimates are that 2-8 ‘hits’ are necessary for a precursor cell to become neoplastic [1, 2, 3]. Although identifying a small number of driver mutations among an excess of passenger mutations is a challenging problem, many methods already exist to do so. Generally, these methods identify genes mutated significantly more than the background mutation rate would predict and their output is a list of significant genes or regions found across a whole cohort. Well known examples include MutSigCV [4] and dNdScv [5] for identification of significantly mutated genes (SMGs), and GISTIC2 [6] for identification of significant somatic copy number variations (SCNVs).

Although useful in identifying known and novel cancer genes, these statistical enrichment methods do not attempt to identify co-occurring or mutually exclusive mutations. For example, activating mutations in *KRAS/NRAS* are frequently accompanied by loss of function of *CDKN2A/B* in melanoma, lung, pancreatic and other cancers. A major driving force behind the co-occurrence is that loss of the G1/S checkpoint (*CDKN2A/B*) is necessary to avoid oncogene induced senescence caused by *KRAS* oncogenic signaling [7, 8, 9]. Thus, mutations in these two genes cooperate to produce a pro-growth phenotype. Many methods can identify mutually exclusive candidate driver genes [10, 11, 12, 13, 14, 15, 16], but there are comparatively few that identify functionally relevant modules of cooccurring gene alterations in individual patients. Consequently, this extra layer of biological information and its possible relevance to therapy is not captured by any of the above methods and it would be beneficial to derive new approaches to identify groups of cooperating mutations.

From a biological perspective, cooccurrence of driver alterations clearly does occur [17, 18, 19, 20], and appears to be a requirement for carcinogenesis, as evidenced by the insufficiency of *BRAF* and *RAS* hotspot mutations to transform benign colon polyps and nevi into invasive carcinoma [21, 22]. However, statistical approaches to identifying cooccurrence have given mixed results. For example, Canisius *et al*. concluded that, after accounting for patient-specific mutation frequencies, there is no evidence of statistically significant cooccurrence of somatic mutations in cancer [23]. Conversely, Mina *et al*. did identify pairwise oncogenic synergies in multiple cancer types based on a model of sequential evolution using their SELECT algorithm [24]. Using mathematical modeling of cancer evolution, Mina *et al*. argue that statistically significant cooccurrence emerges only by conditioning on the sequence of alterations, consistent with the lack of traditional cooccurrence demonstrated by Canisius *et al*. [23]. Zhang *et al*. found evidence of statistically significant cooccurrence at the level of the pathway, but did not investigate at the gene-level [25]. Thus, current approaches can identify pairwise combinations of driver mutations in a subset of individuals. However, methods to detect combinations of three or more drivers, and methods that attempt to identify driver combinations within every sample in a cohort (or at least within a large proportion of samples) are needed.

Here, we describe Cancer Rule-Set Optimization (CRSO), a method to identify modules of cooperating alterations that are essential and collectively sufficient to drive cancer in individual patients. CRSO is developed as part of a theoretical framework that assumes the existence of specific combinations of two or more alterations called *rules* that cooperatively drive cancer if found in the same host cell. Rules are assumed to be minimally sufficient, meaning that exclusion of any of the alterations renders the remaining collection of alterations insufficient to drive cancer. CRSO seeks to find a collection of rules called a *rule set* that represent all of the unique minimal combinations that can explain all of the given tumors in the study population i.e., every sample is required to harbor all of the alterations in at least one of the rules. CRSO is robust to heterogenous mutational rates within the cohort as it uses alteration-specific passenger probabilities that reflect how likely specific observations would have occurred by chance. The output of CRSO can provide biological insights to cancer causation, as well as infer the likely driver combinations in individual patients. We show that CRSO can identify known and novel combinations of driver alterations in 19 tissue types from The Cancer Genome Atlas (TCGA) [26,27], and that some of these combinations correlate with clinical outcomes.

## 2 Results

### 2.1 CRSO overview

CRSO finds combinations of genomic alterations (referred to as *events*) that are predicted to cooperate to drive cancer in individual patients. The inputs into CRSO are two event-by-sample matrices: **D** and **P. D** is a binary alteration matrix, such that *D*_*ij*_ = 1 if event *i* occurs in sample *j*, and 0 otherwise. **P** is a continuous valued penalty matrix, where *P*_*ij*_ is the negative log of the probability of event *i* occurring in sample *j* by chance, i.e., as a passenger event. Three primary event types were considered: coding mutations identified as candidate drivers by dNdScv [5], as well as copy number amplifications and deletions identified as candidate drivers by GISTIC2 [6]. The version of CRSO presented here was restricted to SCNVs and coding mutations because candidate drivers for these alteration classes can be readily generated by statistical enrichment methods. However, CRSO models can include additional alteration types, such as gene-fusions and aneuploidies.

In order to calculate passenger probabilities, mutations and copy number variations were first represented as a categorical event-by-sample matrix, **M**. *M*_*ij*_ can take one of several values called *observation types*, that depend on on the type of event *i*. Mutation events take values in the set {*Z, HS, L, S, I*}, corresponding to wild-type, hotspot mutation, loss mutation, splice site mutation or in-frame indel (Section S1). Amplification events take values in {*Z, WA, SA*}, corresponding to wild-type, weak amplification and strong amplification, respectively. Similarly, deletion events take values in {*Z, WD, SD*}, corresponding to wild-type, weak deletion (hemizygous) and strong deletion (homozygous), respectively. Two additional event types, referred to as hybrid mutDels and mutAmps, were defined to represent genes that were identified by both dNdScv and GISTIC2 in the same tumor type (Methods 5.1, Figure 1A).

**Figure 1:**
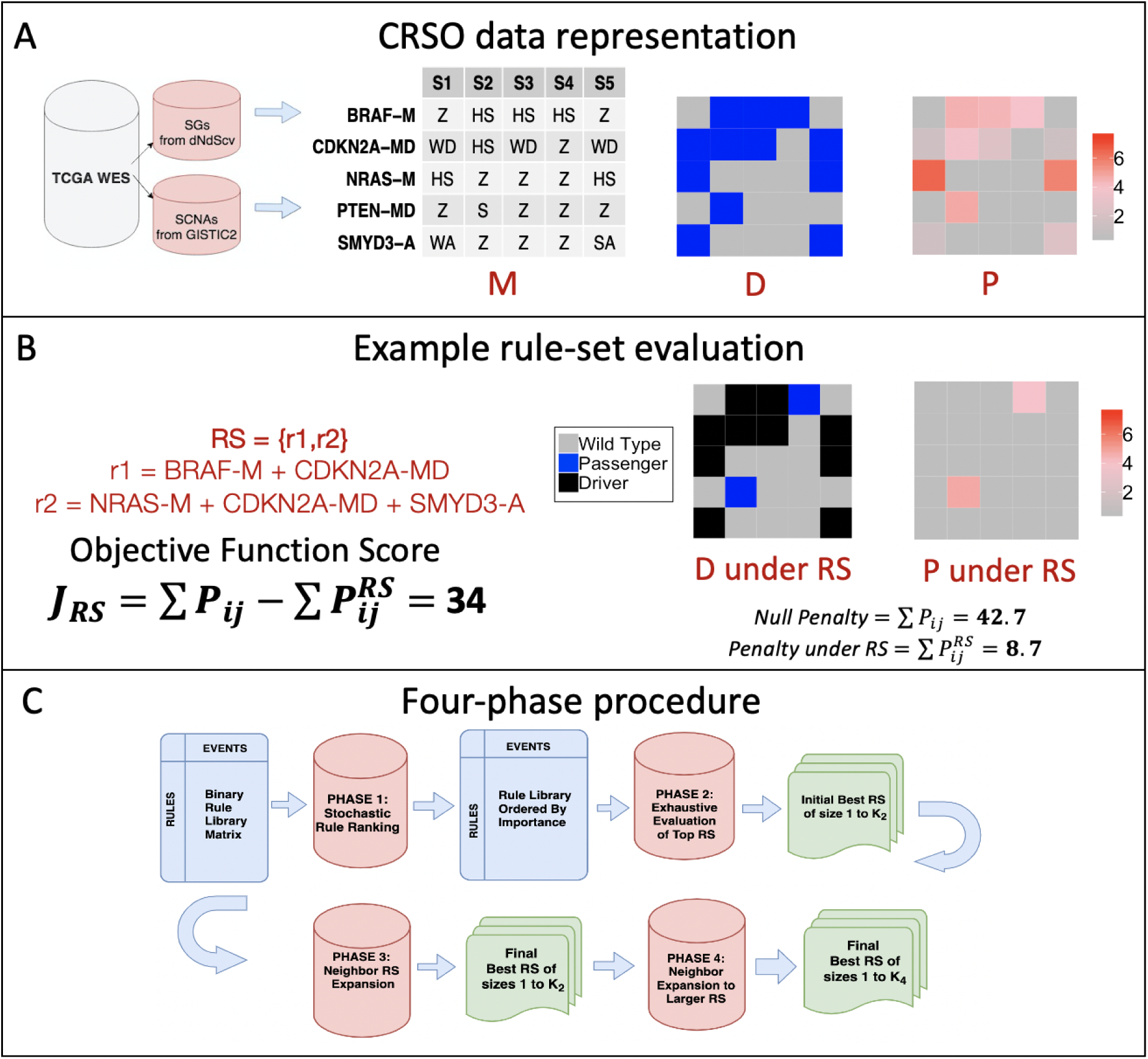
Visualization of CRSO using toy example that was extracted from TCGA melanoma (SKCM). A) Candidate driver events identified by dNdScv and GISTIC2. Data is first represented as categorical matrix **M**, where each event can take one of several observation types, depending on the event type. Event types are indicated by the suffix, -M for mutations, -A for amplifications and -MD for hybrid mutDels. Wild-type observations are represented as *Z* for all event types. Mutations are hotspot (HS), loss (L) splice site (S) or in-frame indel (I). Amplifications are either weak (WA) or strong (SA). Similarly, deletions are either weak (WD) or strong (SD). MutDel hybrid events can take values in any of the mutation or deletion observation types. Events in **M** are represented as a binary matrix **D** for making rule assignments. The penalty matrix **P** contains penalties for each possible observation based patient-specific, event-specific and observation-type specific passenger probabilities. B) Objective function calculated for an example rule set. Under the proposed rule set, the assigned events are designated as drivers, and the corresponding penalties are reduced to 0. C) Workflow of four phase procedure for identifying the best rule sets of size *K*, for a range of *K*s.

The CRSO model defines a **rule** as a collection of two or more events that drive tumors harboring all of the events in the collection. Rules are defined to be minimally sufficient, meaning that any strict subset of events within the rule are insufficient to cause cancer. A **rule set** is defined as a collection of rules that account for all minimally sufficient driver combinations within the entire cohort. Under a proposed rule set, every sample is assigned to at most one rule in the rule set. Two rules are defined to be *family members* if one rule is a strict subset of the other. Rule sets cannot contain any two rules that are family members because it is a contradiction for both rules to be minimally sufficient.

CRSO is a stochastic optimization procedure over the space of possible rule sets. When a sample is assigned to a rule in a rule set *RS*, the events that comprise the rule are considered to be drivers within that sample, and all of the remaining events in that sample are considered passengers. The **objective function score** for *RS, J* (*RS*), is defined to be the reduction in total statistical penalty under *RS* compared to the null rule set (Methods 5.2). Figure 1B shows an example of a rule set consisting of two rules applied to the miniature melanoma dataset. The total penalty is greatly reduced once samples are assigned to rules and the corresponding events are designated as drivers. The *coverage* of a rule set is the percentage of samples that can be assigned to at least one rule in the rule set. In Fig. 1B the coverage of *RS* is 80%.

The goal of CRSO is to find the rule set that achieves the best balance of objective function score and rule set size, which we call the **core rule set**. CRSO uses a **four-phase procedure** (Fig. 1C) to first find the highest scoring rule set of size *K* for *K* ∈ {1 … 40} (Methods 5.2.1). The core rule set is then determined from among all of the solutions of size *K*. A subsampling process is used to identify an expanded list of **generalized core rules** (GCRs). A confidence score is determined for each GCR to be the frequency of inclusion in the sub-sampled iterations. The subset of GCRs that have confidence levels above 50 comprise the **consensus GCRs** (con-GCR), and by definition cannot contain family members. The full CRSO methodology is presented Methods 5.2.

We applied CRSO to 19 cancer types obtained from TCGA (Table 1). The average number of events per cancer type was 86.4. There was a wide distribution of the number of candidate drivers per patient within individual cancers. In 17 out of 19 cancer types, the median number of total candidate drivers (SMGs plus SCNVs) was ≥ 6, and in 6 of them was ≥10 (Figure S1). CRSO identified an average of 11.1 core rules, covering an average of 70.3% of samples per cancer type. Of the 210 total core rules identified, 137 (65%), 47 (22%), 13 (6%), 12 (6%) and 1 (< 0.1%) consisted of 2, 3, 4, 5 and 6 events, respectively. The output of each CRSO run is a detailed report containing summaries and visual representations of the results (Supplementary S5).

**Table 1:** Summary of 19 TCGA tissues types investigated with CRSO

| Cancer | Full Name | Samples | Events (M/C) | RCT | Rules | Kc | Cov | MRS | con-GCRs |
| --- | --- | --- | --- | --- | --- | --- | --- | --- | --- |
| BLCA | Bladder urothelial CA | 392 | 113 (47/66) | 0.033 | 1730 | 14 | 76 | 2.21 | 15 |
| BRCA | Breast invasive CA | 963 | 90 (28/62) | 0.03 | 449 | 11 | 58 | 2 | 13 |
| CESC | Cervical and endocervical cancers | 191 | 82 (23/59) | 0.031 | 173 | 12 | 53 | 2.08 | 15 |
| COAD | Colon AD | 362 | 97 (35/62) | 0.039 | 1792 | 14 | 81 | 3.64 | 10 |
| ESCA | Esophageal CA | 184 | 92 (12/80) | 0.054 | 1894 | 13 | 84 | 3.23 | 10 |
| GBM | Glioblastoma multiforme | 273 | 78 (15/63) | 0.033 | 186 | 11 | 77 | 2.27 | 10 |
| HNSC | Head and neck squamous cell CA | 505 | 94 (25/69) | 0.032 | 926 | 10 | 68 | 2.5 | 7 |
| KIRC | Kidney renal clear cell CA | 433 | 39 (13/26) | 0.014 | 66 | 10 | 52 | 2 | 11 |
| LGG | Brain lower grade glioma | 513 | 67 (19/48) | 0.031 | 210 | 5 | 65 | 2.4 | 6 |
| LIHC | Liver hepatocellular CA | 366 | 75 (18/57) | 0.03 | 146 | 12 | 56 | 2 | 16 |
| LUAD | Lung AD | 478 | 93 (22/71) | 0.031 | 291 | 10 | 60 | 2.1 | 11 |
| LUSC | Lung squamous cell CA | 178 | 112 (37/75) | 0.045 | 1588 | 14 | 83 | 2.93 | 10 |
| OV | Ovarian serous cystAD | 455 | 77 (6/71) | 0.075 | 1990 | 12 | 86 | 3.08 | 12 |
| PAAD | Pancreatic AD | 126 | 65 (11/54) | 0.032 | 549 | 6 | 79 | 2.67 | 3 |
| PRAD | Prostate AD | 492 | 73 (13/60) | 0.03 | 301 | 10 | 51 | 2.5 | 6 |
| READ | Rectum AD | 120 | 67 (13/54) | 0.042 | 1260 | 12 | 89 | 3.17 | 9 |
| SKCM | Skin cutaneous melanoma | 290 | 71 (20/51) | 0.031 | 197 | 10 | 69 | 2 | 9 |
| STAD | Stomach AD | 391 | 121 (37/84) | 0.031 | 1690 | 13 | 64 | 2.46 | 14 |
| UCEC | Uterine corpus endometrial CA | 242 | 136 (39/97) | 0.037 | 1515 | 11 | 84 | 2.36 | 7 |

**Full Name:** AD short for adenocarcinoma, CA short for carcinoma. **Events** total number of events as determined by dNdScv and Gistic2, (M/C) is number of mutations and CNVs. **RCT** rule coverage threshold, chosen to be .03 or the smallest threshold satisfied by at most 2000 rules, except in the case of KIRC. **Rules** starting library size. **Kc** core rule set size. **Cov** core rule set coverage. **MRS** mean core rule size. **con-GCRs** consensus GCR sizes.

### 2.2 CRSO performance on simulated data

In order to assess CRSO’s ability to detect known driver combinations, we initially used simulations with known ground truth rule sets. Ten simulated datasets with ground truth rule sets of size *ntr* were randomly generated for each *ntr* ∈ {2 … 20}, for a total of 190 simulations. Each dataset contained 100 events and 400 samples, chosen to approximate the mean number of events and samples across the TCGA datasets. Passengers rates for events and samples were sampled from a pooled distribution of real data passenger rates (Supplementary S6.1, Figure S2). We evaluated CRSO using 3 metrics: sensitivity, specificity, and rule assignment accuracy. Sensitivity was calculated to be the percentage of ground truth rules included in the consensus GCRs (con-GCRs). Specificity was calculated to be the percentage of con-GCRs that are in the ground truth rule set. Assignment accuracy was the percentage of samples identically assigned in the core RS assignment and the ground truth assignment.

The mean sensitivity was at least 0.85 for all *ntr* ∈ {2 … 20}, and was ≥ 0.90 for 16/19 *ntr* values (Table S3, Figure 2A). The mean specificity was 0.73 for *ntr* = 2 and was ≥ 0.96 for the other 18 true rule set sizes (Table S3, Figure 2B). The lowest mean accuracy was observed for *ntr* = 2, at 0.76, and was above 0.80 for all other *ntr* values. Starting with *ntr* = 3, for which the mean sensitivity was 1.0, and the mean accuracy was 0.98, the general trend was that sensitivity and accuracy decreased as *ntr* increased. For *ntr* = 2, 3 simulations performed poorly with accuracies below 0.4 and the other 7 simulations all had accuracies greater than 0.90 (Figure S3), suggesting that CRSO may be susceptible to over-fitting when the number of true rules is only 2.

**Figure 2:**
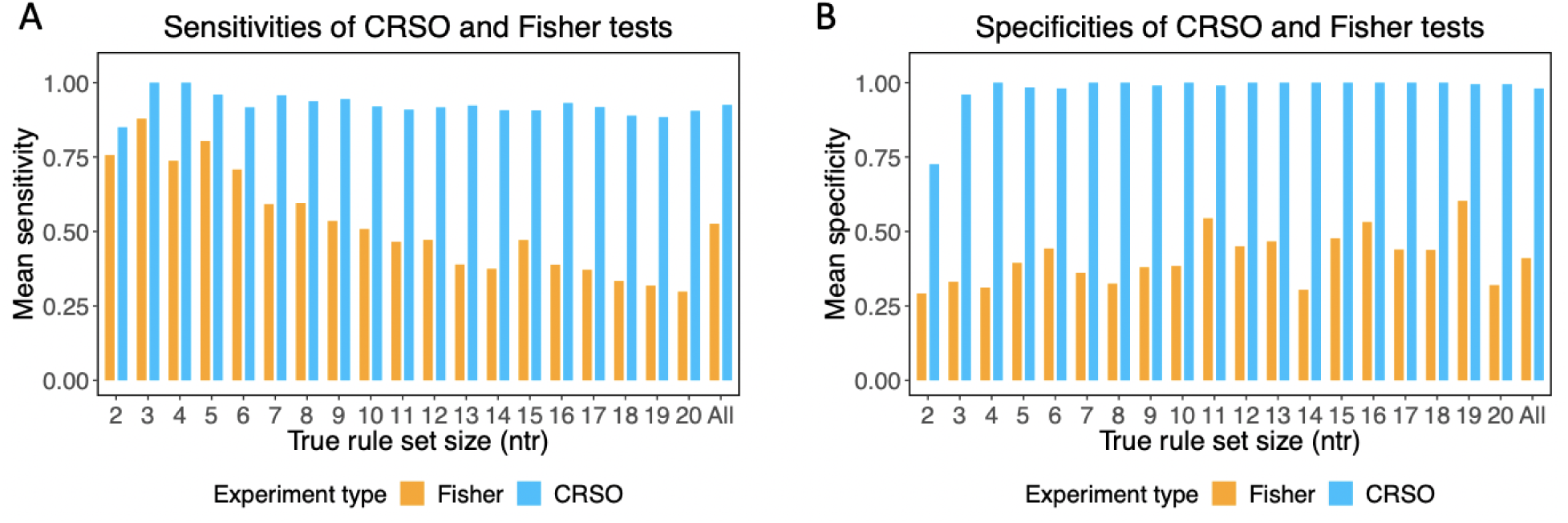
Performance of CRSO and Fisher tests applied to 190 ground truth simulations. Results are grouped according to ground truth rule set size (NTR). The group denoted ‘All’ is the global mean over all NTR. (A) Mean sensitivities of CRSO predictions evaluated against ground truth rules (light blue bars) and Fisher test predictions evaluated against ground truth duos (orange bars). (B) Mean specificities of CRSO predictions evaluated against ground truth rules (light blue bars) and Fisher predictions evaluated against ground truth duos (orange bars).

Across 190 simulated datasets, CRSO was able to navigate an intractable search space of possible rule sets (e.g., for a library of 1500 rules, there are > 10^45^ possible rule sets of size ≤ 20) and identify ground truth rules comprising between 2-6 events with high specificity and high sensitivity. We compared CRSO’s performance on the simulated dataset with that of pairwise cooccurrence tests-a popular strategy for detecting driver-gene cooperation. For each simulation, the Fisher exact test was performed for all pairs of events, and the detected cooccurrences were compared to the ground truth duos, i.e., the superset of all pairs of events that cooccur within at least one ground truth rule. Events with 0 or 1 occurrence were excluded. Multiple-hypothesis correction for each simulation experiment was performed using a false discovery rate of 5%. Sensitivities were calculated as the percentage of true pairwise interactions that were identified as statistically co-occurrent. Specificities were calculated as the percentage of identified pairwise interactions that were part of at least one ground truth rule.

The mean sensitivity achieved by the Fisher test 0.53 ± 0.18 over all *ntr*, and ranged from a high of 0.88 for *ntr* = 3, to a low of 0.30 for *ntr* = 20 (Figure 2A). The mean specificity achieved by the Fisher test was 0.41 ± 0.09 over all *ntr*, and ranged from a low of 0.29 for *ntr* = 2, to a high of 0.60 for *ntr* = 19 (Figure 2B). The mean sensitivities and specificities achieved by CRSO outperformed those of the Fisher test for all *ntr*, with an average sensitivity improvement of 0.4 and an average specificity improvement of 0.57 (Figure 2).

To better understand the limitations of pairwise cooccurrence tests, we applied the Fisher exact test to a version of the simulations having uniform passenger rates across samples and events, and to a version of the simulations with zero passenger events. Figure S2 shows an example simulation with zero passenger events (**D_z_**), uniformly distributed passenger events (**D_u_**), and realistically distributed noise rates that were used for the CRSO performance evaluation (**D_r_**). The mean sensitivity for **D_u_** was 0.47 ± 0.20 across all *ntr*, and was comparable to the mean sensitivity for **D_r_** (Figure S5A). The mean specificity for **D_u_** was 0.96 ± 0.03 across all *ntr*, and was far superior to the those for **D_r_** (Figure S5B). The specificity results suggest that unaccounted-for heterogeneity in passenger rates across samples and events lead to an excess of false-positives, consistent with the findings of Canisius *et al*. [23]. The specificity was always 1.0 for **D_z_**, since it is impossible to have false positives in the absence of noise. The mean sensitivity for **D_z_** was 0.76 ± 0.11 across all *ntr*, and ranged from a high of 1.0 for *ntr* = 2, to a low of 0.64 for *ntr* = 20 (Figure 2A). The sensitivities for simulations without any noise show that many false negatives result directly from the ground truth rule set structure.

### 2.3 Application to TCGA melanoma data

We next present results from 290 TCGA cutaneous melanomas (SKCM) (Figure 3). We use this as an exemplar dataset as SKCM is a heterogeneous cancer type with some known driver combinations [28, 29, 30, 31]. We used as input, 20 SMGs from dNdScv, 34 SCNV deletions and 20 SCNV amplifications from GISTIC2 narrow peak calls. Three genes, *CDKN2A, PTEN* and *B2M*, were represented as hybrid “mutDel” events as they were both significantly mutated and deleted in the dataset (Sections S1, S2 and S3).

**Figure 3:**
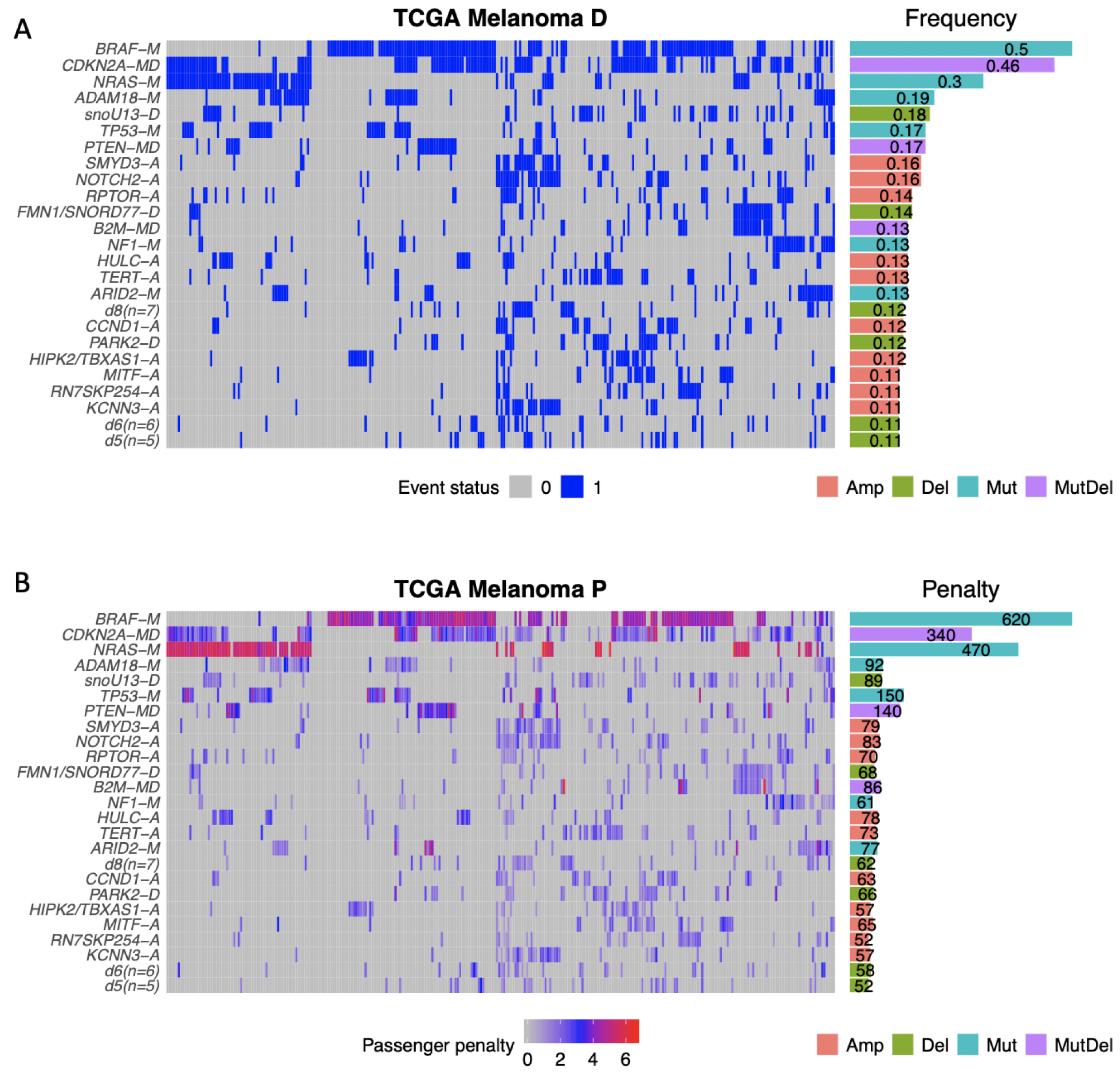
CRSO Representation of TCGA Melanoma Dataset. A) Binary representation of top-25 most frequent events. Horizontal bars on the right show event frequencies across the population. B) Penalty matrix for top-25 most frequent events. Horizontal bars on the right show total event penalties. Event types indicated by suffixes: -A for amplifications, -D for deletions, -M for mutations, and -MD for mutDel hybrid events.

The candidate rule library contained 197 rules that satisfied the coverage threshold of 3%, of which 165 consisted of two events and 32 consisted of three events. The core rule set was determined to be the best rule set of size *K* = 10, as this was the smallest rule set that exceeded both the performance and coverage thresholds of at least 90% (Figure S6A-B). There were no valid rule sets that satisfied the minimum samples assigned (*msa*) threshold of 9 samples for *K* ≥ 18. Figure 4 shows the optimal assignment under the core RS and the corresponding reduction in penalty. Thirty percent of the SKCM patients do not satisfy any of the core rules, indicated by the grey color bar in Fig. 4A.

**Figure 4:**
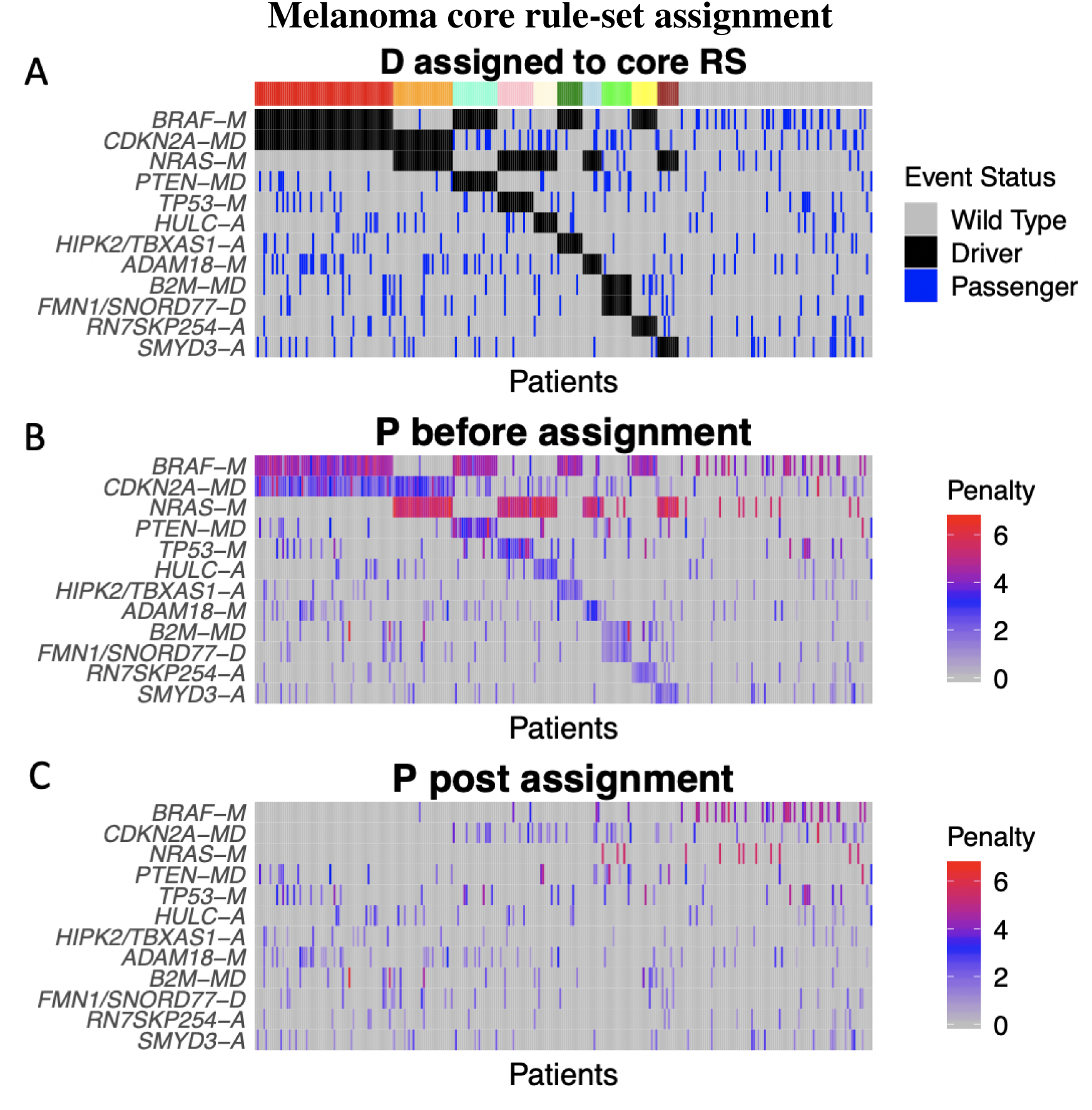
A) Heatmap of the binary alteration matrix **D** under core rule set assignment. Events are ordered by frequency. Samples are ordered according to rule set membership, as indicated by the color bar. The right-most group (red bar), are not assigned to any rule. For each sample, assigned events are designated as drivers (black), and unassigned events (blue) are assumed to be passengers. B–C) Heatmaps of the passenger penalty matrix **P** before and after assignment to the core rule set.

Generalized core (GC) analysis using 100 subsampling iterations identified 9 con-GCRs (Methods 5.2.2). Although all of the melanoma con-GCRs contained 2 events, comparison of the GCRs with GC duos reveals that some con-GCRs appear as part of larger rules in some GC iterations (Figure S7). For example, *BRAF-M*+*CDKN2A-MD* and *BRAF-M*+*PTEN-MD* are observed as core rules in 83% and 84% of GC iterations, respectively, but are both observed as core duos in 100% of GC iterations. The difference in GCD and GCR confidence scores indicates that CRSO is 100% confident that *BRAF-M*+*PTEN-MD* and *BRAF-M*+*CDKN2A-MD* are both essential combinations in a subset of melanoma patients, but is approximately 84% confident that these duos are independently sufficient to produce melanomas.

The core RS and con-GCRs can be considered complementary best rule sets, with the core RS providing the single best performing RS and corresponding assignment over the full dataset, and the con-GCRs providing a robust set of rules with quantified confidence scores. The union of the melanoma core RS and con-GCRs comprise 11 distinct rules, and are dominated by rules that contain either *BRAF* or *NRAS* mutations (Table 2). Of the 11 rules, 5 contain *BRAF*, 5 contain *NRAS* and only 1 rule, *B2M-MD*+*FMN1/SNORD77-D*, contains neither. Hotspot mutations in *BRAF* and *NRAS* define the two major subtypes of melanoma that are mutually exclusive, with 50% of patients harboring *BRAFV600E* mutations and 30% of patients harboring *NRAS* hotspot mutations [32]. CRSO prioritized rules containing these events because it is improbable that these highly recurrent hotspot mutations would have happened by chance.

**Table 2:** Melanoma Core RS and Consensus Generalized Core Rules (conGCR)

| Rule | Core Type | P1 Rank | Confidence | Coverage | SJ | Fraction Assigned |
| --- | --- | --- | --- | --- | --- | --- |
| *NRAS-M + TP53-M* | Both | 4 | 100.00 | 6.6% (r=29) | 168 (r=8) | 0.89 |
| *BRAF-M + RN7SKP254-A* | Both | 11 | 98.00 | 6.6% (r=32) | 137 (r=21) | 0.63 |
| *BRAF-M + HIPK2/TBXAS1-A* | Both | 6 | 97.00 | 7.9% (r=13) | 161 (r=10) | 0.52 |
| *CDKN2A-MD + NRAS-M* | Both | 2 | 84.00 | 14% (r=2) | 327 (r=2) | 0.68 |
| *BRAF-M + PTEN-MD* | Both | 3 | 84.00 | 11% (r=3) | 227 (r=3) | 0.68 |
| *BRAF-M + CDKN2A-MD* | Both | 1 | 83.00 | 26% (r=1) | 524 (r=1) | 0.86 |
| *HULC-A + NRAS-M* | Both | 5 | 71.00 | 6.6% (r=28) | 167 (r=9) | 0.58 |
| *B2M-MD + FMN1/SNORD77-D* | Both | 10 | 66.00 | 9% (r=6) | 144 (r=36) | 0.54 |
| *ADAM18-M + NRAS-M* | Core | 9 | 48.00 | 6.9% (r=20) | 146 (r=11) | 0.45 |
| *NRAS-M + SMYD3-A* | Core | 15 | 40.00 | 5.5% (r=40) | 131 (r=20) | 0.62 |
| *BRAF-M + TP53-M* | conGCR | 8 | 51.00 | 8.3% (r=9) | 156 (r=7) | - |

**Core Type** Core RS, conGCR or Both. **Confidence** is generalized core rules confidence level, conGCRs defined as rules with confidence above 50. **P1 Ranks** phase 1 ranking of rule. **Coverage** is the percentage of samples that cover the rule (and ranking among all the rules in the rule library), **SJ** is the single rule performance (and ranking), **FracA** is the fraction of covered samples that are assigned to the rule. Only applies to core rules.

#### 2.3.1 Expected melanoma combinations detected by CRSO

We categorized the 11 rules as either expected (5 rules) or potentially novel (6 rules) based on literature review. The 5 expected rules involve combinations of either *BRAF* or *NRAS* with well-studied tumor suppressors: 1) *NRAS-M*+*TP53-M*, 2) *BRAF-M*+*TP53-M*, 3) *NRAS-M*+*CDKN2A-MD*, 4) *BRAF-M*+*CDKN2A-MD*, and 5) *BRAF-M*+*PTEN-MD*. The first 4 of these rules are instances of cooccurring MAPK3 pathway activation with P53 inactivation-a synergy that is known promote carcinogenesis [30, 31]. Evidence in multiple cancer types supports the cooperation between activating mutations in *KRAS* and loss of the G1/S checkpoint by inactivation of *CDKN2A* or *TP53* [7, 8, 9]. CRSO identified *KRAS-M*+*TP53-M* as a consensus GCR in 3 other cancer types, LUAD, PAAD and STAD, and identified *KRAS-M*+*CDKN2A-MD* as a con-GCR in LUAD and PAAD. Cooccurrence of *BRAFV600E* and *CDKN2A* loss defines a subset of pediatric brain tumors that are responsive to combined treatment with *BRAF* and *CDK4/6* inhibitors [33]. Unlike *CDKN2A* and *TP53, PTEN* was inferred to exclusively cooperate with *BRAF*. Supporting the minimal sufficiency of *BRAF-M*+*PTEN-MD*, cooccurrence of *BRAFV600E* and *PTEN* loss were shown to induce metastatic melanoma in mouse models, while *BRAFV600E* alone only produced benign nevi in the mice [28]. *PTEN* loss is a mechanism of primary and acquired resistance to *BRAF* inhibitors, inspiring the development of combination strategies co-targeting the *BRAF/MEK* and *PI3K/AKT* pathways in tumors harboring both *BRAFV600E* and *PTEN* loss [29, 34, 35]

These 5 rules exemplify CRSO’s ability to identify combinations with experimental evidence of being minimally sufficient to drive cancer, and with established utility as biomarkers and sources of rational combination strategy. Although these combinations are not novel, CRSO prioritized these rules without any prior biological knowledge. By contrast, 2 published methods for systematically identifying driver combinations were applied to the same TCGA melanoma dataset and did not identify any of these 5 combinations [24, 36]. Since most strategies are based on pairwise cooccurrence, the omission of these well studied synergies in melanoma further demonstrates the inappropriateness of statistical cooccurrence as a criteria for detecting driver combinations.

**Table 3:**
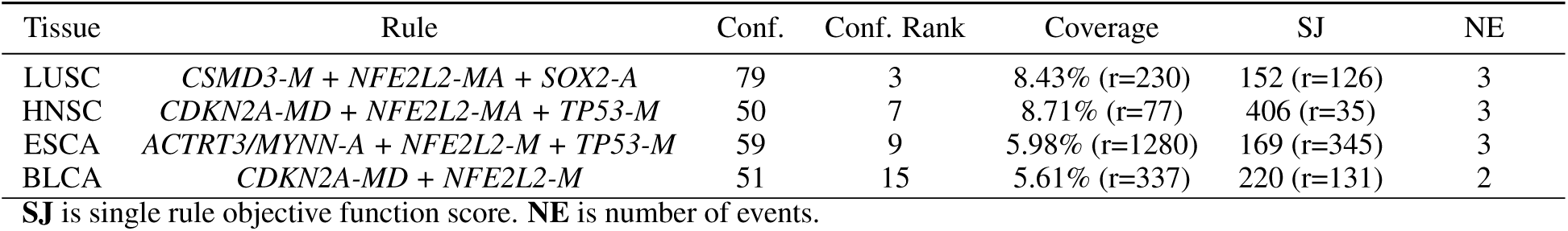
Consensus GCRs involving *NFE2L2*

#### 2.3.2 Novel melanoma combinations detected by CRSO

The other 6 rules in Table 2 involve lesser-known drivers and may represent novel biological subtypes of melanoma. *NRAS-M*+*HULC-A* was prioritized as both a con-GCR (conf. = 71), and as part of the core RS. *HULC* is a long non-coding RNA that has been identified as a driver of tumorigenesis in multiple cancers [37, 38, 39], including liver cancer, osteosarcoma, cervical cancer, pancreatic, stomach and ovarian cancers [40, 41, 42, 43, 44, 45]. *HULC* expression is a predictor of poor prognosis across multiple cancers [46, 47, 48, 49], and *HULC* silencing enhanced the effectiveness of chemotherapy in stomach cancer cell lines [44]. The involvement of *HULC* in the tumorigenesis of a subset of *NRAS* mutant melanomas appears to be unreported and merits further investigation as a possible novel discovery with therapeutic implications.

*NRAS-M*+*SMYD3-A* (conf. = 40) was identified as part of the CRSO core RS. *SMYD3* is a member of the histone lysine methyltransferases enzyme family, and upregulates transcription of a plethora of oncogenes in multiple cancer types, including *CDK2* and *MMP2* in hepatocellular carcinomas [50], *BCLAF1* in bladder cancer [51], androgen receptors in prostate cancer [52], *EGFR* in renal cell carcinoma [53] and *MYC* and *CTNNB1* in colon and liver cancers [54]. Mazur *et al*. showed that *SMYD3* methylation of *MAP3K2* leads to upregulation of MAP kinase signaling and promotes carcinogenesis in *RAS* mutated lung and pancreatic cancers [55]. The authors further showed that preventing *SMYD3* catalytic activity in mouse models with oncogenic *RAS* mutations inhibited tumor development [55]. This presents a direct biological link between *SMYD3* expression and *RAS* mutations, and nominates *SMYD3* as a drug target for *RAS* driven cancers. As with *HULC*, the possible role of *SMYD3* in melanoma appears to be unreported.

CRSO identified two synergies exclusive to *BRAF-M* with very high confidence: *BRAF-M*+*HIPK2/TBXAS1-A* (conf. = 97) and *BRAF-M*+*RN7SKP254-A* (conf. = 98). Although we annotated amplifications according to the GISTIC narrow peak regions, these two amplification events are part of large wide-peak regions containing many genes. The wide peak annotated by *HIPK2* and *TBXAS1* contains 72 genes, including *BRAF. BRAF* amplification is a mechanism of acquired resistance to *BRAF* inhibitors in *BRAFV600E* melanomas [56, 57, 58, 59]. We found that patients harboring *BRAF-M* and *HIPK2/TBXAS1-A* had shorter progression-free intervals (PFI) compared to patients harboring *BRAF-M* without *HIPK2/TBXAS1-A* (cox-ph *P* = 0.016, Table 4, Figure 5). Because TCGA tumors are primary and treatment-naive, this observation suggests that *BRAF* amplification and *BRAFV600E* may be a sufficient combination for tumor formation, and a cause of intrinsic resistance to *BRAF* inhibition.

**Table 4:**
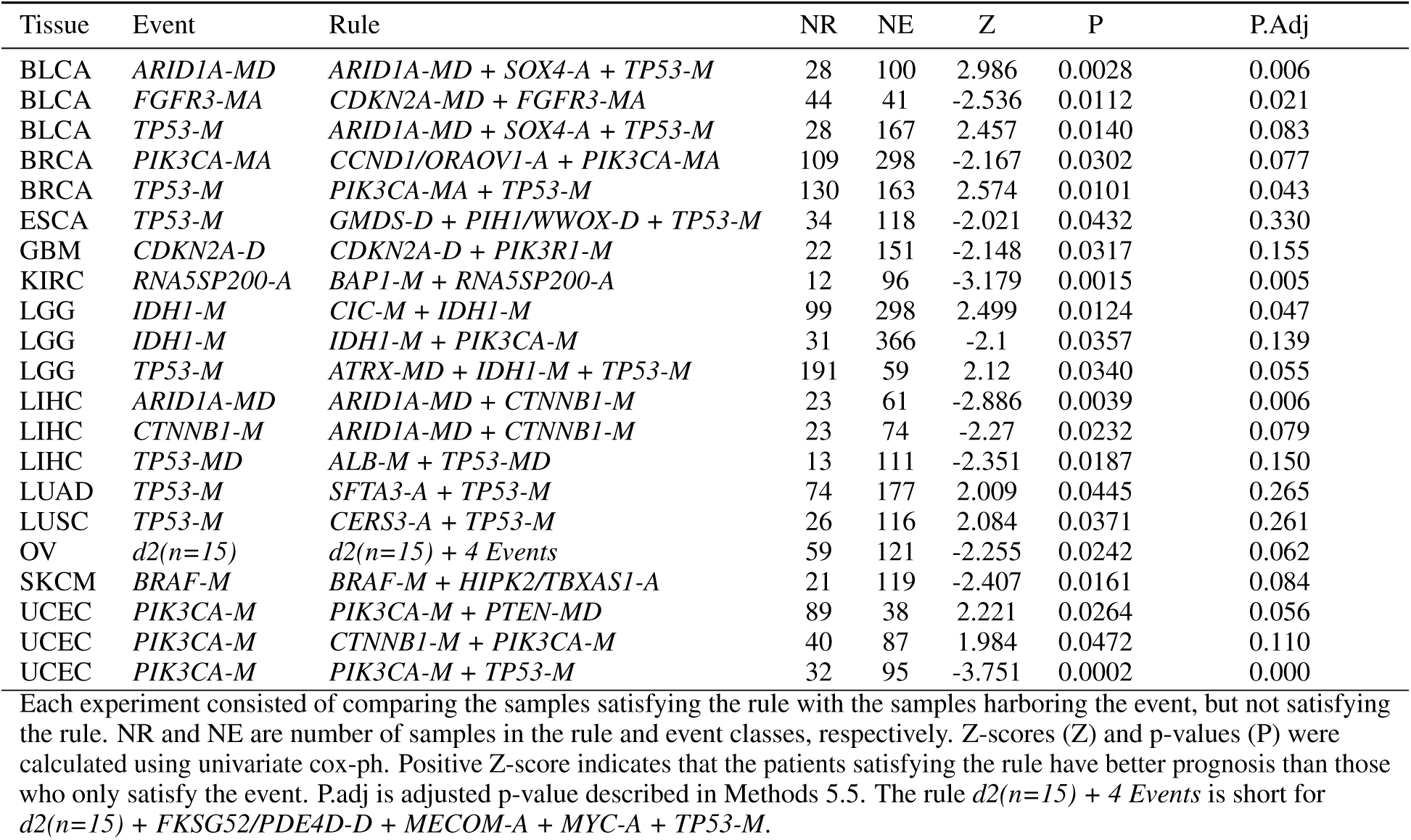
Significant PFI associations among consensus GCRs

**Figure 5:**
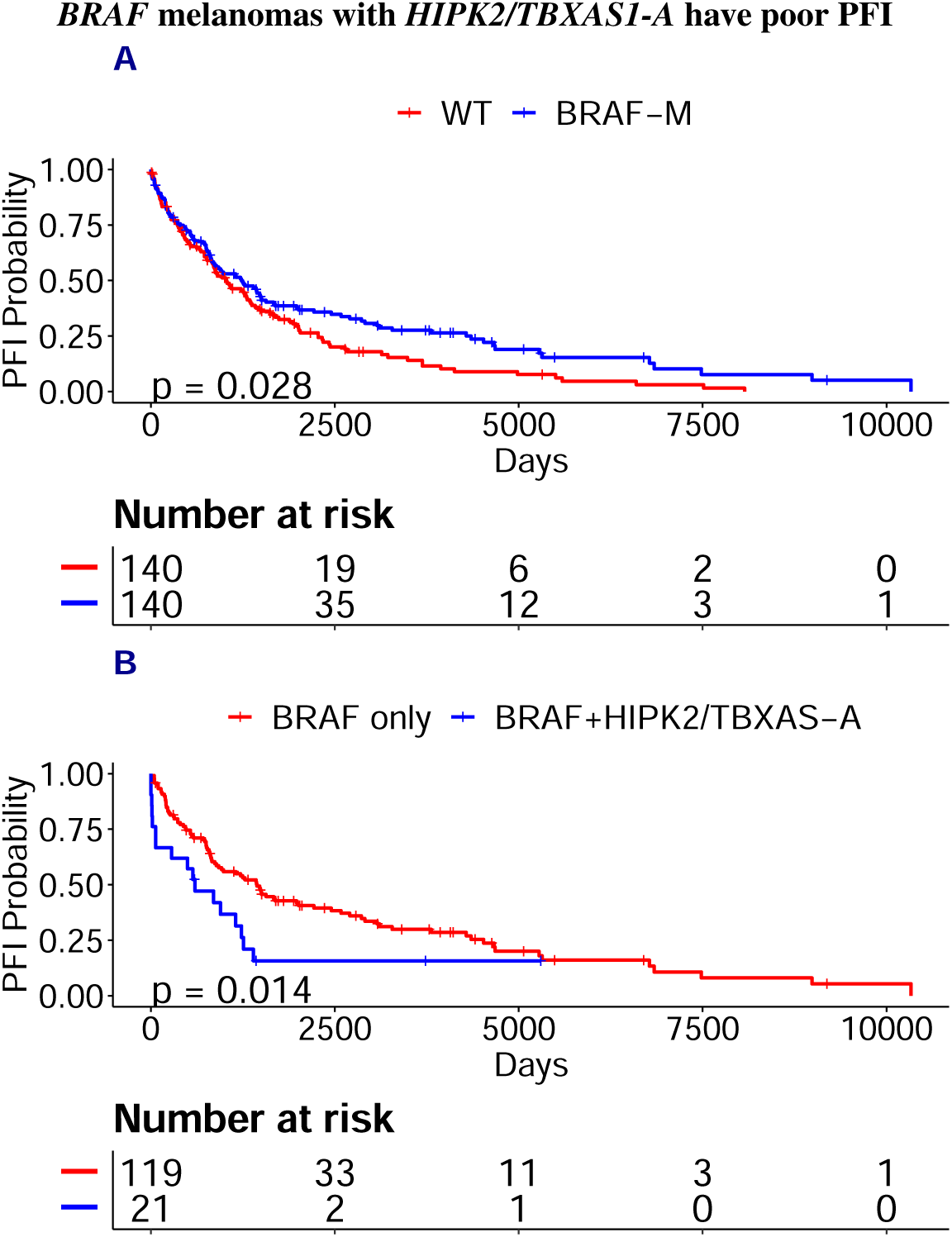
*HIPK2/TBXAS1-A* annotates an amplified region that contains *BRAF* and defines a subtype of *BRAF* mutant SKCM patients with poor PFI. A) *BRAF* patients have improved PFI compared to non-*BRAF* patients. B) SKCM patients with *BRAF*+*HIPK2/TBXAS1-A* have worse PFI than patients with only *BRAF*. P-values calculated from Kaplan–Meier estimator.

The wide amplification peak annotated by *RN7SKP254* is on chromosome 15q26.2 and contains 169 genes. Four genes were classified as cancer genes based on the Sanger Institute Cancer Gene Census [60]: *BLM, IDH2, NTRK2* and *CRTC3*. This rule was prioritized by CRSO as the second highest confidence rule (98%) despite ranking 32nd in coverage (6.6%) and 21st in SJ ranking, suggesting that this amplification may merit further investigation.

### 2.4 Known and novel findings in TCGA

In this section we highlight some findings from the TCGA CRSO results. Table S6 presents all con-GCRs, and provides a concise snapshot of the results from all cancer types.

#### 2.4.1 Known 3-gene combinations in brain and colorectal cancers

One of the distinguishing features of CRSO is the ability to identify functionally relevant cooccurrences involving 3 or more events. The rule *ATRX-M*+*IDH1-M* +*TP53-M* was identified as a con-GCR in both LGG (conf.: 71) and GBM (conf.: 63), despite only occurring in 3.7% of GBM samples. This 3-gene combination defines a well known subtype of brain cancers with differential prognosis, and has been experimentally shown to induce carcinogenesis [61, 62, 63]. Similarly, the well-studied combination of *APC*+*KRAS*+*TP53* in colorectal cancers [64, 65, 66] was identified as a con-GCR in READ (conf.: 79), and as a non-consensus GCR in COAD (conf.: 42).

#### 2.4.2 CRSO prioritized *NFE2L2* combinations in multiple cancers

*NFE2L2* mutations were identified as part of a con-GCR in 4 cancer types: LUSC, HNSC, ESCA and BLCA. All of the con-GCRs involving *NFE2L2* ranked very low in terms of single-rule metrics (Table 3), suggesting that *NFE2L2* is accounting for samples not accounted for by any higher ranking rules. *NFE2L2* encodes NRF2, a key transcription factor that regulates cellular response to oxidative-stress [67]. Initially, NRF2 activation was identified as a mechanism of cellular protection against cancer [68, 69]. However, we now have evidence that constitutive NRF2 activation via mutations in *KEAP1* or recurrent *NFE2L2* exon 2 deletions can drive tumor proliferation and metastasis [70, 67]. *NFE2L2* mutations have recently been shown to define subtypes with differential prognosis in lung and head and neck cancers [71, 72, 73, 74]. Several upcoming clinical are designed to explore the therapeutic benefit of compounds that inhibit NRF2 in advanced cancer patients harboring mutations in *NFE2L2* or *KEAP1* [75, 76, 77]. The rules containing *NFE2L2* may refine our understanding of the contexts in which *NFE2L2* drives cancer and may identify subsets of patients that are responsive to *NFE2L2* inhibition.

#### 2.4.3 CRSO prioritized a rare combination in head and neck cancers (HNSC)

The rule *CASP8-M*+*HRAS-M* was identified in HNSC as the third highest ranking GCR (conf. = 73) even though it ranks 591st in coverage and 301st in SJ out of 926 rules. Patients that have these two mutations and are also wild-type for *TP53* have been shown to define a biologically distinct subtype characterized by low SCNV burden [78]. The prioritization of *CASP8-M*+*HRAS-M* despite low coverage (3.8%) demonstrates CRSO’s capability to systematically identify non-obvious, biologically meaningful combinations among the myriad of possible combinations.

#### 2.4.4 CRSO prioritized rules containing *ALB* mutations in liver cancer

Two high confidence GCRs were identified in LIHC involving *ALB* mutations: *ALB-M*+*CTNNB1-M* (confidence = 98) and *ALB-M*+*TP53-M* (confidence = 80). *ALB* was experimentally shown to be a tumor suppressor in hepatocellular carcinomas [79], and has been discussed as an important part of the somatic mutation landscape [80]. Cooperations involving *ALB* have not been systematically reported, and the two combinations we identified may help inform context-dependent treatments of *ALB* mutant patients. Evidence of the relevance of these rules on PFI is presented in Section 2.5.

#### 2.4.5 CRSO identified a novel 3-gene combination in bladder cancer

*ARID1A-MD*+*SOX4-A*+*TP53-M* was identified as a con-GCR in BLCA (conf. = 79, coverage = 7%). *SOX4* over-expression has been studied experimentally and has been reported to be an important contributor in bladder cancer tumorigenesis [81, 82]. The hypothesis that *SOX4* cooperates with *ARID1A* and *TP53* to initiate bladder carcinogenesis is novel and may merit further experimental investigation.

### 2.5 Associations of rules with patient outcomes

We evaluated whether stratifying patients according to con-GCRs predicted by CRSO instead of individual driver events provided extra prognostic information. We restricted this analysis to events that appear in more than one con-GCR, (referred to as **multi-rule events**), since these events are predicted by CRSO to occur in distinct genetic contexts. For every con-GCR, *R*, containing a multi-rule event, *E*, we performed univariate cox-ph analysis to compare the PFIs of samples that satisfy *R* versus samples that harbor *E* but do not satisfy *R* (Methods 5.5).

A total of 289 tests were performed across the 19 cancer types, and 21 (7.3%) significant associations (| *Z*| ≥ 1.96) were detected (Table 4). Positive *Z*-scores indicates better outcomes in the cohort satisfying *R* versus that only harbors *E*, and negative *Z*-scores indicate the opposite. To account for multiple hypotheses, an adjusted *P* value, *P*_*Adj*_, (Table 4) was calculated using a permutation procedure (Methods 5.5).

#### 2.5.1 CRSO combinations refine the classification of *IDH1*-mutant LGGs

*IDH1* mutations occur in 78% of LGGs and is biomarker for improved outcome in LGG patients [61] (Figure 6A). We detected differential outcomes associated with the event/rule pairings *IDH1-M* vs. *IDH1-M*+*CIC-M* and *IDH1-M* vs. *IDH1-M*+*PIK3CA-M* that could further stratify *IDH1* mutant samples (Figure 6). Among the patients with *IDH1-M* (*n* = 397), those that also have *CIC-M* (*n* = 99) have better PFI than those that are *CIC* wild-type (*n* = 298, Figure 6B, *Z* = 2.5). On the other hand, the 31 *IDH1*-mutant samples that also harbor *PIK3CA-M* have worse PFI than those that are *PIK3CA* wild-type (*n* = 366, Figure 6C, *Z* = −2.1). Collectively, these results suggest that LGG patients could be stratified into 4 groups: *IDH1* wild-type, *IDH1* mutant+*CIC* mutant, *IDH1* mutant+*PIK3CA* mutant, and *IDH1* mutant+*CIC/PIK3CA* wild-type. Figure 6D suggests that *PIK3CA-M* appears to nullify the improvement in PFI conferred by *IDH1*, and that *CIC-M* enhances the improvement in PFI. The improved prognosis of *IDH1-M*+*CIC-M* relative to other *IDH1* mutant LGGs has been previously reported and has been independently characterized by p1/q19 co-deletions, which overlap highly with *CIC* mutations [61, 83, 63]. We did not find literature evidence of the poor prognosis of *IDH1* tumors harboring *PIK3CA* mutations in LGGs

**Figure 6:**
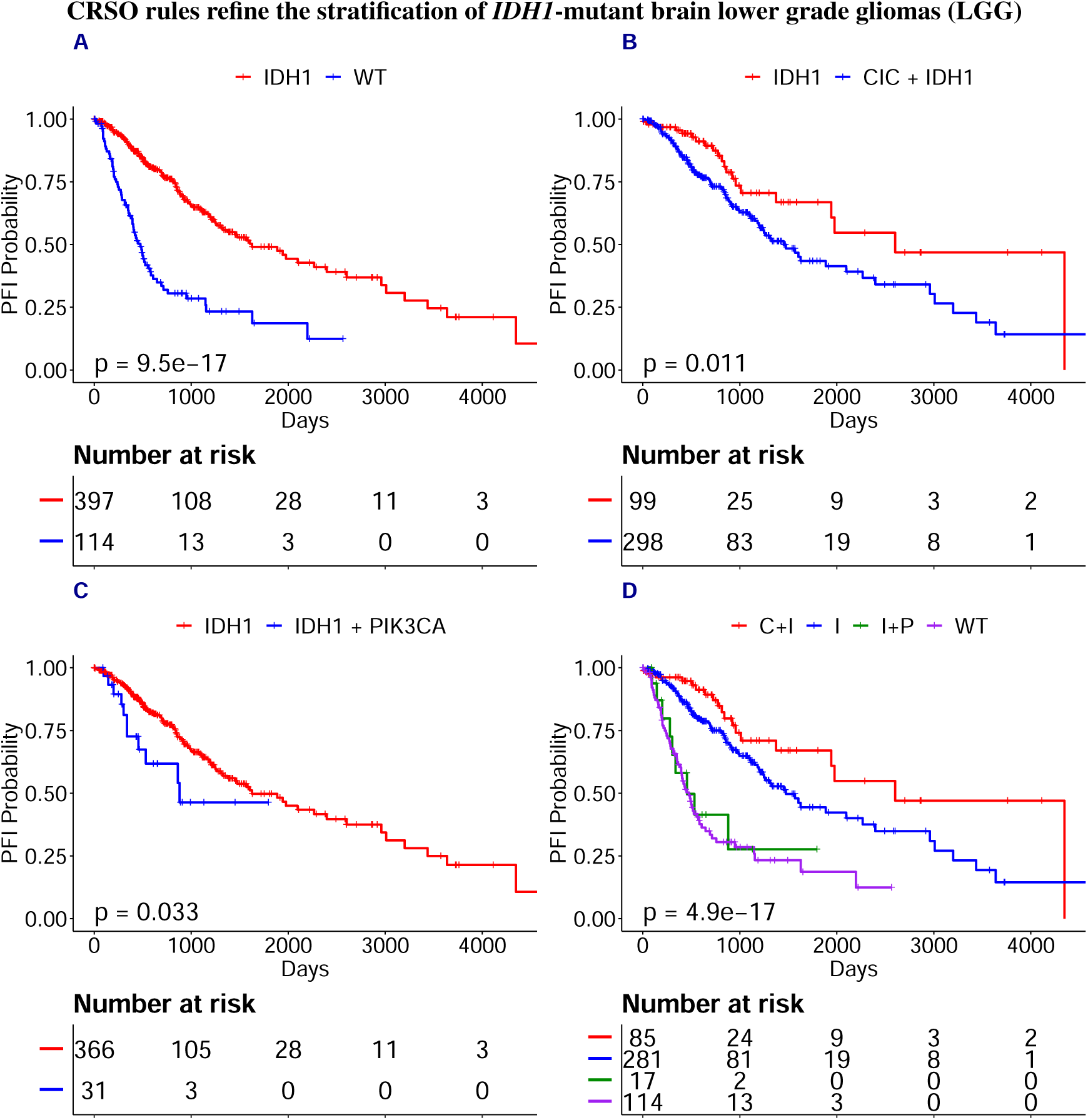
A) *IDH1* mutations define a subset of LGG patients with better PFI. B) Patients with *IDH1* and *CIC* have better PFI than patients with *IDH1* only. C) Patients with *IDH1* and *PIK3CA* have worse PFI than patients with *IDH1* only. D) LGG patients can be stratified into four classes: *IDH1* wild-type (WT), *IDH1* and *CIC* (C+I), *IDH1*+*PIK3CA* (I+P) and *IDH1* wild-type for both *PIK3CA* and *CIC*. Samples harboring *IDH1, PIK3CA* and *CIC* were excluded from panel D (n = 14). P-values calculated from Kaplan–Meier estimator.

#### 2.5.2 CRSO identifies potential multi-gene biomarkers in liver cancer

LIHC patients with both *ARID1A-MD* and *CTNNB1-M* (*n* = 23) had worse PFI expectancy than patients with *ARID1A-MD* but not *CTNNB1-M* (Table 4, *n* = 61, *Z* = −2.89, *P*_*Adj*_ = 0.006). The patients satisfying this rule also had worse PFI expectancy than those with *CTNNB1-M* but not *ARID1A-MD* (*n* = 74, *Z* = −2.27, *P*_*Adj*_ = 0.079). Surprisingly, neither *ARID1A-MD* nor *CTNNB1-M* is a biomarker individually (Fig. 7A-B), and yet together they appear to define a subtype with significantly worse prognosis (Fig. 7C-D).

**Figure 7:**
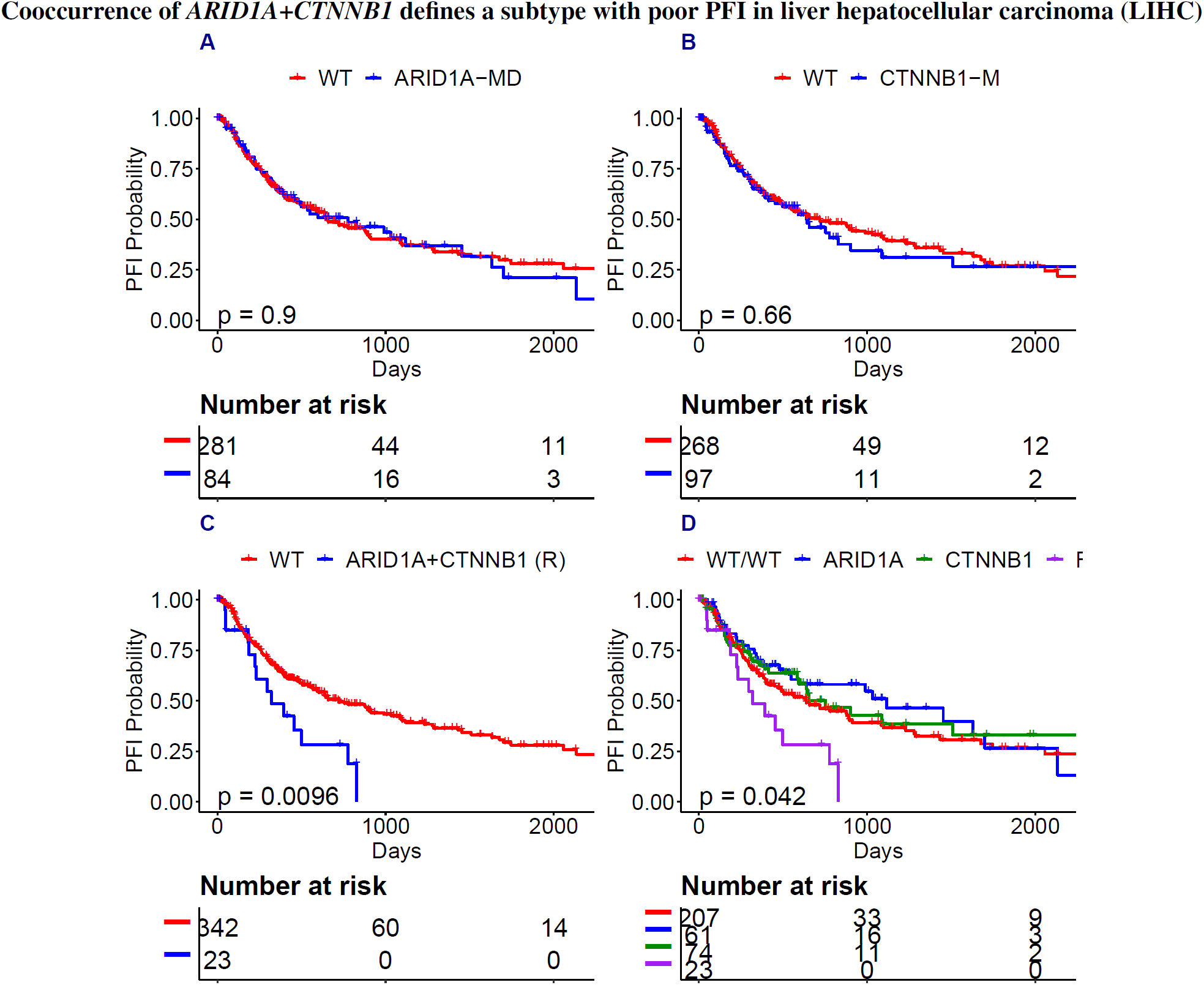
A) *ARID1A-MD* is not a biomarker in LIHC. B) *CTNNB1-M* is not a biomarker. C-D) LIHC patients with *ARID1A-MD* and *CTNNB1-M* define a subtype with significantly worse prognosis compared to all other patients. P-values calculated from Kaplan–Meier estimator.

A second subtype with poor prognosis in LIHC was defined by *ALB-M*+*TP53-M*. Patients harboring both of these events had significantly worse PFI than those harboring either event individually, or neither event (Figure 8).

**Figure 8:**
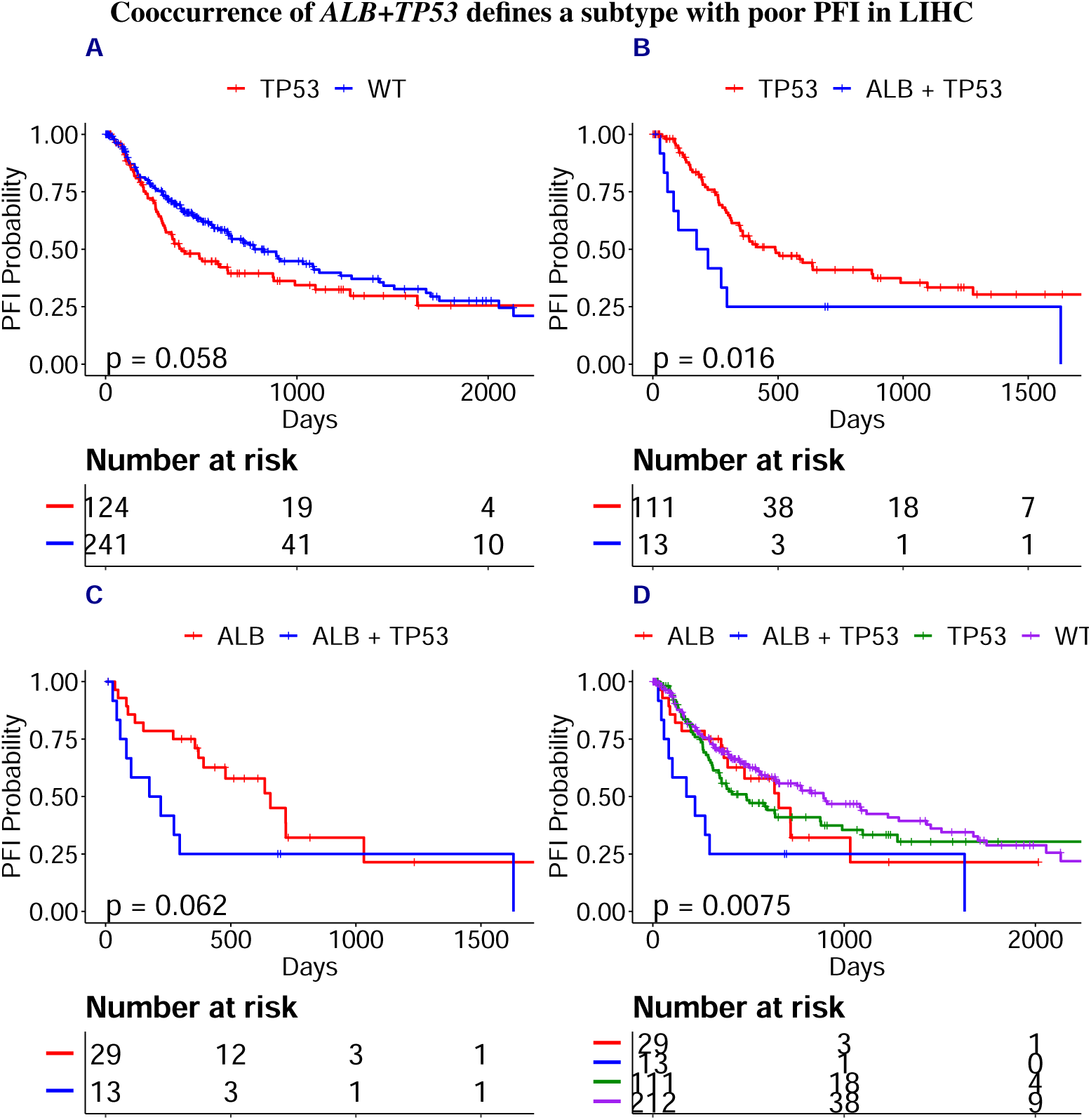
A) *TP53* correlates with worse PFI in LIHC. B-C) *ALB*+*TP53* patients have worse PFI than patients with only one mutation. D) LIHC patients with *ARID1A-MD* and *CTNNB1-M* define a subtype with significantly worse prognosis compared to all other patients. P-values calculated from Kaplan–Meier estimator.

#### 2.5.3 CDKN2A and FGFR3 co-mutation status suggest a 3-tier stratification of BLCA tumors

*FGFR3-MA*+*KDM6A-MD* and *FGFR3-MA*+*CDKN2A-MD* are the 1st and 3rd highest confidence GCRs in BLCA. The association between *CDKN2A* and prognosis is complicated in bladder cancer, as both p16 protein over-expression and complete lack of expression have been shown to be biomarkers of poor prognosis in bladder cancers [84, 85]. In our study, *CDKN2A* deletion and *CDKN2A* mutations were combined into a single hybrid event, resulting in a simple association between *CDKN2A-MD* status and poor PFI (Figure 9A). We found that *FGFR3-MA* confers improved PFI within *CDKN2A* wild-type tumors (Figure 9A), but does not associate with any PFI difference in tumors harboring *CDKN2A-MD* (Figure 9B-C). These results suggest a 3-tier classification, with *CDKN2A-MD* defining a tier with poorer prognosis, *CDKN2A/FGFR3* double wild-type defining a tier with intermediate prognosis, and *FGFR3-MA*+/*CDKN2A-MD*-defining a tier with improved prognosis (Figure 9D).

**Figure 9:**
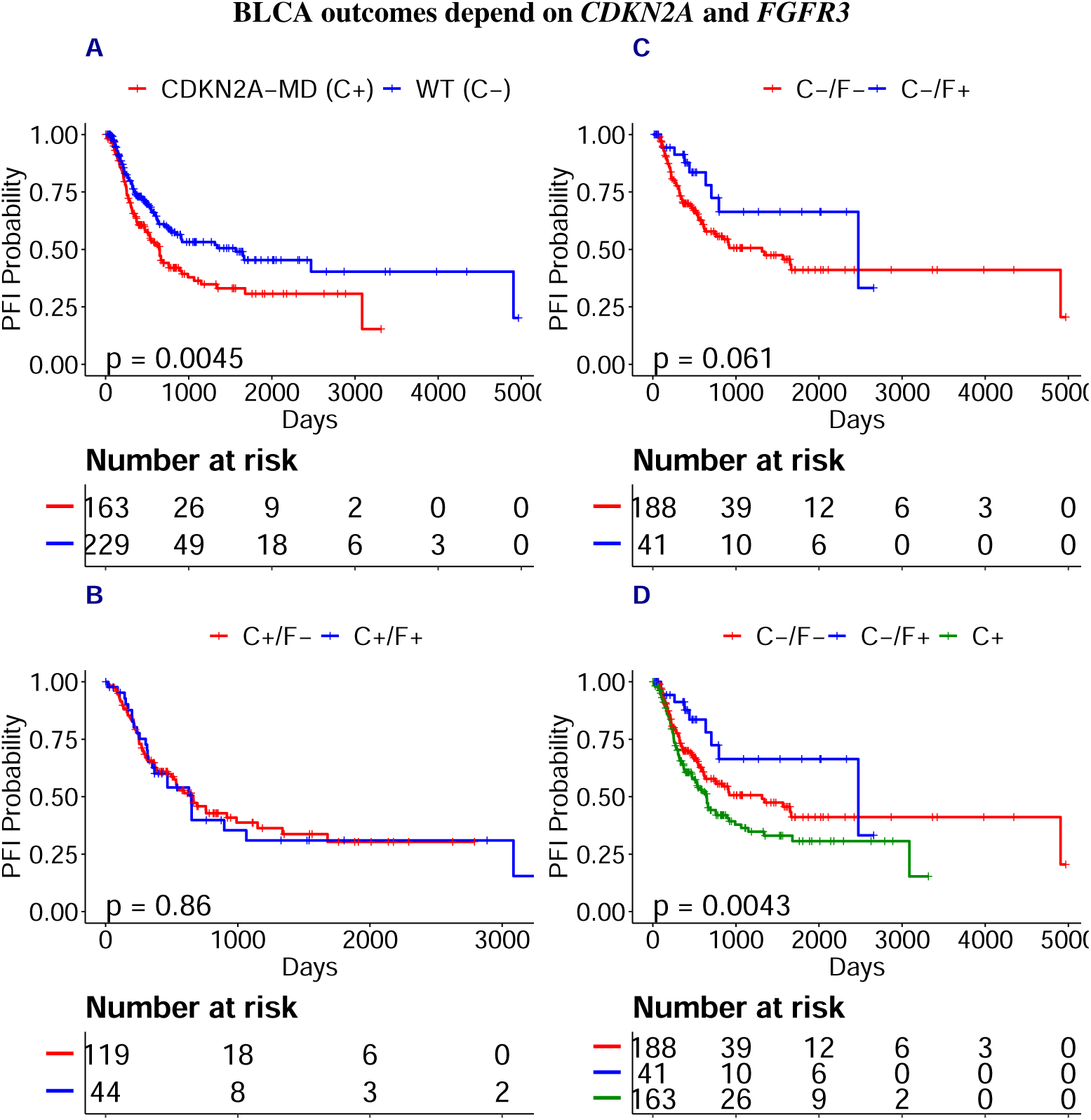
C+ indicates tumors harboring *CDKN2A-MD*, C-indicates tumors wild-type for *CDKN2A*. F+ indicates tumors harboring *FGFR3-MA*, and F-indicates tumors wild-type for *FGFR3* A) *CDKN2A-MD* is a single-gene biomarker for poor PFI in bladder cancer. B) *FGFR3-MA* is a biomarker for improved survival among *CDKN2A* wild-type samples. C) *FGFR3-MA* status does not correlate with PFI in *CDKN2A-MD* tumors. D) Proposed 3 tier stratification of BLCA defined by C+, C-/F- and C-/F+. P-values calculated from Kaplan–Meier estimator.

### 2.6 Comparison with SELECT

We compared the CRSO TCGA results with the pairs of cooccurrences identified by Mina *et al*. using SELECT [24]. Sixteen TCGA cancer types were analyzed by both SELECT and CRSO (Table 5). COAD and READ were excluded from the side-by-side comparison because they were analyzed as a single colorectal cancer type in Mina *et al*.. Pancreatic adenocarcinoma (PAAD) was excluded because it was not analyzed in Mina *et al*.. For each tissue we compared the total coverage of all statistically co-occurrent pairs identified by SELECT within a curated set of 505 pan-cancer mutations and CNVs [24], versus the coverages of the core rule sets identified by CRSO (Table 5). Although both methods identified a similar mean number of synergies per cancer type, (10.4 ± 10.2 for SELECT versus 11.1 ± 2.2 for CRSO), the CRSO core rules covered an average of 68% of samples (sd = 12%) per cancer type compared to an average of 19% of samples (sd = 11%) covered by the SELECT synergistic pairs. The large discrepancy in coverage is because CRSO does not require statistical cooccurrence between the events in a cancer rule. CRSO’s ability to identify a likely driver combination in the majority of samples, and to identify combinations of more than 2 events, may facilitate precision oncology advances that could benefit many patients.

**Table 5:** Coverages of SELECT pairs and CRSO core RS or 16 TCGA cancer types

|  | Select_Cov | CRSO_Core_Cov | Select_Num_Pairs | CRSO_Num_rules |
| --- | --- | --- | --- | --- |
| BLCA | 0.38 | 0.75 | 20 | 14 |
| BRCA | 0.27 | 0.58 | 29 | 11 |
| CESC | 0.05 | 0.53 | 2 | 12 |
| ESCA | 0.29 | 0.84 | 10 | 13 |
| GBM | 0.38 | 0.77 | 3 | 11 |
| HNSC | 0.28 | 0.68 | 9 | 10 |
| KIRC | 0.07 | 0.52 | 2 | 10 |
| LGG | 0.06 | 0.65 | 2 | 5 |
| LIHC | 0.09 | 0.56 | 0 | 12 |
| LUAD | 0.20 | 0.60 | 7 | 10 |
| LUSC | 0.15 | 0.83 | 2 | 14 |
| OV | 0.20 | 0.86 | 32 | 12 |
| PRAD | 0.10 | 0.51 | 7 | 10 |
| SKCM | 0.11 | 0.69 | 9 | 10 |
| STAD | 0.28 | 0.64 | 24 | 13 |
| UCEC | 0.11 | 0.84 | 9 | 11 |
| **Average** | **0.19** | **0.67** | **10.4** | **11.1** |

All SELECT predicted synergies for each cancer type were extracted, and coverage for the cancer type was calculated using the union of covered samples. CRSO Core RS coverages for each cancer type.

For each cancer type, we compared the CRSO con-GCDs (conf. ≥ 50) with the co-occurring pairs detected by SELECT using the subset of mutation events that overlap between the datasets. Copy-number events were excluded because they were processed and defined differently in Mina *et al*. (Methods 5.6). In total, 99 duos were identified by either CRSO or SELECT, 5 duos were identified by both algorithms, 19 duos were identified by SELECT only and 75 duos were identified by CRSO only (Table S7). The large discrepancy in the number of pairs is partially because the common mutation set represents a larger fraction of the CRSO event pool. Additionally, a single CRSO rule can contain 3, 6 or 10 duos if it contains 3, 4 or 5 events. Of the 19 duos identified by SELECT that were not con-GCDs, 13 had coverage below the CRSO rule-inclusion threshold of 3% (6 had coverages ≤1%). Of the 6 duos with coverage ≥ 3% identified by SELECT only, 4 were identified by CRSO but did not achieve 50% confidence threshold, and only 2 were not identified at all by CRSO.

CRSO identified many well-known and high coverage combinations that were missed by SELECT, including *IDH1*+{*ATRX/TP53/CIC/PIK3CA*} in LGG, *BRAF*+{*PTEN/TP53*} in SKCM, and many high coverage combinations involving *PIK3CA* in UCEC. Moreover, the full set of cooccurrences identified by SELECT (i.e., not restricted to common events), included 0 combinations involving *IDH1* in LGG, 0 combinations involving *BRAF* in SKCM, and 1 low-coverage combination involving *NRAS* in SKCM. Many of the CRSO con-GCDs missed by SELECT consisted of a common tumor-suppressor such as *TP53* and *ARID1A* cooperating with a growth-promoting oncogene. Tumor-suppressors that can cooperate with many genes appear to be independent from all of them and are overlooked by approaches that rely on statistical cooccurrence. Supporting this explanation, Mina *et al*. reported that both mutual-exclusivity and cooccurrence are found at higher rates between genes that are within the same pathway [24].

## 3 Discussion

We developed CRSO as a stochastic optimization procedure for predicting combinations of alterations that are minimally sufficient to drive cancer in individual patients. We applied CRSO to 19 TCGA cancer types, using SMGs identified by dNdScv [5] and SCNVs identified by GISTIC2 [6] as input features. An optimal core rule set was determined for each cancer type, as well as sets of generalized core rules, trios and duos, along with confidence scores.

CRSO is an optimization over a computationally intractable search space of rule sets, i.e., sets of combinations of events. We performed extensive simulation analyses with known ground truth rule sets and found that CRSO was able to reproduce the ground truth rule sets with high sensitivity and specificity, and that CRSO was able to assign individual patients to rules with high accuracy. Analysis of TCGA results showed that CRSO identified synergies predicted by a leading algorithm for detecting pairwise synergies, and nominated many additional cooperations. We showed that CRSO prioritized biologically important events and combinations with literature support that would not be prioritized based on single-event analysis or other tools.

CRSO is developed based on a theoretical model that assumes that every tumor results from a specific minimally essential driver combination, and that a small number of such combinations account for the majority of tumors in each cancer type. Ultimately, whether a rule is minimally sufficient cannot be proven by CRSO alone. We also do not presume that CRSO is finding the “ground truth rule set” for the 19 cancer types, nor that an unambiguous “ground truth rule set” exists in all cases. Rather, we hypothesize that some of the rules identified by CRSO are of biological and clinical significance, and that the probability of a rule being of biological significance is positively correlated to its generalized core confidence score.

Several characteristics of CRSO distinguish it from other available methods for detecting biological cooperation from genetic profiles. Most approaches can only detect pairwise synergies, whereas CRSO is able identify combinations of 3 or more events. CRSO takes into account passenger event probabilities by optimizing over a probabilistic representation of genetic alterations in addition to the binary representation used by most approaches. CRSO does not use pairwise statistical significance as a criteria for driver gene cooperation. Instead, CRSO optimizes for coverage of the population and minimization of passenger penalties, using a minimal number of rules.

### 3.1 Novel findings and multi-gene biomarkers from 19 TCGA cancers

We highlighted several novel combinations from among the high confidence rules inferred by CRSO in 19 TCGA cancer types (Table S6).

*NRAS-M*+*HULC-A* and *NRAS-M*+*SMYD3-A* were identified as core rules that may represent distinct mechanisms of carcinogenesis in *NRAS* mutant melanomas 2.3. *HULC* and *SMYD3* amplifications are lesser-known events that have been shown in the past few years to contribute to tumorigenesis in multiple cancer types, but have not been identified as driver events in melanoma. *SMYD3* was shown halt tumor progression in *RAS* driven mouse models of lung and pancreatic cancers [55], and may represent a novel therapeutic target in *NRAS*-mutant melanomas.

CRSO identified *NFE2L2* as con-GCRs in 4 cancer types as part of 4 distinct rules that had comparatively low coverage and single-rule performance (3).The CRSO findings support that *NFE2L2*-mediated NRF2 activation may be an essential driver in multiple cancers types.

In LIHC, CRSO nominated *ALB-M*+*CTNNB1-M* and *ALB-M*+*TP53-MD* as the 2nd and 9th highest confidence GCRs (Table S6), respectively, suggestion that *ALB* mutations may play a vital role in the formation of some liver cancers. *ALB-M*+*TP53-MD* was further identified to be a subtype of LIHC with poor prognosis.

Additional examples were presented in LGG, LIHC and BLCA of significant differences in patient PFIs that were found based on rules identified by CRSO. In all of these examples the PFI differences could not have been identified by consideration of individual alterations. Rather, these differences appear to be a consequence of specific combinations of alterations who’s cooccurrences define subtypes with better or worse prognosis. These combinations, as well as other combinations in Table 4 that were not discussed, merit further investigation as prognostic biomarkers. Evaluating these findings on independent datasets is a necessary next step toward determining whether any of the combinations are reliable biomarkers that can assist clinical stratification. Incorporating information about patient treatments may help refine this analysis. The interpretability of potential biomarkers identified by CRSO may provide actionable information that goes beyond current clinical stratification, such as suggesting combination treatments or identifying specific alterations as drug targets.

### 3.2 Limitations of pairwise cooccurrence tests and comparison with SELECT

We evaluated whether pairwise cooccurrence tests could reproduce the pairwise ground-truth synergies in 190 simulated datasets based CRSO model assumptions and empirically estimated passenger alteration probabilities. We found that pairwise cooccurrence testing resulted in overall poor performance, with low sensitivity and low specificity. Moreover, we found that pairwise tests had poor sensitivity even when all passenger events were excluded (Fig. S5A). The inadequacy of pairwise cooccurrence tests can be explained by the complicated cooperation network of cancer rule sets that allows for individual alterations to be part of multiple different rules. Consider as a minimal illustrative example that a cancer rule set is defined by 3 rules encompassing 3 events: *RS* = {*A* + *B, A* + *C, B* + *C*}. If these three rules each occur in 1/3 of the population and are perfectly mutually exclusive, we would observe perfect statistical independence between the three events using pairwise Fisher-exact tests.

A real data comparison was performed between CRSO and SELECT, a popular, recently developed algorithm [24] for finding pairwise relationships based on a model of conditional selection. CRSO and SELECT were applied to 16 common cancer types. Although both methods use TCGA mutations and SCNVs as inputs, and both methods predict a similar number of pairs/rules per cancer type, the percentage of the cohort harboring at least one CRSO rule is much higher than the percentage harboring both events of at least one SELECT pair (see Table S7). The results filtered through a common mutation set suggest that CRSO is able to capture many of the higher coverage SELECT pairs without screening for cooccurrence.

SELECT uses an information theoretic approach to prioritize event pairs based on the strength of pairwise inter-dependency patterns [24]. Ultimately, SELECT seeks to find pairs of events that co-occur more or less frequently than expected if the events were independent. Our analysis shows that this criterion leads to the omission of many well-known, high coverage oncogenic synergies. Many of the combinations detected by CRSO and missed by SELECT involve tumor suppressors that can cooperate with multiple oncogenes in different tumors. For example, SELECT did not detect any of the well-studied and highly-recurrent combinations in melanoma between *TP53* or *CDKN2A* with either *BRAF* or *NRAS*. CRSO does not assume that pairwise statistical dependence is required for biological synergy and is therefore able to identify driver combinations involving tumor suppressors that cooperate with many oncogenes to contribute to carcinogenesis.

### 3.3 CRSO limitations and future improvements

CRSO is a tool for prioritizing biologically relevant combinations that may merit further investigation from a huge space of possible combinations, and it is inevitable that some identified combinations will be false positives, and some important biological combinations will be missed. In this section we discuss likely sources of false positives and false negatives, and possible steps that can be taken to improve the accuracy of the results.

#### 3.3.1 Inclusion of missing driver events

The coverages achieved by the core rule sets in different cancers ranges from a low of 51% in prostate adenocarcinoma (PRAD), to a high of 89% in rectum adenocarcinoma (READ) cancers, revealing that a substantial subset of patients in every cancer type were not assigned to any rule. One strategy for accounting for these samples would be to expand the set of mutations and SCNVs included as events by relaxing the significance threshold used for dNdScv or GISTIC2, or by including additional events identified by other approaches.

The rules identified by CRSO are further limited by the types of events that are used as inputs. We chose to use SCNVs and SMGs because these are the most common types of driver alterations in cancer, and can be systematically prioritized by statistical enrichment methods. However, many other types of alterations were omitted that may contribute to cancer formation, including germline alterations, arm level CNVs, gene fusions, chromosomal translocations and epigenetic alterations. For examples, *TMPRSS2-ERG* fusions are observed in 40% of prostate cancers [86], and p1/q19 chromosomal co-deletions are observed in 30% of lower-grade gliomas [83]. CRSO supports inclusion of any binary-encoded event of interest. In case it is hard to calculate passenger probabilities for an event of interest, we recommend assigning a penalty equal to 1.05 times the largest penalty in each sample that harbors this event. Doing so would ensure that the event has maximum priority in the samples that harbor it. Using a much larger penalty compared to those associated with SCNVs and mutations could adversely impact the coverage of CRSO by over-prioritizing rules that cover very few samples.

Since it is inevitable that some important drivers are missing from our datasets, we performed an experiment investigating the robustness of the melanoma results to exclusion of important events. The results suggest that most rules identified in the absence of an important driver event will also be identified when the event is included (Section 5.4, Table S5). Nonetheless, it is likely that exclusion of very high frequency driver events, such as *TMPRSS2-ERG* fusion, will lead to identification of some false positives.

#### 3.3.2 Improved variant annotation can refine inputs and improve detection of patient-specific driver combinations

It is sometimes possible to predict the functional consequences of specific alterations within candidate driver events, e.g., using CIViC [87]. Variant annotation can be integrated into CRSO by modifying the inputs so that high confidence neutral alterations are labeled wild-type.

Future implementations of CRSO may consider distinguishing between bi-allelic and mono-allelic mutations. For example it is possible that a combination involving bi-allelic loss of a tumor suppressor is sufficient to drive cancer initiation, whereas mono-allelic loss in the same combination may require additional alterations. Unfortunately, distinguishing between bi-allelic and mono-allelic mutations from bulk tumor samples remains challenging due to intra-tumor heterogeneity [88].

#### 3.3.3 Limitations of CNV representation and passenger probability calculation

CRSO uses the focal amplification and deletion peaks identified by GISTIC2 as input features. Some GISTIC2 peaks contain many genes, making it difficult to identify the genes of biological significance.

For mutations, we were able to account for covariates that impact mutation rates and estimate passenger probabilities rates of individual amino-acid substitutions using CancerEffectSizeR [89]. Unfortunately, analogous tools for calculating passenger probabilities of SCNVs that account for built-in biases do not exist, and we relied on a crude approach to estimate these probabilities based on alterations rates in control cytobands. One strategy towards better prioritizing likely driver SCNVs would be to filter out deletions and amplifications that do not have a corresponding impact on gene-expression dosage in individual patients, similar to the approach use by Mina *et al*. [24].

We observed retrospectively that some rules identified by CRSO contained multiple GISTIC peaks on the same chromosome, with high correlation across patients are likely to be artifacts. One strategy for avoiding this unwanted bias would be to disallow rules involving multiple co-directional copy-number events on the same chromosome, or to implement a distance threshold. Fortunately, only a small fraction of core rules and con-GCRS were in this category, and only no rules that we highlighted in the manuscript belonged in this category.

#### 3.3.4 Refinement of cohorts may improve accuracy

In the present analysis we applied CRSO to the full TCGA cohorts for 19 cancer types. It may be of interest to apply CRSO to previously established subtypes to better understand cancer causation within minor subtypes. For example, the melanoma analysis only revealed 1 rule that was wild-type for both *BRAF* and *NRAS*, and many samples in this subtype were not assigned to any rule.To better understand these samples, users may run CRSO only on the subtype of double-wild-type melanomas. We recommend running dNdScv and GISTIC2 on the smaller cohort to identify subtype-specific candidate drivers. Another interesting example would be to run CRSO separately on the established pam50 subtypes in breast cancer [90]. This could identify driver combination specific to basal-like breast cancers, which have limited treatment options and poor prognosis [91]. It would also be interesting to run CRSO separately on microsatellite stable/unstable subtypes of colorectal cancers.

### 3.4 Comparison of CRSO with evolutionary progression models

Computational advances in recent years have enabled inference of the evolutionary histories of individual cancers [92, 93, 94]. In 2017 Jamal-Hanjani *et al*. showed that it was possible to track the evolutionary histories of non-small cell lung cancers by comparing the exome sequences from multiple regions of individual tumors [95]. The authors were able to distinguish early clonal drivers from late subclonal drivers. A recently published study by Gerstung *et al*. used the ratio of duplicated to non-duplicated mutations within amplified regions to reconstruct the timing of driver events in 2,658 cancers [96]. Their analysis found that the earliest driver events overlapped highly with the most recurrent. The ability to reproduce the timing of mutations represents a major advance in our understanding of cancer evolution and presents opportunities for early cancer detection and customizing treatments.

CRSO differs from these methods by introducing a dichotomization of driver events as essential or non-essential. This dichotomization allows for stratification of cancers into a small number of subgroups, each defined by its own cancer rule. Compared to methods that infer patient-specific progression models, CRSO sacrifices comprehensiveness for simplicity and testability. The CRSO results represent a small collection of testable hypothesis about driver event cooperations that recur in significant subsets of tumors.

There is a large gap between the current state of clinical oncology decision making and the computational advances that allow for reconstruction of evolutionary trajectories. It is unlikely these approaches will translate into novel precision treatments in the short term. CRSO can move the needle forward by transitioning from clinical stratification of patients based on single events towards stratifying patients based on cancer rules. Understanding the biological cooperation of a small number of essential drivers can inform novel treatment directions. Information about evolutionary trajectories of individual tumors may improve the accuracy of CRSO predictions by filtering out alterations with strong evidence of being sub-clonal.

### 3.5 Comparison of CRSO with weighted set covering approaches

Dash *et al*. [97] proposed a method for identifying two-hit combinations that are likely responsible for carcinogenesis. The authors represented the problem as an instance of the weighted set cover problem [98] and showed that the identified set of combinations could discriminate normal and cancer samples with high accuracy. The combinations in Dash *et al*. are similar to the rules in CRSO and the set of combinations identified as the solution to the weighted set cover algorithm is similar to the optimal rule set identified by CRSO. However, there is a major difference in the optimization criteria in each of the methods. In [97] sets of combinations are prioritized based on how well they can discriminate normal and tumor samples in a training set. This choice of criteria assumes that combinations that occur only in tumors are likely to be cooperating drivers. There are many differences in the mutational landscapes of tumors and normal tissue and it is possible that many combinations involving passenger mutations are also likely to be observed almost exclusively in tumors. An updated version extending the approach to 3-hit and some 4-hit combinations was recently published by Al Hajri *et al*. [36]. The updated version implements customized GPU hardware to dramatically reduce computation time. The performance of the multi-hit combinations was comparable to the 2-hit combinations, using the criteria of distinguishing normal from tumor. The authors considered all genes containing any coding mutation, and did not annotate variants into functional categories (e.g., they did not distinguish hotspots from other missense mutations). The authors report that the 2-hit version of the algorithm only identified 9 confirmed cancer genes (according to COSMIC) across 197 combinations, missing well-known drivers such as *CDKN2A, BRAF, PIK3CA, EGFR, ARID1A, CTTNB1* and many others. Specific genes were not discussed in the multi-hit version [36]. In contrast to weighted set covering approaches for identifying multi-combinations that are unique to cancer, CRSO reproduces well-known driver combinations and nominates novel combinations with evidence of being biologically relevant to carcinogenesis.

## 4 Conclusion

It has been almost two decades since the advent of next generation sequencing. Projects such as TCGA have produced high quality datasets containing extensive molecular characterization of tens of thousands of cancer patients. Computational tools have been developed to extract information from these data that has greatly advanced our understanding of the molecular underpinnings of cancer evolution. Despite all of this progress, most targeted treatments are prescribed to target a single genetic alteration. It is still not possible to explain much of the heterogeneity in patient responses to different treatments. We developed CRSO as an approach to infer essential driver combinations that cooccur in many patients. CRSO is highly flexible and easy for researchers to use. The results represent testable hypotheses that are easy to interpret. We hope that CRSO will prove helpful in identifying biologically meaningful and clinically actionable combinations of driver alterations.

From a broader perspective, we envision that wide-spread adoption of CRSO by domain experts will propel a shift in the precision oncology community from single-gene thinking towards multi-gene thinking, and that patients will be classified and treated according to driver combinations. For example, we suggest that development of a cancer combination census, analogous to the Sanger Institute Cancer Gene Census [60], would accelerate therapeutic advances by nominating novel therapeutic strategies and refining clinical trial recruitment based on driver combinations. This census would be comprehensive, and include tiers based on the strength of evidence of specific combinations in specific cancer types. For example, one tier would consist experimentally validated driver combinations, such as *BRAF*+*PTEN* in melanomas [28, 34, 29] and *ATRX*+*IDH1*+*TP53* in gliomas [61, 62, 63], and lower tiers would consist of computationally predicted combinations nominated by methods such as CRSO and SELECT, with varying amount of literature support.

## 5 Methods

### 5.1 Representation of TCGA inputs

TCGA data from 19 cancer types (Table 1) were obtained from the January 28, 2016 GDAC Firehose [99]. Candidate driver mutations were defined to be the set of significantly mutated genes (SMGs) identified by dNdScv as significantly mutated using the threshold qsuball < 0.1—using tissue-specific mutational covariates developed within Cannataro *et al*. [89]. Candidate copy number variations were defined to be the set of genomic regions identified by GISTIC2 as amplified or deleted, using the threshold q-residual < 0.25.

The inputs into CRSO are two event-by-sample matrices: **D** and **P. D** is a binary alteration matrix, such that *D*_*ij*_ = 1 if event *i* occurs in sample *j*, and 0 otherwise. **P** is a continuous valued penalty matrix, where *P*_*ij*_ is the negative log of the probability of event *i* occurring in sample *j* by chance, i.e., as a passenger event. In order to calculate passenger probabilities, TCGA mutations and copy number variations were first represented as a categorical event-by-sample matrix, **M**. *M*_*ij*_ can take one of several values called **observation types**. The possible observation types for event *i* differ according to the **event type** of event *i*. Three primary event types were considered: mutations, amplifications and deletions. Mutations were represented at the gene level, whereas both copy number types were represented at the region level as defined by the GISTIC2 narrow peaks. Mutation events take values in the set {*Z, HS, L, S, I*}, corresponding to wild-type, hotspot mutation, loss mutation, splice site mutation or in-frame indel. Amplification events take values in {*Z, WA, SA*}, corresponding to wild-type, weak amplification and strong amplification. Similarly, deletion events take values in {*Z, WD, SD*}, corresponding to wild-type, weak deletion (hemizygous) and strong deletion (homozygous). Two additional event types, referred to as hybrid mutDels and mutAmps, were defined to represent genes that were identified by both dNdScv and GISTIC2 in the same tumor type. When an amplification peak contained an SMG, the amplification and mutation events were consolidated into a single mutAmp event, taking values in the set of mutational observation types, amplification observation types as well as combined observation types for when both an amplification and mutation of the gene were observed in the same sample. MutDels were defined analogously (Section S3).

The entries of **D** only depend on whether an observation is wild-type or not. That is, *D*_*ij*_ equals 0 if *M*_*ij*_ is wild-type, and equals 1 otherwise. By contrast, the entry *P*_*ij*_ is highly sensitive to specific observation type of event *i* that is observed in sample *j*. For example, suppose *TP53* is identified as an SMG within a cancer population. Some tumors may contain one of many nonsense point mutations within *TP53*, whereas other tumors may contain a highly recurrent missense mutation, or a splice site mutation that produces an alternative isoform of the TP53 protein. Although all of these alterations are *TP53* mutations, they occur with very different passenger probabilities, sometimes spanning multiple orders of magnitude. The dual representation defined by **D** and **P** reflects the assumption that different types of alterations within the same event are functionally similar but probabilistically distinct. This representation also allows CRSO to account for sample-specific differences in passenger probabilities, which can sometimes be very substantial. Wild-type events are defined to have passenger probability of 1, so that if *M*_*ij*_ = *Z* then *P*_*ij*_ = 0. Figure 1A shows an example of the input matrices using a miniature dataset that was extracted from TCGA melanoma (SKCM) data for the purposes of illustration.

Exact definitions of observation types and calculation of passenger probabilities are provided in the Supplemental Information, Sections S1 for mutations, S2 for CNVs and S3 for hybrid events.

### 5.2 CRSO Algorithm

CRSO is an optimization procedure over the space of possible rule sets. The ability of a rule set to account for the distribution of events in the population is quantified by an **objective function**. Consider a rule set *RS* = (*r*_1_, …, *r*_*k*_), and suppose each sample has been assigned to one rule in *RS*, or to the null rule. The null rule is implicitly included in every rule set as an assignment placeholder for samples that do not satisfy any of the rules in *RS*. When a sample is assigned to a rule, the events that comprise the rule are considered to be drivers within that sample, and all of the remaining events in that sample are considered passengers. The **statistical penalty** of *RS* is the sum of the penalties of all of the events designated as passengers under *RS*. A penalty matrix, **P^RS^**, is derived by modifying the full penalty matrix, **P**, such that the penalties of all assigned events are changed to 0. For example, suppose sample *j* is assigned to a rule containing events *x* and *y*. This is represented in **P^RS^** by assigning 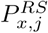 and 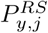 to be 0, instead of the original values they took in **P**.

The objective function score for *RS, J* (*RS*), is defined to be the reduction in total statistical penalty under *RS* compared to the null rule set: 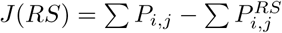. In cases when a sample satisfies multiple rules in *RS*, the sample is always assigned to the rule that maximizes *J* (*RS*), i.e., the rule with largest cumulative penalty in the sample.

#### 5.2.1 Identification of best RS of size K

Given **D**, a starting rule library is built by identifying all rules that contain at least 2 events and occur in a minimum percentage of samples. Except where otherwise indicated, a minimum rule coverage threshold was chosen to be the larger of 3% of the population, or the minimum threshold that at most 2000 rules satisfy. CRSO uses a **four phase procedure** (Fig. 1C) to find the best scoring ruleset of fixed size, *K*, for *K* ∈ {1 … 40}. Since there will generally be a large number of rules in the starting rule library it is impossible to exhaustively evaluate every possible rule set, even for small *K*. The computational time of exhaustive evaluation grows exponentially with the size of the rule pool, i.e., *O*(2^*n*^) for *n* rules. To address this computational limitation, phase 1 is uses a random sampling procedure to prioritize rules according to how likely they are to be among the best rule sets. In phase 2, a subset of the top-performing rules determined from phase 1 are exhaustively evaluated to identify the best rule set of size *K*, for *K* ∈ {1 … 10}. The number of rules that are exhaustively evaluated is subject to a computational constraint on the number of total rule-sets that are considered for each *K*. In general a larger number of rules can be exhaustively evaluated only for very small *K*. The number of rules that can be exhaustively considered is generally small for *k* > 4, meaning that only the top 20-30 rules have the opportunity to be among the best rule sets. Since this subset may be too restrictive, the algorithm is exposed to additional rules in phase 3 by considering many rulesets that include a small number (e.g., 1-3) of unexplored rules mixed with rules from the current best rule set. In phase 4, best rulesets of sizes 11-40 are determined by sampling many rule sets of size *K* + 1 that overlap highly with the best scoring rule set of size *K*. The full description of the four phase procedure is presented in Section S4.

#### 5.2.2 Determination of core RS and generalized core rules

In general larger rule sets perform better than smaller rule sets because the samples in **D** have more assignment opportunities. On the other hand larger rule sets may be more likely to reflect noise in the dataset rather than true biological signal, i.e., over-fitting. Fixing the size of the rule set enforces competition between rules based on the number of events covered per sample, the passenger rate of events covered and the number of samples covered. To mitigate the impact of over-fitting, CRSO requires that every rule in a rule set is assigned to a minimum number of samples, denoted as the *msa* parameter, for *minimum samples assigned*. The default *msa* parameter is 3% of the population. Rule sets that do not satisfy the *msa* threshold are automatically assigned an objective function score and coverage of 0, i.e., they are discarded.

The four phase procedure results in a list of best rule sets of size *K*, for *K* ∈ {1 … *K*_*max*_}, where *K*_*max*_ is the largest *K* for which a valid rule set exists and satisfies the *msa* threshold. Typically *K*_*max*_ is much lower than 40, as large rule sets are constrained by the *msa* requirement as well as the requirement that rule sets do not contain family members, i.e., are valid. The goal of CRSO is to find the rule set that achieves the best balance of objective function score, sample coverage and rule set size, which is called the **core rule set**. The core rule set is defined to be the smallest of the best rule sets of size *K* that achieves 90% of the maximum coverage and performance. Using an arbitrary threshold is a heuristic for automatically choosing a core rule set. In some cases, closer inspection of the results and the convergence curves of performance and coverage can help identify a better choice if a clear plateau is observed. To evaluate the stability of the core rule set and automatically identify a more robust set of rules, 100 iterations of a subsampling procedure are used to identify **generalized core rules** (GCRs).

In each iteration, between 67-85% of the samples are randomly chosen, and a core rule set is determined by evaluating the top 100 phase 4 rule sets for each *K*. The *msa* is scaled according to the subset size, and the core coverage and performance thresholds are randomly sampled uniformly between 85-99%. The full set of GCRs consists of all rules that appear in any of the subsampled cores. After performing 100 iterations, a confidence score is determined for each GCR to be the frequency of inclusion in the sub-core iterations. The subset of GCRs that have confidence levels above 50 by definition cannot contain family members, and are a valid rule set. We refer to this rule set as the **consensus GCR** (con-GCR). The consensus GCR represents a robust estimate of the highest confidence rules for a given dataset that are most likely to reflect true biological cooperation.

In some cases there can be a lack of high confidence GCRs, typically occurring when the core rule set contains many rules with many events. In such a situation, there may be sets of 2 events (duos) or 3 events (trios) that recur often within the sub-cores of each iteration. We calculate **generalized core duos** (GCDs) and **generalized core trios** (GCTs), to better identify recurrent pairwise and three-way synergies. **Generalized core events** (GCEs) are also calculated and represent the frequency of individual events being part of sub-sampled core rule sets.

### 5.3 Phase 1 performance analysis and parameter optimization

We evaluated the performance of phase 1 on the simulations in order to assess whether alternative parameter choices would improve CRSO’s overall performance. Phase 1 prioritizes rules based on their contribution to random rule sets, and is critical for reducing the search space of possible rule sets, which would otherwise be computationally intractable. This is an inherently heuristic strategy whose performance is mathematically difficult to evaluate without knowledge of the ground truth rule set. A phase 1 score was defined for each of the 190 simulations to quantify the similarity of the phase 1 ranking to the maximum possible ranking, corresponding to all true rules in the top *ntr* positions (Section S6.2.1). The mean phase 1 score was more than 0.94 for all ground truth rule set sizes, indicating that phase 1 rankings successfully prioritized the ground truth rules (Table S4). Phase 1 rankings were far superior at prioritizing ground truth rules compared to coverage or SJ rankings for all *ntr*. In 186 out of 190 (0.98) total simulations, 100% of ground truth rules were identified within the top 40 rules post phase 1. By comparison, this was the case for 30 simulations (16%) using coverage rankings and for 19 simulations (10%) using single rule performance (SJ) rankings. On average, 99.8% of true rules were identified among the top 40 phase 1 rules, compare to 64% for coverage rankings and 58% for SJ ranking (see Table S4 for breakdown by *ntr*).

The simulation results suggest that the phase 1 procedure is very effective at prioritizing ground truth rules using the default parameters of *p*1.*ntpr* = 20 and *p*1.*cs* = 0.25. In order to understand how the parameter choices impact the phase 1 performance, we performed phase 1 over a grid of parameter values for each of the 190 simulations. The results showed that phase 1 performance was robust to parameter choices, and that optimal performance can be expected as long as *p*1.*ntpr* ≥ 10 and *p*1.*cs* ≤ 0.5 (Section S6.2.2).

### 5.4 Robustness of CRSO to missing features

The rules that can be identified by CRSO depend on the collection of events included in the inputs. In order to test the robustness of CRSO to the exclusion of important features, CRSO was applied to TCGA melanoma data excluding a single event, for each of the top 15 events. The results were compared to the results from the full dataset. Overall the results suggest that if a driver event is excluded from the inputs, the duos/rules that do not contain the event will generally be comprehensively captured in the incomplete dataset. The rate of false associations caused by the event exclusion is generally small for most events, but can be large if the excluded event is frequent and part of many duos/rules in the full dataset (Supplemental S7, Table S5).

### 5.5 Clinical outcome analysis

Consider multi-rule event *E* that appears in rule *R*. The following test was performed to determine if there is statistical evidence of differential patient prognosis associated with *R*. The subset of patients that contain *E* are considered, and those that are wild-type for *E* are disregarded. Patients within this cohort were assigned to one of two classes: those that satisfy *R* are assigned to class “rule”, and those that harbor *E* but do not satisfy *R* are assigned to class “event”. Univariate cox-ph analysis was performed to test for differences in progression-free intervals (PFI) between the two classes. The result of each test is summarized with a *Z* score, as recommended in [100]. Compared to using *P* values for ascertaining statistical significance, *Z* scores provide additional information about the direction of the PFI differences. Positive *Z* scores indicate better PFI for patients in the rule class compared to patients in the event class, and negative *Z* scores indicate the opposite. PFI data were obtained from a recently published resource for TCGA outcome analysis [101], and the use of PFI as primary endpoint is consistent with the authors’ recommendations for best practices.

For each cancer type, each multi-rule event, *E*, was tested against all of the eligible GCRs that include *E*. The reasons for testing *E* against each rule separately, instead of directly comparing all of the rules containing *E*, are two-fold. First, some multi-rule events occur in many GCRs making it difficult to interpret the results. Second, comparing multiple rules at once introduces a problem of dealing with patients that satisfy multiple rules.

A total of 289 tests were performed across the 19 cancer types, and 21 (7.3%) significant associations (| *Z* | ≥ 1.96) were detected (Table 4). Only those pairings for which both the event and rule classes contained at least 10 patients were considered. It is tempting to argue that the expected percentage of significant associates should be 5% and that we are finding more associations than expected by chance. However, the tests should not be assumed to be independent, and conventional multiple hypothesis corrections may not be appropriate as a consequence. Instead, a permutation test was performed in order to address the multiple hypotheses associated with each multi-event rule. For each multi-rule event, the PFIs of the samples that contain the event were scrambled 1000 times, and the smallest *P* value attained by any of the rule vs. event comparisons was stored for each iteration. An adjusted *P* value, *P*_*Adj*_, (Table 4), was defined to be the fraction of iterations that have permuted *P* value smaller than the real data *P* value.

### 5.6 Criteria for common-event comparison with SELECT

For each cancer type a set of common mutations was identified as the intersection of the SELECT pan cancer mutations with the cancer specific mutation events identified by dNdScv that were used in the CRSO analysis. *CDKN2A* mutations were excluded from the common set of mutations in cancers for which *CDKN2A* was identified as hybrid mutation/deletion in the CRSO analysis. In these cancers, large coverage discrepancies were observed in duos involving *CDKN2A* because SELECT encoded *CDKN2A* mutations as separate events from *CDKN2A* copy number loss. Common event duos for each cancer were defined to be the union of GCDs identified by CRSO and the co-occurrent pairs identified by SELECT for which both events were in the common set of mutations.

### 5.7 Software Details

CRSO is written in R utilizing “ggplot2”, “survival”, “survminer”, “foreach” and “doMPI” packages [102, 103, 104, 105, 106]. We used cancereffectsizeR [89] to calculate amino acid level mutational rates at individual amino acid positions (see Section S1). The CRSO R code is available at https://github.com/mikekleinsgit/CRSO/.

## 6 Frequently Used Acronyms

- CRSO Cancer Rule Set Optimization
- TCGA The Cancer Genome Atlas
- SCNV Somatic Copy Number Variation
- SMG Significantly Mutated Gene
- dNdScv Name of method for identifying significantly mutated genes
- GISTIC2 Name of method for identifying significant copy number variations
- GCR Generalized Core Rule
- GCT Generalized Core Trio
- GCD Generalized Core Duo
- GCE Generalized Core Event
- GC Generalized Core (sometimes used in other contexts such as GC iterations)
- con-GCR Consensus Generalized Core Rule, i.e., GCRs with confidence > 50%
- RS Rule set
- MSA Minimum samples assigned. All rules in a rule set are required to be assigned to at least *msa* samples.
- NTR Number of true rules. Used to describe ground truth rule set size of simulation.
- PFI Progression-Free Interval

TCGA cancer type acronyms referenced in the manuscript:

- SKCM Skin cutaneous melanoma
- LIHC Liver hepatocellular carcinoma
- COAD Colon adenocarcinoma
- READ Rectum adenocarcinoma
- LGG Low grade glioma
- HNSC Head and neck squamous cell carcinoma

## Supporting information

CRSO reports for 19 TCGA cancer types

## 9 CRSO Availability

The CRSO R package is freely available for download at https://github.com/mikekleinsgit/CRSO/.

## 10 Acknowledgements

The results published here are based on data generated by the TCGA Research Network: http://cancergenome.nih.gov/ [107]. The authors thank Dr. Michael Kane for his help with developing the CRSO software package. The authors thank Dr. Christos Hatzis for his counsel.

## 11 Author Contributions

MK, DS and HZ conceived of the project. MK programmed the CRSO software and drafted the manuscript. VC and JT provided SNV mutation rates and dNdScv values that were used to calculate TCGA passenger probabilities. SN reviewed the TCGA results and identified interesting and potentially novel combinations via literature exploration. JT, SN, DS and HZ edited the manuscript. All authors read and approved the final manuscript.

