## Supplementary Materials for "Identifying Modules of Cooperating Cancer Drivers"

#### Contents

**S1 TCGA Mutational Observation Types** 3

**S2 TCGA Copy Number Observation Types** 4

**S3 TCGA Hybrid Events** 5

**S4 Four Phase Algorithm** 5

**S5 CRSO Report Overview from 19 TCGA Cancers** 6

**S6 Simulation Methods** 6

### Supplemental Materials and Methods

#### S1 TCGA Mutational Observation Types

The MAF files for each TCGA dataset are annotated with many different mutation types (Table S1). To account for the fact that different kinds of mutations occur at different baseline probabilities, mutations were subdivided into four observation types: hotspots (HS), loss mutations (L), splicing mutations (S) and in-frame insertions and deletions (I).

**Hotspot mutations** A hotspot mutation was defined to be any SNP that leads to an alteration at a specific amino acid position that is observed in at least three samples within the population. Silent mutations and intronic mutations do not lead to amino acid changes and by definition cannot be hotspots. Most hotspot mutations are missense mutations, but the definition allows for other recurrent SNPs, such as splice site mutation or nonsense mutations to be hotspots as well. Note that the definition of hotspot does not require three instances of the exact same substitution, but rather three instances of substitutions at the same amino acid position. This choice is motivated by the fact that multiple amino acid changes in known hotspots such BRAFV600 and NRASQ61 are observed.

**Loss mutations** A loss mutation was defined as one occurring in a given gene if any mutation is detected except for those mutations that are silent, intronic, splice site, hotspot, in-frame insertions or in-frame deletions. The definition of loss mutations includes missense mutations, nonsense mutations, frame-shift indels and the other rarely observed mutations types shown in Table S1. All of these mutation types were combined under the general category of loss mutations because the majority of non-recurrent mutations will lead to loss of function.

**In-frame Indels** In-frame indels are mutations that are in-frame deletions or in-frame insertions. In-frame insertions/deletions were categorized separately from frame-shift indels because in-frame indels have been shown to be much more likely than frame-shift indels to produce gain of function alterations [1].

**Splice site mutations** The majority of splice site mutations present as point mutations, and they are the fourth most common class of point mutation behind missense, silent and nonsense mutations (Table S1). Splice site mutations can present as point mutations within exons that lead to exon-exclusion as well as point mutations within introns that lead to intron inclusion. Because of this, many splice site mutations are not annotated with a specific amino acid change. Splice site mutations encompass only those splice sites that are non-recurrent single amino acid substitutions. When a splice site mutation occurs as a SNP at a recurrent amino acid position it was designated as a hotspot mutation because this designation permits more accurate calculation of the associated passenger probabilities.

##### Passenger Probability Calculation for Mutations

Passenger probabilities were calculated for every observed mutation. These passenger probabilities are patient-specific, gene-specific and observation-type specific. Mutation rates for every possible amino acid substitution in each SMG were calculated as described in Cannataro *et al.* [2]. These rates are calculated by incorporating both gene-level estimates of mutation rate [3] and tumor-type-specific mutational processes that affect nucleotide substitution rates [4].

Hotspot and loss mutation passenger probabilities were calculated based on the mutation rates of individual amino acid substitutions calculated directly from the *cancereffectsizeR* R package [2]. Hotspot mutation probabilities were calculated for each gene as the sum of the rates of all possible amino acid substitutions at hotspot positions. The loss mutation probability for a gene was calculated as the sum of the rates of all possible amino acid substitutions, except for those that occur at hotspot positions. Non-recurrent splice site substitutions were not excluded because the analysis did not include annotation of all possible splice site amino acid positions. The impact of excluding these sites will be minor, since the number of amino acids per protein is much larger than the number of splice junctions.

Control genes were used to calculate the in-frame indel and splice site probabilities. Control genes were defined to be all genes that are expressed above RSEM = 0 (RNAseq by Expectation Maximization) in at least 5% of samples and were not identified by dNdScv as SMGs. Population level in-frame indel mutation probabilities were calculated to be the frequency of in-frame indel mutations per control gene per sample. Population-level splice site mutation rates were calculated to be the average number of splice site mutations per sample per control gene. The frequencies of in-frame indels and splice site mutations were assumed to be proportional to the number of amino acids in the protein product of each gene. A gene length adjustment factor was defined for each gene to be the number of amino acids in the gene protein product divided by 480 (approximate mean number of amino acids per protein). Gene-specific probabilities for both in-frame indels and splice site mutations were calculated to be the respective population level probabilities multiplied by the gene length adjustment factor.

Mutation frequencies can vary greatly across patients within the same cancer type. To account for this, a patient adjustment factor was used that is based on the number of point mutations in each patient. Mutation counts for each patient were determined to be the total number of point mutations observed outside of the SMGs identified by dNdScv. Patient's with 0 mutations were assigned a mutation count of 1. In general we do not want to remove outliers with large mutation counts because we want the penalties for observations in these samples to be down weighted accordingly. However, there were a few cases where one or two patients had such extreme outliers that they had mutation counts more than 100 times larger than the 90th percentile for the cohort. To mitigate the impact of these extreme outliers, a maximum mutation count was chosen to be the 10 times the 75th percentile of all mutation counts. Patients with mutation counts above the maximum were assigned the maximum mutation counts. Patient adjustment factors were defined to be the patient's mutation count divided by the mean mutation count across the population. The patient-specific probabilities for every mutational observation are the product of the population-level probabilities and the patient adjustment factors.

#### S2 TCGA Copy Number Observation Types

The outputs of GISTIC2 are a set of significantly amplified copy number regions and a set of significantly deleted copy number regions. Each of the significant amplifications/deletions were represented as a single event. To do so, the copy number results were first represented at the gene level by a discrete gene-by-sample matrix of focal copy number status, $\mathbf{M_G}$, and then the scores of individual genes within each region were combined to obtain event level features. The entries of $\mathbf{M_G}$ take values in {SD,WD,Z,WA,SA}, corresponding respectively to strong deletions (SD), weak deletions (WD), wild type (Z), weak amplifications (WA) and strong amplifications (SA). $\mathbf{M_G}$ was constructed by thresholding the continuous value matrix from "focal_data_by_genes.txt". Focal copy number values in $[-0.3, 0.3]$ were designated as copy neutral, as per the noise threshold recommendation in the GDC CNV pipeline [5]. Values above 0.3 were designated as amplifications and values below -0.3 were designated as deletions. To designate copy number alterations as strong or weak the sample specific thresholds provided in the file "sample_cutoffs.txt" were used.

The peak genes for each amplification/deletion were extracted from the tables in the files "table_amp.conf_99"/"table_del.conf_99". For each amplification peak, each sample was assigned to the maximum copy number value attained by any of the peak genes within that sample (i.e., the extreme method). Because the amplification peaks were selected for having evidence of significant amplification, amplification events are only allowed to take values in {Z,WA,SA}. If a deletion is observed within an amplification event it is assigned to be wild type. This procedure results in a discrete matrix of amplification peaks by samples, $\mathbf{M_{AMP}}$, that takes values in {Z,WA,SA}. Each row in $\mathbf{M_{AMP}}$ corresponds to an amplification event identified by GISTIC2. A deletion event matrix, $\mathbf{M_{DEL}}$, was prepared analogously. Deletion peaks were assigned to the minimum copy number value attained by any of the genes in the peak. Amplifications observed within deletion peaks were assigned to be wild type, so that $\mathbf{M_{DEL}}$ takes values in {SD,WD,Z}.

###### Passenger Probability Calculation for Copy Number Events

It is difficult to estimate copy number probabilities at the gene level because of the strong dependence between genes that are near each other. Copy number passenger probabilities were instead estimated at the cytoband level, using control cytobands as the basis for estimating probabilities. Control cytobands were defined to be all cytobands that do not contain any genes that are within any of the significant wide regions reported in "table_amp.conf_99" and "table_del.conf_99".

A control cytoband matrix, $\mathbf{M_C}$, was constructed by assigning each cytoband to the mode of the cytoband's genes observed in each sample from $\mathbf{M_G}$. Consider $\mathbf{M_C}$ to be an $n$ x $m$ matrix, and consider $C_{SD}, C_{WD}, C_{WA}$ and $C_{SA}$ to be the counts of each observation type observed in the population. A population rate for each observation type was defined as follows:

$$\mu_{SD} = C_{SD}/(n * m)$$
$$\mu_{SA} = C_{SA}/(n * m)$$
$$\mu_{WD} = (C_{SD} + C_{WD})/(n * m)$$
$$\mu_{WA} = (C_{SA} + C_{WA})/(n * m)$$

The probabilities for WA and WD were defined to be the frequency of observing any amplification or deletion in order to ensure that $\mu_{WD}$ and $\mu_{WA}$ are always larger than $\mu_{SD}$ and $\mu_{SA}$, respectively.

To account for variation in copy number rates between patients patient-specific adjustment factors were introduced for amplifications and deletions. Let $C_j^{AMP}$ and $C_j^{DEL}$denote the number of control cytobands amplified and deleted in patient $j$, respectively. The amplification and deletion adjustment factors for each patient were defined to be:

$$f_j^{AMP} = (C_j^{AMP} + 0.5)/(mean(C^{AMP}) + 0.5)$$
$$f_j^{DEL} = (C_j^{DEL} + 0.5)/(mean(C^{DEL}) + 0.5)$$

The addition of 0.5 to the counts of all patients ensures non-zero probabilities. The sample specific probabilities for SD and WD were respectively given by: $\mu_{SD,j} = \mu_{SD} * f_j^{DEL}$ and $\mu_{WD,j} = \mu_{WD} * f_j^{DEL}$. The sample specific probabilities for SA and WA were calculated analogously.

#### S3 TCGA Hybrid Events

When a particular gene within a tissue type is identified by dNdScv as an SMG and is also identified by GISTIC2 as part of a SCNV, the SMG and SCNV were combined into a special event type called hybrid events. This reflects the assumption that the SCNV and SMG are exerting similar functional changes in the tumor cells. Supporting this assumption is the observation that oncogenes are frequently amplified and tumor suppressors are frequently deleted in the same cancer types. If the SCNV was a deletion the hybrid event was denoted as “gene-MD", for mutations/deletion, and if it was an amplification it was denoted as “gene-MA", for mutation/amplification. The hybrid events could take values in any of the mutation types or any of the copy number types. A single hybrid event could also take two values if it was observed as an SCNV and an SMG in the same patient. For example, if *CDKN2A* loss mutation and *CDKN2A* weak deletion co-occur in a patient, this observation would have been denoted as “L,WD". In such cases the penalty associated with the combined observation was the sum of the penalties of each observation independently (recall that penalties are -log probabilities). By increasing the penalty associated with co-occuring alterations of the same gene, the algorithm is encouraged to assign the event as a driver.

Two exceptions were encountered among the 19 TCGA cancer types that required special handling. In rectum adenocarcinoma (READ) *KRAS* was identified as part of an amplification peak and as part of a large deletion peak. Since *KRAS* is a known oncogene that is often part of amplification peaks in other cancer types we chose to represent *KRAS* as *KRAS-MA*. The deletion peak was retained in the dataset as an ordinary deletion event, but it was not annotated as a *KRAS* hybrid event. In kidney renal clear cell carcinoma (KIRC) both *ARID1* and *MTOR* were found to be part of the same deletion peak. Since *ARID1-MD* is observed in multiple cancer types whereas *MTOR-MD* is never observed in other cancer types, we chose to represent this deletion event as part of *ARID1-MD*.

#### S4 Four Phase Algorithm

The four phase procedure seeks to find the best scoring rule set of fixed size $K$, for $K \in \{1 \dots 40\}$. The core rule set is subsequently chosen from among the best rule sets of size $K$. Finding the best rule set of size $K$ from among $n$ rules is computationally intractable, and so we developed a heuristic procedure involving random sampling to approximate the global optima for each $K$. The four phase procedure involves several parameter choices that require specification. The methodology is presented using the default parameter values (Table S2) that were used for the presented applications to 19 TCGA cancer types.

**Phase 1: Stochastic Rule Prioritization** In phase 1, an iterative stochastic procedure is used to rank all of the rules in the rule library according to how likely they are to be included in the best performing rule set. For each rule, a *rule importance score* is calculated based on the average contribution of the rule within many random subsets of rules. Consider a set of rules $RS$. The contribution of rule $r_j$ within $RS$ is defined to be the percentage decrease in performance when $r_j$ is excluded from $RS$. Randomly sampled rule sets are allowed to contain family members, i.e., are not required to be valid, because we want to allow for direct competition between rules that are family members. To determine the rule importance score, multiple iterations (parameterized by *p1.spr*) of sampling multiple sizes of rule sets are evaluated separately in order to make fair comparisons between rules across a broad range of rule set sizes. For each rule set sample size, a $Z$ score is determined from the distribution of average contributions, and the rule importance score is determined to be the average of the $Z$ scores from different sampling sizes. The sizes of the random rule sets depend on the rule library size, as shown in Table S2. Once the importance scores are obtained for each rule in the rule pool, the 25% of rules (denoted as *p1.cut.size*) that have the smallest contribution scores are eliminated. The procedure proceeds until there are at most 24 rules, at which point the remaining rules are ranked according to a final round of importance score calculation. The purpose of using an iterative procedure, rather than ranking all the rules once is to enforce direct competition among the strongest scoring rules. *Our results on both simulations and real data* *show that the algorithm is highly robust to choice of p1.cut.size and p1.spr and that these choices matter more for very large rule libraries.*

**Phase 2: Exhaustive rule set evaluation** In phase 2, a subset of the top rules from phase 1 are exhaustively evaluated to determine the best rule set of each $K$. In contrast to phase 1, only valid rule sets that do not contain family members are considered in phase 2. For each $K$ the candidate rule pool is determined to be the maximum number of top rules that can be exhaustively evaluated with at most 200,000 rule sets. This computational parameter (denoted as *p2.mnrs*) was chosen to balance run time with depth of coverage. The number of rules that can be exhaustively evaluated for a given $K$ and *p2.mnrs* is referred to as the *pool size* and depends on the rule-family network between rules. For example, it is possible that no valid rule sets exists of size $K = 8$ within the top 20 rules, whereas $\approx 126,000$ valid rule sets would exist if none of the rules were family members. Since less computation is required to determine if a rule set is valid than to evaluate it, a larger computational parameter (*p2.max.compute*) is used to determine the pool sizes that lead to approximately *p2.mnrs*.

**Phase 3: Neighbor Rule Set Expansion** Phase 2 results in identification of best rules of sizes $K = 1 \dots 10$. However, because of computational constraints only a subset of top rules can be considered by phase 2 for inclusion among the best size-$K$ rule sets. In phase 3 the number of rules that can be included in the top rule sets is increased by making the assumption that the global best size-$K$ rule sets will overlap highly with the size-$K$ rule sets determined from phase 2. Consider as an example that in phase 2 the best rule set of size $K = 8$ is identified from among the top $n = 30$ rules. Denote this rule set as $RS_K$. The $d_L$ neighbors of $RS_K$ are defined to be the set of all rule sets that contain $K - L$ common rules with $RS_K$. In phase 3 the search for the best performing rule sets is expanded to include rule sets that are $d_L$ neighbors of $RS_K$ and contain $L$ rules from outside of the initial top 30 rule pool, for $L = 1, 2, 3$. For each $L$ the number of new rules that can be considered is determined subject to a computational constraint (*p3.mrns*). A similar expansion of candidate rule sets is performed by considering rule sets that overlap highly with $RS_{K-1}$, allowing for consideration of new rule sets of size $K$ that contain rules that are within $RS_{K-1}$ but absent from $RS_K$. This choice is motivated by the observation the best rule set of size $K$ tends to overlap highly with the best rule set of size $K - 1$.

**Phase 4: Expansion to Larger K** Phase 3 results in identification of best rules of sizes $K = 1 \dots 10$. In phase 4, the best rule set of size $K + 1$ is sought using a similar procedure to the neighbor expansion of phase 3. The top 3 rule sets of size $K$ are used as seed rule sets. The candidate rule sets of size $K + 1$ that are considered in phase 4 are the rule sets that share $K$, $K - 1$ or $K - 2$ rules in common one of the best 3 rule sets of size $K$. The maximum number of rule sets evaluated is constrained by the parameter *p4.mrns*. If for some $K'$ there are no valid rule sets that satisfy the $msa$ requirement, then the algorithm stops searching for larger rule sets, and the maximum $K$ is determined to be $K_{max} = K' - 1$.

#### S5 CRSO Report Overview from 19 TCGA Cancers

The CRSO reports in the supplemental folder "TCGA.REPORTS" provide a a detailed presentation of the CRSO findings for all 19 TCGA cancer types. Each report contains seven sections, which we prefix with SR to indicate that we are referring to sections in the supplemental reports. SR Section 1 presents a basic overview of the dataset and a summary table of the parameters. Default parameter values (Table S2) were used for all cancers, with 2 exceptions: in kidney renal clear cell carcinoma (KIRC), the rule coverage threshold and $msa$ were reduced to 1.4% because of sparsity of eligible rules, and in colon adenocarcinoma (COAD) the $max.nrs.p2$ and $max.considered.p2$ were increased to $10^6$ and $10^7$, respectively, because many of the top phase 1 rules were family members leading to a sparsity of valid phase 2 rule sets. SR Section 2 shows heatmaps of the binary matrix $\mathbf{D}$ and the penalty matrix $\mathbf{P}$ for the 20 most frequent events. SR Section 3 presents a summary of the K best rule sets identified by the four-phase procedure. SR Section 4 presents a deeper dive into the core rule set. SR Section 4.1 is a table showing different characteristics of the core rules, including the coverage, phase 1 importance rank and single rule performance. SR Section 4.2 shows an event-by-rule breakdown of the core rule set. SR Section 4.3 shows heatmaps of $\mathbf{P}$ before and after assignment to the core rule sets. This is a visual representation of how well the rule set accounts for observations in $\mathbf{D}$. SR Section 5 presents the generalized core analysis, consisting of GCRs, GCTs, GCDs and GCEs, SR Section 6 is a table summarizing the core RS rules and the consensus GCRs, similar to Table **??**. SR Section 7 is a dictionary of copy number events.

#### S6 Simulation Methods

In order to evaluate whether CRSO could identify the ground truth rule sets, ten simulation datasets were produced as described above for each $ntr \in \{2 \dots 20\}$, for a total of 190 simulations.

##### S6.1 Simulation Design

In order to challenge CRSO to solve problems similar to those presented by the real data, the passenger probability distributions, rule size distribution, and ground truth rule set network structure were informed by the characteristics of the 19 TCGA input datasets, and the CRSO results for these cancer types.

The size of each synthetic rule was sampled from a distribution of rule sizes of all 194 aggregated consensus GCRs (con-GCRs), denoted $\vec{d}_{size}$, so that each synthetic rule consisted of 2, 3, 4, 5 or 6 events with approximate probabilities 73%, 19%, 4%, 3% and 1% respectively. The distributions of events across the con-GCRs in the TCGA datasets were non-uniform, with some events being part of many rules and many events being part of a few rules. The distribution of events also depends on $ntr$, since rule sets with fewer rules generally comprise fewer total events. To address this, we define an event inclusion probability vector, $\vec{p}_{inc}$ for each simulation, based on the inclusion fractions of TCGA cancer types. The inclusion fraction of a cancer type is the vector of percentage rule inclusion of each event among the con-GCRs, normalized to sum to 1 and ordered from largest to smallest. $\vec{p}_{inc}$ is determined by taking the element-wise average of the inclusion fractions over the set of cancer types that comprise between $ntr - 5$ and $ntr + 5$ consensus GCRs, and then normalizing to sum to 1.

Given $ntr$, the following procedure was used to construct a simulated dataset:

1. **Determine the ground truth rule set, $RS_{GT}$:**
   (a) Make rule 1:
      i. Choose rule size $r_s$ by sampling from $\vec{d}_{size}$
      ii. Sample $r_s$ distinct events from the pool of events using probabilities $\vec{p}_{inc}$
   (b) For rule $j$ in 2 through $ntr$
      i. Make a random rule as in step 1
      ii. If rule $j$ is distinct and not a family member with rules $r_1 \ldots r_{j-1}$ proceed. Otherwise restart rule $j$.
2. **Assign samples to rules in $RS_{GT}$ and populate a matrix of true drivers, denoted by $D_0$:**
   (a) Determine the rule coverage distribution for the rules in $RS_{GT}$ such that each rule is assigned to at least 3% of samples. First each rule is assigned probability 3% and then the remaining probability is added by randomly partitioning the excess probability interval.
   (b) Determine a fraction of rules uniformly sampled between 0.01 and 0.20 that will be assigned to the null rule.
   (c) Initialize 100x400 matrix $D_0$ representing the ground truth drivers in each sample. Assign samples to rules or to the null rule according to the above probabilities. $D_0(i,j) = 1$ if and only if sample $j$ is assigned to a rule containing event $i$. Otherwise $D_0(i,j) = 0$
3. **Add noise sampled from empirical distributions to populate simulation matrix $D$ and $P$:**
   (a) Determine event-specific passenger rates $\vec{p_e}$ by sampling with replacement from the pool of TCGA passenger event rates. The pool of TCGA passenger event rates consisted of the pooled frequencies of all non-driver events in each cancer type, where driver events are any event that is part of at least one con-GCR. Passenger event rates below .02 were excluded from the pool in order to make the simulation more challenging to CRSO. Determine sample adjustment factors, $\vec{a_s}$ from the pool of TCGA passenger sample adjustment factors. TCGA passenger sample adjustment factors for each cancer type were defined as the fraction of non-driver events in each sample normalized across the tumor population to have mean 1. The pool of adjustment factors was the collection of all cancer-specific adjustment factors. $\vec{a_s}$ was sampled with replacement from the pool of adjustment factors and then renormalized to have mean 1.
   (b) Define a passenger probability matrix, $P_{prob}$, so that $P_{prob}(i,j) = min(p_e(i) * a_s(j), 0.95)$.
   (c) Use $P_{prob}$ to populate $D$ and $P$. For entry $(i,j)$:
      i. If $D_0(i,j) = 1$ then $D(i,j) = 1$ and $P(i,j) = -log(P_{prob}(i,j))$ .
      ii. If $D_0(i,j) = 0$ then generate a uniform random value $u$. If $u \leq P_{prob}(i,j)$ then $D(i,j) = 1$ and $P(i,j) = -log(P_{prob}(i,j))$ If $u > P_{prob}(i,j)$ then $D(i,j) = 0$ and $P(i,j) = 1$.

CRSO was applied to each simulation with the default parameters (Table S2), except for a few modifications to reduce computational cost. The $p1.ntpr$ was reduced from 40 to 20, max rule sets evaluated for phases 2, 3 and 4 were reduced by 50%, and the number of GC iterations was reduced from 100 to 40. The upper limit for the max library size was removed. The starting rule library sizes for the 190 simulations ranged from a low of 192 rules to a high of 5447 rules, and was between 350 and 2000 for 182 out of 190 simulations (Figure S4).

#### S6.2 Phase 1 Evaluation Methods

##### S6.2.1 Phase 1 Performance Score On Simulations

Given a rule library of $R_L$ rules, the phase 1 results can be represented as a ranking of rules, denoted by $\vec{r}_{p1}$. As a baseline for comparison, we order the rules in the rule library according to decreasing coverage (i.e., frequency of occurrence) across the population, and we denote this by $\vec{r}_{cov}$. Given knowledge of the ground truth rule set, we define $y(\vec{r}, x)$ to be the fraction of ground truth rules identified by position $x$, for all $x \in \{1 \ldots R_L\}$. We then define $AUC(\vec{r}) = \sum y(\vec{r}, x_k)$ from $x = 1 \ldots xmax$, where $xmax$ is minimum between 50 and the position of the last ground truth rule within $\vec{r}_{cov}$. The maximum possible AUC corresponds to a ranking in which all $ntr$ true rules are ranked in the top $ntr$ positions, and we denote this by $AUC_{max}$. The AUC of the frequency ranking is defined to be a baseline AUC: $AUC_0 = AUC(\vec{r}_{cov})$. Finally, we define the phase 1 score of the ranking $\vec{r}_{p1}$ to be $S_{p1}(\vec{r}_{p1}) = (AUC(\vec{r}_{p1}) - AUC_0)/(AUC_{max} - AUC_0)$. In case $AUC_0 = AUC_{max}$, we define $S_{p1}(\vec{r}_{p1}) = AUC(\vec{r}_{p1})/AUC_{max}$. This happened only 3 times in the 190 simulations, and $ntr$ equaled 2 for all 3 cases.

##### S6.2.2 Phase 1 Parameter Optimization

Phase 1 requires specification of 4 parameters, reviewed here:

1. $p1.cutsize$: the phase 1 cut size ($cs$ for short) is the fraction of lowest ranking rules eliminated in each phase 1 iteration. The default value of $cs = 0.25$ used in the simulations and TCGA experiments corresponds to 25% of lowest performing phase 1 rules being eliminated, or frozen in place, while the next iteration proceeds with the top 75% performing rules. Phase 1 ends when only 24 rules remain, at which point the final 24 rules are ranked producing a full ranking of all rules. The stopping point of 24 rules is itself a parameter called $p1.stop$. In the case of $cs = 1.0$, all of the rules in the rule library are ranked once, and then the top 24 rules are ranked a second time.
2. $p1.stop$: the stopping point of phase 1. This was chosen to be 24 because this is a number that can be exhaustively evaluated in phase 2 for $K \leq 10$.
3. $p1.ss.vec$: specifies the sizes of the random rule sets per rule for a single trial. The $p1.ss.vec$ was hardcoded so that $p1.ss.vec = [15, 25, 35]$ when $n_r > 300$, $p1.ss.vec = [8, 12, 16]$ when $n_r \in [150, 300)$, and $p1.ss.vec = [6, 8, 10, 12]$ when $n_r \leq 150$. Consider that there are more than 300 rules remaining (not yet eliminated), a single trial of rule $r$ involves sampling $r$ within 3 independent rule sets of sizes 15, 25 and 35, and calculating 3 corresponding contribution scores.
4. $p1.ntpr$: the number of trials per rule ($ntpr$ for short) within each phase 1 iteration. An $ntpr$ of 2 means that every rule's contribution score for each rule set size in $p1.ss.vec$ is determined by averaging of 2 random rule sets of each size. The average contribution scores for each rule set size are normalized over all of the remaining rules and the final importance score of each rule is determined by averaging the 3 or 4 Z-scores calculated for each rule.

To evaluate the impact of different parameter values on the CRSO results, phase 1 was performed using multiple parameter choices on each of the 190 simulations. Specifically, phase 1 was evaluated over a grid of parameter pairs for $p1.cutsize$ and $p1.ntpr$. The other two parameters, $p1.ss.vec$ and $p1.stop$ were fixed to the default values. We chose to fix $p1.ss.vec$ because it is an ensemble of 3 or 4 parameters designed to evaluate rules robustly across multiple rule set sizes. $p1.stop$ was fixed at 24 because this value allows for exhaustive evaluation for small $K$. Decreasing $p1.stop$ below a number that can be exhaustively evaluated would have no impact on the end CRSO results. Increasing $p1.stop$ would produce a minimal computational speed up while counter-acting the benefit of using an iterative procedure.

First we compared the performance of $cs = 1$, corresponding to all of the rules ranked in a single pass through, versus $cs = 0.5$, corresponding to 50% of rules being eliminated in each iteration. We found that $cs = 0.5$ was statistically superior over the 190 simulations for each $ntpr \in \{1, 2, 10, 20, 40\}$. We also found that was $ntprs = 10$ is significantly better than $ntprs \leq 2$ in for all $cs$ values. There was no statistical difference observed between $ntprs$ pairs ntprs= (40, 20), (20, 10) or (40, 10) for any cut size. There were no statistically significant difference between $cs$ pairs (.50, .25), (.25, .10) for any $ntprs$ values, the same was almost the case for for $(0.5, 0.1)$, except a statistical difference was observed for $ntprs = 1$. Since $ntprs =\leq 2$ is statistically inferior compared to $ntprs = 10$ for all cut sizes, we are not concerned about this statistical difference. We can conclude that a pairing of $(cs = 0.5, ntprs = 10)$ is optimal since no benefit is gained by reducing $cs$ or increasing $ntprs$, both of which would incur a computational cost. There was no statistical difference between $(cs = 0.5, ntprs = 10)$ and $(cs = 0.1, ntprs = 40)$. The results suggest that the parameters used in the simulations and in the TCGA experiments were sufficient to optimize phase one performance, but the same level of performance could have been achieved using less computation.

#### S7 Robustness to Missing Features

CRSO was applied to TCGA melanoma data excluding a single event, for each of the top 15 events. The results were compared to the results from the full dataset. We calculated several metrics for each event excluded. First we summarize the excluded event E by calculating: 1) frequency in the population, 2) percentage of GCDs containing E and 3) percentage of weighted GCDs, calculating by summing over the confidence scores of the GCDs that contain E and dividing by the sum of all GCD confidence scores. These are reflections of the event's importance to the full results, and can be thought of as unavoidable losses in the case of missing events (see Table S5, columns 2-4).

Next we calculate the agreement of the smaller dataset (i.e., dataset with event excluded) with the full dataset results that do not contain the event. Consider the truth to be the full input results. Eligible duos are those duos identified in the full results that do not contain E, since it is possible for these duos to by identified in the event-excluded results. Retention is the percentage of eligible duos from the full results that are also identified in the event-excluded results. Weighted retention accounts for the confidence scores of the retained/not retained rules, and is calculated by summing over the confidence scores of all retained duos and dividing by the sum of the confidence scores of all eligible duos.

Rules or duos that are missing from the full dataset can be gained in the smaller dataset results. Considering the full input results to be the ground truth, these gained duos are false positives. The false positive rate (FPR) for each exclusion experiment is the percentage of smaller dataset results that were identified in the full dataset results. Weighted FPR is summed confidence scores of the gained duos divided by the summed confidence scores of all duos identified by the smaller dataset.

The melanoma experiment found that weighted retention was over 93% for all exclusion experiments (Table S5). The weighted FPR was below 10% for all events except for *CDKN2A-MD* and *NRAS-M*, both of which are high frequency, highly included events. Overall the results suggest that if a driver event is excluded from the inputs, the duos/rules that do not contain the event will generally be comprehensively captured in the incomplete dataset. The rate of false associations caused by the event exclusion is generally small for most events, but can be large if the excluded event is frequent and part of many duos/rules in the full dataset.

### S8 Supplemental Tables

#### S8.1 Table S1: Mutation Types

**Table S1:** Distribution of mutation types observed across 19 TCGA cancers

| Mutation Type | Percentage of All Mutations[a] |
|---|---:|
| Missense_Mutation | 61.3 |
| Silent | 24.9 |
| Nonsense_Mutation | 4.65 |
| Splice_Site | 3.08 |
| Frame_Shift_Del | 2.85 |
| Frame_Shift_Ins | 0.941 |
| Intron | 0.597 |
| In_Frame_Del | 0.524 |
| RNA | 0.397 |
| IGR | 0.308 |
| 5'Flank | 0.165 |
| 3'UTR | 0.0797 |
| Nonstop_Mutation | 0.0677 |
| In_Frame_Ins | 0.0655 |
| 5'UTR | 0.0594 |
| De_novo_Start_OutOfFrame | 0.0124 |
| Start_Codon_Del | 0.0118 |
| De_novo_Start_InFrame | 0.0114 |
| Stop_Codon_Del | 0.00515 |
| Start_Codon_Ins | 0.00287 |
| Stop_Codon_Ins | 0.0015 |

[a] Mutations occurring in SMGs were excluded.

##### S8.2 Table S2: Default Parameters

**Table S2:** Default Parameters

| Parameter | Default Value | Description | Adjustable |
|---|---|---|---|
| rule.cov.thresh | 3% (minimum 6 samples) | Min rule coverage | yes, as desired |
| msa | 3% (minimum 6 samples) | Min samples assigned for every rule in RS | yes, as desired |
| max.lib.size | 2000 | Max # rules in rule library | yes, as desired |
| p1.ss.vec | depends on # rules $(n_r)^a$ | Sample sizes per P1 random sampling iterations | no |
| p1.spr | 40 | Random sets per rule in P1, for each ss.vec | yes, recommend $\geq$ 10 |
| p1.cut.size | 25% | Rules eliminated in each P1 iteration | yes, recommend $\leq$ 50% |
| p1.stop | 24 | Stop point for P1 | yes, not recommended |
| $K_2$ | 10 | Max K considered in P2 | yes, recommend $\leq$ 16 |
| p2.mnrs | 200,000 | Max # RS evaluated for each K in P2 | yes, rec. $\geq$ 20,000 |
| p2.max.compute | 5*p2.max.nrs | Max # non constrained rule sets considered for evaluation for each K in P2 | yes, rec. $\geq$ 2*p2.mnrs |
| p3.mnrs | 100,000 | Max # RS considered for each K in P3 | yes, rec. $\geq$ 20,000 |
| $K_4$ | 40 | Max K considered in P4 | yes, must be $\geq K_2$ + 1 |
| p4.mnrs | 100,000 | Max # RS considered for each K in P4 | yes, rec. $\geq$ 20,000 |
| core.cov.thresh | 90% | Coverage (relative to max) criteria for choosing core RS | yes, recommend $\geq$ 85% |
| core.perf.thresh | 90% | Performance (relative to max) criteria for choosing core RS | yes, recommend $\geq$ 85% |
| gc.iter | 100 | subsampling iterations for finding generalized cores | yes, recommend $\geq$ 10 |
| gc.eval | 100 | Rule sets evaluated per $K$ in each GC iteration | yes, recommend $\geq$ 10 |
| gc.sample.dist | uniform(67,85) | % sub sampled per GC iteration | no |
| gc.cov.thresh | uniform(85,99) | core coverage thresh per GC iteration | no |
| gc.perf.thresh | uniform(85,99) | core perf.thresh per GC iteration | no |

$^a ss.vec = [15, 25, 35]$ when $n_r > 300$, $ss.vec = [8, 12, 16]$ when $n_r \in [150, 300)$, $ss.vec = [6, 8, 10, 12]$ when $n_r \leq 150$

##### S8.3 Table S3: Phase 1 Performance on 190 Simulations

**Table S3:** Summary of CRSO Performance on Ground Truth Simulations

| Number True Rules | Mean Sensitivity | Mean Specificity | Mean Accuracy | Median Accuracy |
|---|---|---|---|---|
| 2 | 0.85 | 0.73 | 0.76 | 0.96 |
| 3 | 1 | 0.96 | 0.98 | 0.99 |
| 4 | 1 | 1 | 0.97 | 0.98 |
| 5 | 0.96 | 0.98 | 0.94 | 0.95 |
| 6 | 0.92 | 0.98 | 0.92 | 0.93 |
| 7 | 0.96 | 1 | 0.91 | 0.91 |
| 8 | 0.94 | 1 | 0.9 | 0.9 |
| 9 | 0.94 | 0.99 | 0.9 | 0.9 |
| 10 | 0.92 | 1 | 0.87 | 0.88 |
| 11 | 0.91 | 0.99 | 0.87 | 0.88 |
| 12 | 0.92 | 1 | 0.84 | 0.83 |
| 13 | 0.92 | 1 | 0.86 | 0.88 |
| 14 | 0.91 | 1 | 0.87 | 0.86 |
| 15 | 0.91 | 1 | 0.85 | 0.86 |
| 16 | 0.93 | 1 | 0.83 | 0.83 |
| 17 | 0.92 | 1 | 0.84 | 0.84 |
| 18 | 0.89 | 1 | 0.83 | 0.83 |
| 19 | 0.88 | 0.99 | 0.81 | 0.81 |
| 20 | 0.9 | 0.99 | 0.8 | 0.8 |
| All | 0.93 | 0.98 | 0.87 | 0.88 |

Ten simulations were performed for each ground truth rule set size. Mean sensitivity is the mean fraction of true rules identified among the con-GCRs. Mean specificity is the mean fraction of con-GCRs that are part of the ground truth rule set. Mean and median accuracy of each simulation. Simulation accuracy is the fraction of samples assigned assigned correctly by CRSO, using the core RS. The last row shows the mean/median values over all 190 simulations

##### S8.4 Table S4: Phase 1 Performance on 190 Simulations

**Table S4:** Phase 1 Performance on Ground Truth Simulations

| NTR | P1_Score | P1_Top_10 | P1_Top_20 | P1_Top_30 | P1_Top_40 | Cov_Top_40 | SJ_Top_40 |
|---|---|---|---|---|---|---|---|
| 2 | 0.99 | 1 | 1 | 1 | 1 | 0.9 | 0.85 |
| 3 | 0.98 | 1 | 1 | 1 | 1 | 0.9 | 0.7 |
| 4 | 0.98 | 1 | 1 | 1 | 1 | 0.78 | 0.7 |
| 5 | 0.96 | 0.98 | 1 | 1 | 1 | 0.88 | 0.72 |
| 6 | 0.97 | 0.97 | 1 | 1 | 1 | 0.85 | 0.78 |
| 7 | 0.98 | 0.96 | 1 | 1 | 1 | 0.71 | 0.69 |
| 8 | 0.96 | 0.91 | 0.99 | 0.99 | 1 | 0.75 | 0.62 |
| 9 | 0.96 | 0.89 | 1 | 1 | 1 | 0.72 | 0.69 |
| 10 | 0.94 | 0.85 | 0.97 | 0.98 | 0.99 | 0.65 | 0.6 |
| 11 | 0.97 | 0.83 | 0.99 | 1 | 1 | 0.74 | 0.66 |
| 12 | 0.97 | 0.77 | 1 | 1 | 1 | 0.69 | 0.64 |
| 13 | 0.97 | 0.71 | 1 | 1 | 1 | 0.57 | 0.5 |
| 14 | 0.97 | 0.66 | 0.98 | 1 | 1 | 0.56 | 0.49 |
| 15 | 0.95 | 0.6 | 0.93 | 0.99 | 1 | 0.44 | 0.39 |
| 16 | 0.94 | 0.56 | 0.95 | 1 | 1 | 0.51 | 0.49 |
| 17 | 0.94 | 0.54 | 0.91 | 0.99 | 0.99 | 0.36 | 0.38 |
| 18 | 0.94 | 0.54 | 0.91 | 0.98 | 0.99 | 0.38 | 0.38 |
| 19 | 0.97 | 0.51 | 0.94 | 0.99 | 1 | 0.49 | 0.49 |
| 20 | 0.94 | 0.46 | 0.86 | 0.98 | 0.99 | 0.28 | 0.26 |
| All | 0.96 | 0.78 | 0.97 | 0.99 | 1 | 0.64 | 0.58 |

Mean performance over ten iterations of ground truth rule set size (NTR). P1_Score is the mean phase 1 score, which compares the positions of ground truth rules in phase 1 rankings to the theoretical maximum ranking. P1_Top_X indicates the mean fraction of ground truth rules identified in the top X phase 1 rules. Nearly 100 percent of ground truth rules are identified within the top 30 rules, for all ntr between 2 and 20. Cov_Top_40 is the mean fraction of rules within the top 40 rules ranked according to coverage. SJ_Top_40 is the mean fraction of rules within the top 40 rules ranked according to single rule objective function score (i.e, SJ for Single rule J score). Phase 1 importance rankings are much better at prioritizing ground truth rules compared to either coverage or SJ rankings. Last row indicates mean over all 190 simulations.

##### S8.5 Table S5: Robustness to Event Exclusion in Melanoma

**Table S5:** Robustness of CRSO GCDs to Event Exclusion in Melanoma

| Excluded E | E Freq % | FR Incl E % | WFR Incl E % | FR Retention % | WFR Retention % | FPR % | WFPR % |
|---|---|---|---|---|---|---|---|
| BRAF-M | 50 | 35 | 40.5 | 94.3 | 98 | 31.4 | 9.53 |
| CDKN2A-MD | 46.2 | 25 | 22.4 | 83.9 | 96.3 | 19.4 | 19.9 |
| NRAS-M | 30 | 27.5 | 40.6 | 90.2 | 95.5 | 39 | 27.7 |
| ADAM18-M | 18.6 | 10 | 6.66 | 92.3 | 99.8 | 15.4 | 4.51 |
| snoU13-D | 17.6 | 7.5 | 2.37 | 77.4 | 98.3 | 3.23 | 0.667 |
| TP53-M | 16.6 | 7.5 | 11.3 | 92.3 | 96 | 12.8 | 1.34 |
| PTEN-MD | 16.6 | 5 | 8.59 | 87.2 | 93.7 | 15.4 | 9.1 |
| SMYD3-A | 15.5 | 10 | 4.29 | 62.1 | 93.2 | 13.8 | 3.48 |
| NOTCH2-A | 15.5 | 7.5 | 6.59 | 91.7 | 96.5 | 5.56 | 0.657 |
| RPTOR-A | 14.5 | 0 | 0 | 77.1 | 98.3 | 8.57 | 0.239 |
| d7(n=3) | 13.8 | 5 | 9.47 | 74.3 | 94.3 | 17.1 | 6.03 |
| B2M-MD | 13.4 | 7.5 | 8.51 | 85.7 | 98.5 | 8.57 | 1.75 |
| NF1-M | 13.1 | 2.5 | 1.26 | 95 | 97.7 | 7.5 | 1.19 |
| HULC-A | 13.1 | 2.5 | 5.26 | 81.1 | 94.7 | 13.5 | 2.19 |
| TERT-A | 13.1 | 0 | 0 | 82.9 | 98.5 | 2.86 | 0.961 |

**FR** Full results (rules or duos), i.e., results obtained using all inputs. **FR Inclusion E** is the percentage of FR that contain E. **WFR Inclusion E** is the sum of confidences of FR that contain E/ sum of confidences of FR. **FR Retention** is the percentage of eligible FR were retained among the new results. **WFR Retention** is the sum of confidences of retainted eligible FR / sum of confidences of eligible FR. **FPR** is the percentage of new results that were not in the full results (i.e., emurgent). **WFPR** is the sum of confidences of emurgent rules / sum of confidences all rules.

##### S8.6 Table S6: All con-GCRs from 19 TCGA tissues

**Table S6:** Consensus GCRs from 19 TCGA Cancer Types

| Tissue | Rule | P1 Rank | Conf. | Cov. | SJ | NE |
|---|---|---|---|---|---|---|
| BLCA | *FGFR3-MA + KDM6A-MD* | 5 | 100 | 10.7% (r=42) | 332 (r=12) | 2 |
| BLCA | *ERBB2-MA + TP53-M* | 6 | 100 | 12.8% (r=22) | 330 (r=10) | 2 |
| BLCA | *CDKN2A-MD + FGFR3-MA* | 1 | 99 | 11.2% (r=33) | 388 (r=17) | 2 |
| BLCA | *RB1-M + TP53-M* | 4 | 97 | 13.8% (r=15) | 336 (r=3) | 2 |
| BLCA | *ARID1A-MD + PIK3CA-M* | 11 | 97 | 8.16% (r=99) | 292 (r=47) | 2 |
| BLCA | *a1(n=4) + ENSA/SNORA40-A + TP53-M* | 12 | 96 | 12.8% (r=23) | 287 (r=8) | 3 |
| BLCA | *PIK3CA-M + TP53-M* | 3 | 89 | 11.7% (r=28) | 343 (r=6) | 2 |
| BLCA | *CDKN2A-MD + KDM6A-MD* | 7 | 89 | 15.1% (r=11) | 319 (r=15) | 2 |
| BLCA | *CDKN2A-MD + TP53-M* | 2 | 88 | 16.3% (r=6) | 382 (r=1) | 2 |
| BLCA | *ERBB4/RNA5SP119-D + KCNJ13-D* | 15 | 81 | 19.1% (r=1) | 278 (r=28) | 2 |
| BLCA | *ARID1A-MD + SOX4-A + TP53-M* | 37 | 79 | 7.14% (r=172) | 204 (r=46) | 3 |
| BLCA | *CASC8-A + YWHAZ-A* | 61 | 79 | 13.3% (r=17) | 176 (r=99) | 2 |
| BLCA | *ELF3-M + KDM6A-MD + TP53-M* | 104 | 53 | 3.57% (r=1393) | 152 (r=199) | 3 |
| BLCA | *CDKN2A-MD + NFE2L2-M* | 26 | 51 | 5.61% (r=337) | 220 (r=131) | 2 |
| BLCA | *ARID1A-MD + CDKN2A-MD* | 10 | 57 | 13.3% (r=18) | 300 (r=27) | 2 |
| BRCA | *MAP3K1-M + PIK3CA-MA* | 4 | 100 | 3.74% (r=245) | 478 (r=49) | 2 |
| BRCA | *AQP11/CLNS1A-A + CCND1/ORAOV1-A* | 6 | 100 | 12.6% (r=3) | 457 (r=7) | 2 |
| BRCA | *IKBKB/POLB-A + ZNF703-A* | 10 | 100 | 8.93% (r=10) | 424 (r=36) | 2 |
| BRCA | *CDH1-M + PIK3CA-MA* | 2 | 99 | 5.61% (r=71) | 676 (r=10) | 2 |
| BRCA | *PTEN-MD + TP53-M* | 7 | 99 | 6.96% (r=32) | 448 (r=17) | 2 |
| BRCA | *ERBB2-MA + VMP1-A* | 11 | 95 | 9.35% (r=7) | 420 (r=28) | 2 |
| BRCA | *d2(snoU13)-D + THYN1-D* | 12 | 94 | 12.3% (r=4) | 414 (r=31) | 2 |
| BRCA | *ERBB2-MA + TP53-M* | 8 | 83 | 8.41% (r=14) | 446 (r=4) | 2 |
| BRCA | *PIK3CA-MA + TP53-M* | 1 | 72 | 13.5% (r=1) | 911 (r=1) | 2 |
| BRCA | *CCND1/ORAOV1-A + PIK3CA-MA* | 3 | 66 | 11.3% (r=5) | 627 (r=3) | 2 |
| BRCA | *MYC-A + TP53-M* | 5 | 62 | 12.8% (r=2) | 462 (r=2) | 2 |
| BRCA | *GATA3-M + VMP1-A* | 23 | 54 | 3.43% (r=307) | 344 (r=174) | 2 |
| BRCA | *FBLN5-D + ZFP36L1-MD* | 41 | 52 | 5.61% (r=68) | 300 (r=178) | 2 |
| CESC | *NTM/RNU6ATAC12P-D + TMEM136-D* | 1 | 100 | 18.3% (r=1) | 131 (r=1) | 2 |
| CESC | *PIK3CA-M + PTEN-MD* | 2 | 100 | 5.24% (r=32) | 113 (r=8) | 2 |
| CESC | *LINC00393-A + RASA3-A* | 10 | 100 | 5.76% (r=20) | 69.4 (r=30) | 2 |
| CESC | *a11(n=6) + ZNF750-MD* | 37 | 89 | 3.66% (r=66) | 43.1 (r=78) | 2 |
| CESC | *FNDC3B/RN7SL141P-A + PTEN-MD* | 14 | 84 | 3.14% (r=159) | 59.2 (r=70) | 2 |
| CESC | *GCDH/SYCE2-A + STK11-MD* | 18 | 84 | 3.66% (r=68) | 53.4 (r=84) | 2 |
| CESC | *FAT1-MD + FNDC3B/RN7SL141P-A* | 15 | 78 | 5.24% (r=25) | 58.7 (r=53) | 2 |
| CESC | *FNDC3B/RN7SL141P-A + KCNJ13/RN7SL359P-D* | 34 | 78 | 6.81% (r=14) | 44 (r=34) | 2 |
| CESC | *a6(n=5) + PIK3CA-M* | 5 | 91 | 6.28% (r=18) | 77.1 (r=7) | 2 |
| CESC | *BCL2L1/COX4I2-A + PIK3CA-M* | 7 | 77 | 3.66% (r=100) | 73.5 (r=27) | 2 |
| CESC | *FGF3-A + LRP1B-D* | 52 | 75 | 3.66% (r=74) | 37.7 (r=97) | 2 |
| CESC | *EP300-M + PIK3CA-M* | 9 | 74 | 4.71% (r=42) | 71.6 (r=15) | 2 |
| CESC | *a6(n=5) + FBXW7-M* | 16 | 74 | 3.14% (r=111) | 58.1 (r=48) | 2 |
| CESC | *HLA-B-M + PIK3CA-M* | 33 | 72 | 3.66% (r=97) | 44.3 (r=38) | 2 |
| CESC | *a2(n=6) + KCNJ13/RN7SL359P-D + TMEM136-D* | 57 | 71 | 5.24% (r=30) | 36.6 (r=14) | 3 |
| COAD | *APC-M + PTEN-M + SMAD4-MD* | 34 | 97 | 20.4% (r=33) | 711 (r=28) | 3 |
| COAD | *ATM-M + CTNNB1-M + KRAS-M + PIK3CA-M + PTEN-M + TP53-M* | 147 | 92 | 4.7% (r=1074) | 443 (r=293) | 6 |
| COAD | *BRAF-M + PIH1/WWOX-D + RBFOX1-D* | 6 | 81 | 5.52% (r=735) | 1030 (r=817) | 3 |
| COAD | *APC-M + ATM-M + PIK3CA-M + PTEN-M* | 16 | 78 | 16.9% (r=51) | 825 (r=22) | 4 |
| COAD | *ATM-M + PIK3CA-M + SMAD4-MD + TP53-M* | 87 | 64 | 11.3% (r=151) | 512 (r=68) | 4 |
| COAD | *RBFOX1-D + TP53-M* | 8 | 60 | 25.4% (r=18) | 999 (r=67) | 2 |
| COAD | *APC-M + TP53-M* | 1 | 56 | 52.5% (r=1) | 1330 (r=1) | 2 |
| COAD | *BRAF-M + PIK3CA-M + PTEN-M + SMAD4-MD + TP53-M* | 196 | 56 | 5.25% (r=847) | 403 (r=342) | 5 |
| COAD | *APC-M + KRAS-M + PIK3CA-M* | 7 | 53 | 24.3% (r=20) | 1010 (r=7) | 3 |
| COAD | *APC-M + d2(n=4)* | 37 | 51 | 23.5% (r=22) | 697 (r=108) | 2 |
| ESCA | *MYC-A + TP53-M* | 6 | 100 | 33.2% (r=5) | 340 (r=5) | 2 |
| ESCA | *GMDS-D + PIH1/WWOX-D + TP53-M* | 13 | 99 | 18.5% (r=42) | 276 (r=19) | 3 |
| ESCA | *SMAD4-MD + TP53-M* | 15 | 94 | 19% (r=36) | 272 (r=35) | 2 |
| ESCA | *CDKN2A-D + FGF3-A + TP53-M* | 2 | 91 | 26.6% (r=10) | 428 (r=4) | 3 |

| | | | | | | |
|---|---|---|---|---|---|---|
| ESCA | *ACTRT3/MYNN-A + FGF3-A + IMMP2L/LRRN3-D +PTPRN2-D + TP53-M* | 17 | 87 | 7.07% (r=797) | 261 (r=157) | 5 |
| ESCA | *MIR5707/MIR595/PTPRN2-D + PIH1/WWOX-D* | 140 | 84 | 16.3% (r=53) | 139 (r=486) | 2 |
| ESCA | *TP53-M + ZNF750-MD* | 23 | 71 | 13.6% (r=102) | 242 (r=108) | 2 |
| ESCA | *LRP1B-D + TP53-M* | 7 | 65 | 32.6% (r=6) | 336 (r=7) | 2 |
| ESCA | *ACTRT3/MYNN-A + NFE2L2-M + TP53-M* | 71 | 59 | 5.98% (r=1280) | 169 (r=345) | 3 |
| ESCA | *d9(n=6) + FHIT/NPCDR1/U3-D + IMMP2L/LRRN3-D + PIH1/WWOX-D + TP53-M* | 112 | 51 | 5.98% (r=1378) | 149 (r=260) | 5 |
| GBM | *CDKN2A-D + PTEN-MD* | 1 | 100 | 27.8% (r=2) | 557 (r=2) | 2 |
| GBM | *CDK4/MARCH9/TSPAN31-A + CPM/MDM2-A* | 9 | 99 | 7.69% (r=29) | 256 (r=47) | 2 |
| GBM | *CDK4/MARCH9/TSPAN31-A + TP53-M* | 6 | 89 | 6.96% (r=37) | 304 (r=36) | 2 |
| GBM | *CDKN2A-D + NF1-MD* | 12 | 82 | 9.89% (r=20) | 212 (r=24) | 2 |
| GBM | *CDKN2A-D + EGFR-M + SNORA73-A* | 3 | 77 | 14.7% (r=8) | 395 (r=4) | 3 |
| GBM | *PTEN-MD + TP53-M* | 5 | 77 | 12.1% (r=13) | 308 (r=9) | 2 |
| GBM | *CDKN2A-D + PIK3R1-M* | 15 | 70 | 8.06% (r=27) | 200 (r=30) | 2 |
| GBM | *ATRX-M + IDH1-M + TP53-M* | 11 | 63 | 3.66% (r=131) | 221 (r=43) | 3 |
| GBM | *CDKN2A-D + PDGFRA-MA* | 8 | 59 | 11% (r=15) | 269 (r=16) | 2 |
| GBM | *EGFR-M + PTEN-MD + SNORA73-A* | 30 | 52 | 6.96% (r=40) | 143 (r=14) | 3 |
| HNSC | *FAT1-MD + TP53-M* | 4 | 98 | 27.1% (r=3) | 827 (r=5) | 2 |
| HNSC | *CDKN2A-MD + FAT1-MD + NOTCH1-MD* | 25 | 74 | 8.51% (r=83) | 400 (r=45) | 3 |
| HNSC | *CASP8-M + HRAS-M* | 8 | 73 | 3.76% (r=549) | 621 (r=302) | 2 |
| HNSC | *RN7SKP265-A + TP53-M* | 7 | 69 | 26.9% (r=4) | 671 (r=6) | 2 |
| HNSC | *NOTCH1-MD + TP53-M* | 13 | 64 | 15.4% (r=15) | 516 (r=16) | 2 |
| HNSC | *CDKN2A-MD + PPFIA1-A + TP53-M* | 2 | 52 | 21.2% (r=8) | 892 (r=2) | 3 |
| HNSC | *CDKN2A-MD + NFE2L2-MA + TP53-M* | 24 | 50 | 8.71% (r=77) | 406 (r=35) | 3 |
| KIRC | *PBRM1-M + VHL-MD* | 1 | 100 | 20.1% (r=1) | 615 (r=1) | 2 |
| KIRC | *RNU6ATAC4P-D + VHL-MD* | 2 | 100 | 15% (r=3) | 475 (r=3) | 2 |
| KIRC | *RNA5SP200-A + VHL-MD* | 3 | 100 | 16.6% (r=2) | 459 (r=2) | 2 |
| KIRC | *ARID1A-MD + VHL-MD* | 4 | 100 | 7.16% (r=5) | 205 (r=5) | 2 |
| KIRC | *CAHM/QKI-D + PARK2-D* | 6 | 100 | 4.39% (r=12) | 190 (r=17) | 2 |
| KIRC | *BAP1-M + VHL-MD* | 5 | 98 | 5.54% (r=6) | 201 (r=8) | 2 |
| KIRC | *PBRM1-M + RNA5SP200-A* | 9 | 97 | 7.16% (r=4) | 166 (r=10) | 2 |
| KIRC | *MTOR-M + VHL-MD* | 8 | 76 | 3.93% (r=15) | 169 (r=11) | 2 |
| KIRC | *NRXN3-D + VHL-MD* | 15 | 74 | 2.54% (r=22) | 98.4 (r=19) | 2 |
| KIRC | *PBRM1-M + SETD2-M* | 17 | 65 | 4.39% (r=13) | 95.3 (r=13) | 2 |
| KIRC | *BAP1-M + RNA5SP200-A* | 18 | 50 | 3% (r=18) | 88.4 (r=26) | 2 |
| LGG | *CIC-M + IDH1-M* | 1 | 100 | 19.3% (r=5) | 2610 (r=5) | 2 |
| LGG | *EGFR-M + SNORA73-A* | 10 | 86 | 3.9% (r=132) | 655 (r=153) | 2 |
| LGG | *IDH1-M + PIK3CA-M* | 8 | 84 | 6.04% (r=61) | 695 (r=42) | 2 |
| LGG | *IDH1-M + ISOC2/NAT14/ZNF628-D + TP53-M* | 13 | 82 | 10.3% (r=15) | 591 (r=9) | 3 |
| LGG | *ATRX-MD + IDH1-M + TP53-M* | 2 | 71 | 37.4% (r=4) | 2520 (r=1) | 3 |
| LGG | *FUBP1-M + IDH1-M* | 6 | 59 | 8.77% (r=21) | 889 (r=21) | 2 |
| LIHC | *ARID1A-MD + MIR4689-D* | 1 | 100 | 14.8% (r=1) | 209 (r=1) | 2 |
| LIHC | *ALB-M + CTNNB1-M* | 4 | 98 | 4.37% (r=48) | 177 (r=12) | 2 |
| LIHC | *CTSS-A + TP53-MD* | 5 | 98 | 7.92% (r=5) | 176 (r=7) | 2 |
| LIHC | *LINC00676-A + PCCA-A* | 12 | 98 | 13.4% (r=2) | 134 (r=5) | 2 |
| LIHC | *CTSS-A + RB1-MD* | 16 | 94 | 6.83% (r=6) | 124 (r=23) | 2 |
| LIHC | *ARID1A-MD + CTNNB1-M* | 7 | 91 | 6.28% (r=15) | 162 (r=8) | 2 |
| LIHC | *CTNNB1-M + RN7SKP226-A* | 11 | 86 | 6.28% (r=14) | 135 (r=9) | 2 |
| LIHC | *RN7SKP226-A + TP53-MD* | 6 | 85 | 8.74% (r=3) | 166 (r=4) | 2 |
| LIHC | *ALB-M + TP53-MD* | 15 | 82 | 3.55% (r=86) | 128 (r=27) | 2 |
| LIHC | *RN7SKP96-D + TACR3-D* | 29 | 82 | 4.64% (r=31) | 92.8 (r=77) | 2 |
| LIHC | *RB1-MD + TP53-MD* | 2 | 50 | 8.47% (r=4) | 201 (r=2) | 2 |
| LIHC | *C19orf77/NFIC-D + TP53-MD* | 9 | 80 | 6.83% (r=7) | 143 (r=10) | 2 |
| LIHC | *CCND1/ORAOV1-A + TP53-MD* | 8 | 80 | 4.64% (r=38) | 152 (r=17) | 2 |
| LIHC | *CTNNB1-M + RB1-MD* | 20 | 53 | 4.37% (r=49) | 112 (r=21) | 2 |
| LIHC | *LRP1B-D + TP53-MD* | 19 | 52 | 5.19% (r=26) | 112 (r=24) | 2 |
| LIHC | *PTEN-MD + TP53-MD* | 25 | 50 | 4.64% (r=37) | 98.3 (r=22) | 2 |
| LUAD | *SFTA3-A + TP53-M* | 5 | 97 | 15.5% (r=1) | 362 (r=5) | 2 |
| LUAD | *CDKN2A-MD + KRAS-MA* | 7 | 97 | 8.16% (r=16) | 307 (r=10) | 2 |
| LUAD | *KRAS-MA + SFTA3-A* | 4 | 95 | 10.7% (r=10) | 362 (r=6) | 2 |
| LUAD | *CDKN2A-MD + TP53-M* | 8 | 95 | 13.4% (r=5) | 305 (r=4) | 2 |
| LUAD | *KRAS-MA + STK11-M* | 3 | 90 | 10% (r=11) | 380 (r=3) | 2 |
| LUAD | *EGFR-MA + TP53-M* | 2 | 74 | 11.5% (r=9) | 408 (r=2) | 2 |
| LUAD | *ATM-M + KRAS-MA* | 6 | 73 | 6.07% (r=48) | 338 (r=16) | 2 |

| | | | | | | |
|---|---|---|---|---|---|---|
| LUAD | *KRAS-MA + TP53-M* | 1 | 68 | 15.5% (r=2) | 593 (r=1) | 2 |
| LUAD | *CDH10-M + TP53-M* | 10 | 67 | 13.8% (r=4) | 289 (r=8) | 2 |
| LUAD | *NF1-M + TP53-M* | 11 | 77 | 8.79% (r=15) | 270 (r=18) | 2 |
| LUAD | *BRAF-M + TP53-M* | 13 | 66 | 5.23% (r=70) | 240 (r=41) | 2 |
| LUSC | *TP53-M + WHSC1L1-A* | 3 | 100 | 21.3% (r=16) | 299 (r=17) | 2 |
| LUSC | *FOXP1/MIR1284-D +ROBO1-D + PROS1/STX19-D + ROBO2-D + TP53-M* | 17 | 97 | 6.74% (r=456) | 193 (r=111) | 5 |
| LUSC | *CSMD3-M + NFE2L2-MA + SOX2-A* | 30 | 79 | 8.43% (r=230) | 152 (r=126) | 3 |
| LUSC | *RB1-MD + SOX2-A + TP53-M* | 10 | 64 | 8.43% (r=251) | 218 (r=59) | 3 |
| LUSC | *PIK3CA-M + SOX2-A + TP53-M* | 11 | 63 | 8.43% (r=252) | 216 (r=64) | 3 |
| LUSC | *CDKN2A-MD + EGFR-A + LRP1B-D + SOX2-A + TP53-M* | 18 | 61 | 6.18% (r=605) | 185 (r=84) | 5 |
| LUSC | *CERS3-A + TP53-M* | 44 | 55 | 15.2% (r=37) | 135 (r=50) | 2 |
| LUSC | *TP53-M + TSPAN4-D* | 21 | 90 | 15.7% (r=33) | 172 (r=35) | 2 |
| LUSC | *CDH10-M + TP53-M* | 12 | 68 | 16.3% (r=30) | 212 (r=34) | 2 |
| LUSC | *NF1-MD + TP53-M* | 16 | 65 | 15.7% (r=34) | 194 (r=38) | 2 |
| OV | *RN7SL501P-D + TP53-M* | 5 | 100 | 31.4% (r=9) | 895 (r=8) | 2 |
| OV | *BSPH1-D + RN7SL526P-D + TCF3-D + TP53-M* | 6 | 100 | 11.9% (r=472) | 862 (r=85) | 4 |
| OV | *d2(n=15) + FKSG52/PDE4D-D + MECOM-A + MYC-A + TP53-M* | 8 | 99 | 13.2% (r=339) | 717 (r=39) | 5 |
| OV | *d15(n=5) + PPP2R2A-D + TP53-M* | 15 | 96 | 12.5% (r=387) | 617 (r=159) | 3 |
| OV | *BRD4-A + TP53-M* | 11 | 95 | 24.2% (r=27) | 687 (r=24) | 2 |
| OV | *d6(n=6) + MECOM-A + MYC-A + TP53-M* | 76 | 92 | 12.7% (r=373) | 456 (r=108) | 4 |
| OV | *MYC-A + TCF3-D + TP53-M* | 13 | 89 | 30.5% (r=11) | 674 (r=5) | 3 |
| OV | *CBX8-A + MYC-A* | 117 | 82 | 25.1% (r=23) | 405 (r=389) | 2 |
| OV | *MECOM-A + TCF3-D + TP53-M* | 9 | 66 | 29.2% (r=14) | 713 (r=6) | 3 |
| OV | *CCNE1-A + RN7SL566P/SAMD4B-A + TP53-M* | 31 | 56 | 13% (r=349) | 539 (r=119) | 3 |
| OV | *ANKS1B/FAM71C/RNA5SP366-D + d13(n=19) + TP53-M* | 55 | 52 | 12.7% (r=367) | 488 (r=208) | 3 |
| OV | *d2(n=15) + FKSG52/PDE4D-D + TCF3-D* | 275 | 51 | 18.5% (r=97) | 321 (r=432) | 3 |
| PAAD | *KRAS-M + TP53-M* | 1 | 95 | 61.1% (r=1) | 754 (r=1) | 2 |
| PAAD | *CDKN2A-MD + KRAS-M* | 3 | 67 | 45.2% (r=2) | 518 (r=3) | 2 |
| PAAD | *KRAS-M + SMAD4-MD* | 4 | 63 | 28.6% (r=5) | 332 (r=6) | 2 |
| PRAD | *FAM92B-D + ZFHX3-D* | 2 | 100 | 13.8% (r=3) | 296 (r=8) | 2 |
| PRAD | *RNY1P8-D + ZC3H13-D* | 3 | 93 | 16.1% (r=2) | 282 (r=6) | 2 |
| PRAD | *FOXA1-M + ZNF292-D* | 9 | 84 | 3.86% (r=187) | 207 (r=102) | 2 |
| PRAD | *CHD1-D + SPOP-M + ZNF292-D* | 11 | 72 | 6.5% (r=49) | 195 (r=3) | 3 |
| PRAD | *PTEN-MD + TP53-M* | 5 | 69 | 5.28% (r=85) | 258 (r=36) | 2 |
| PRAD | *ERG-D + TMPRSS2-MD* | 1 | 66 | 19.9% (r=1) | 373 (r=1) | 2 |
| READ | *APC-MD + INS/MIR4686/TH-A + KRAS-MA* | 13 | 83 | 5.83% (r=459) | 231 (r=240) | 3 |
| READ | *APC-MD + KRAS-MA + TP53-M* | 3 | 79 | 35.8% (r=4) | 521 (r=2) | 3 |
| READ | *APC-MD + d2(n=4)* | 6 | 74 | 30.8% (r=6) | 278 (r=14) | 2 |
| READ | *a1(n=8) + APC-MD + CTNNBL1-A + TP53-M* | 16 | 72 | 8.33% (r=194) | 221 (r=88) | 4 |
| READ | *APC-MD + PIK3CA-M + TP53-M* | 4 | 71 | 25.8% (r=12) | 430 (r=5) | 3 |
| READ | *APC-MD + d3(n=623) + TP53-M* | 10 | 65 | 21.7% (r=20) | 239 (r=8) | 3 |
| READ | *APC-MD + SMAD4-MD* | 12 | 65 | 21.7% (r=19) | 237 (r=28) | 2 |
| READ | *KRAS-MA + PIK3CA-M* | 56 | 64 | 16.7% (r=33) | 147 (r=36) | 2 |
| READ | *d2(n=4) + KRAS-MA + PARK2-D + TP53-M* | 192 | 55 | 4.17% (r=1011) | 90.9 (r=339) | 4 |
| SKCM | *NRAS-M + TP53-M* | 4 | 100 | 6.55% (r=29) | 168 (r=8) | 2 |
| SKCM | *BRAF-M + RN7SKP254-A* | 11 | 98 | 6.55% (r=32) | 137 (r=21) | 2 |
| SKCM | *BRAF-M + HIPK2/TBXAS1-A* | 6 | 97 | 7.93% (r=13) | 161 (r=10) | 2 |
| SKCM | *CDKN2A-MD + NRAS-M* | 2 | 84 | 14.1% (r=2) | 327 (r=2) | 2 |
| SKCM | *BRAF-M + PTEN-MD* | 3 | 84 | 10.7% (r=3) | 227 (r=3) | 2 |
| SKCM | *BRAF-M + CDKN2A-MD* | 1 | 83 | 26.2% (r=1) | 524 (r=1) | 2 |
| SKCM | *HULC-A + NRAS-M* | 5 | 71 | 6.55% (r=28) | 167 (r=9) | 2 |
| SKCM | *B2M-MD + FMN1/SNORD77/snoU13-D* | 10 | 66 | 8.97% (r=6) | 144 (r=36) | 2 |
| SKCM | *BRAF-M + TP53-M* | 8 | 51 | 8.28% (r=9) | 156 (r=7) | 2 |
| STAD | *ARID1A-MD + PIK3CA-M* | 1 | 100 | 9.97% (r=38) | 508 (r=34) | 2 |
| STAD | *d4(n=4) + TP53-M* | 2 | 100 | 17.9% (r=3) | 408 (r=3) | 2 |
| STAD | *ARID1A-MD + TP53-M* | 6 | 96 | 13.8% (r=14) | 319 (r=13) | 2 |
| STAD | *CCSER1-D + PIH1/WWOX-D + TP53-M* | 9 | 91 | 10.5% (r=32) | 300 (r=7) | 3 |
| STAD | *ARID1A-MD + KRAS-MA* | 11 | 89 | 8.18% (r=78) | 292 (r=70) | 2 |
| STAD | *SMAD4-MD + TP53-M* | 5 | 83 | 11.5% (r=23) | 340 (r=17) | 2 |
| STAD | *GMDS-D + TP53-M* | 10 | 76 | 14.6% (r=10) | 296 (r=8) | 2 |
| STAD | *KRAS-MA + TP53-M* | 12 | 67 | 8.18% (r=82) | 289 (r=39) | 2 |

| | | | | | | |
|---|---|---|---|---|---|---|
| STAD | *ERBB2-A + FKSG52/PDE4D-D + PIH1/WWOX-D + TP53-M* | 79 | 66 | 5.12% (r=369) | 173 (r=60) | 4 |
| STAD | *PIH1/WWOX-D + PTPRD/RN7SL5P/SNORD27-D* | 21 | 64 | 13.8% (r=13) | 255 (r=25) | 2 |
| STAD | *d4(n=4) + IMMP2L/LRRN3-D +PTPRN2-D + PIH1/WWOX-D* | 110 | 57 | 3.32% (r=1236) | 156 (r=337) | 4 |
| STAD | *ARID1A-MD + SMAD4-MD* | 22 | 55 | 7.93% (r=84) | 255 (r=170) | 2 |
| STAD | *ARID1A-MD + FHIT/NPCDR1/U3-D + PIH1/WWOX-D* | 14 | 52 | 7.42% (r=106) | 281 (r=57) | 3 |
| STAD | *PIH1/WWOX-D + PIK3CA-M* | 19 | 50 | 7.42% (r=104) | 268 (r=56) | 2 |
| UCEC | *PIK3R1-M + PTEN-MD* | 2 | 97 | 28.9% (r=2) | 534 (r=2) | 2 |
| UCEC | *ARID1A-MD + CTNNB1-M + PTEN-MD* | 38 | 81 | 9.92% (r=45) | 189 (r=17) | 3 |
| UCEC | *SNORD37-D + TP53-M* | 8 | 79 | 15.7% (r=10) | 366 (r=25) | 2 |
| UCEC | *PIK3CA-M + PTEN-MD* | 1 | 78 | 36.8% (r=1) | 749 (r=1) | 2 |
| UCEC | *CTNNB1-M + PIK3CA-M* | 6 | 75 | 16.5% (r=7) | 421 (r=8) | 2 |
| UCEC | *ARID1A-MD + KRAS-M + PTEN-MD* | 28 | 65 | 10.7% (r=37) | 214 (r=14) | 3 |
| UCEC | *PIK3CA-M + TP53-M* | 5 | 74 | 13.2% (r=18) | 439 (r=16) | 2 |

**SJ** Single rule objective function score. **NE** Number of events in rule.

##### S8.7 Table S7: SELECT Common Event Duos Comparison

**Table S7:** Duos Identified by SELECT or CRSO Across 16 TCGA Cancer Types

| Tissue | Mut1 | Mut2 | Algorithm | CovCRSO | CovSELECT | DuoConf |
|---|---|---|---|---|---|---|
| STAD | *ARID1A* | *RNF43* | **SELECT Only** | 8.70 | 10.00 | 2 |
| BLCA | *KDM6A* | *STAG2* | **SELECT Only** | 7.90 | 6.00 | 11 |
| BLCA | *FGFR3* | *STAG2* | **SELECT Only** | 6.10 | 5.10 | 0 |
| HNSC | *CASP8* | *FAT1* | **SELECT Only** | 5.50 | 2.60 | 5 |
| LUAD | *KEAP1* | *STK11* | **SELECT Only** | 5.20 | 2.60 | 10 |
| LUAD | *KRAS* | *RBM10* | **SELECT Only** | 3.60 | 4.30 | 0 |
| HNSC | *FAT1* | *RASA1* | **SELECT Only** | 2.20 | 1.30 | 0 |
| CESC | *HLA-A* | *HLA-B* | **SELECT Only** | 2.10 | 2.10 | 0 |
| BLCA | *C3orf70* | *CREBBP* | **SELECT Only** | 2.00 | 1.30 | 0 |
| BLCA | *CDKN1A* | *RB1* | **SELECT Only** | 2.00 | 2.60 | 0 |
| BLCA | *RBM10* | *STAG2* | **SELECT Only** | 1.80 | 1.30 | 0 |
| BRCA | *CDH1* | *ERBB2* | **SELECT Only** | 1.60 | 1.00 | 0 |
| LUAD | *RBM10* | *STK11* | **SELECT Only** | 1.50 | 2.20 | 0 |
| BRCA | *CDH1* | *FOXA1* | **SELECT Only** | 0.93 | 0.92 | 0 |
| BRCA | *CDH1* | *TBX3* | **SELECT Only** | 0.83 | 0.82 | 0 |
| BRCA | *CDH1* | *RUNX1* | **SELECT Only** | 0.83 | 0.82 | 0 |
| BRCA | *CBFB* | *GATA3* | **SELECT Only** | 0.62 | 0.61 | 0 |
| BRCA | *AKT1* | *GATA3* | **SELECT Only** | 0.10 | 0.92 | 0 |
| BRCA | *AKT1* | *MAP2K4* | **SELECT Only** | 0.10 | 0.31 | 0 |
| BLCA | *FGFR3* | *KDM6A* | **BOTH** | 11.00 | 6.00 | 100 |
| HNSC | *FAT1* | *NOTCH1* | **BOTH** | 10.00 | 4.60 | 74 |
| LUAD | *KRAS* | *STK11* | **BOTH** | 10.00 | 6.50 | 99 |
| BRCA | *CDH1* | *PIK3CA* | **BOTH** | 5.60 | 6.20 | 99 |
| KIRC | *PBRM1* | *SETD2* | **BOTH** | 4.40 | 6.40 | 65 |
| LGG | *IDH1* | *TP53* | **CRSO Only** | 46.00 | 40.00 | 100 |
| LGG | *ATRX* | *IDH1* | **CRSO Only** | 40.00 | 18.00 | 100 |
| LGG | *ATRX* | *TP53* | **CRSO Only** | 39.00 | 15.00 | 71 |
| UCEC | *PIK3CA* | *PTEN* | **CRSO Only** | 37.00 | 30.00 | 100 |
| UCEC | *PIK3R1* | *PTEN* | **CRSO Only** | 29.00 | 24.00 | 100 |
| UCEC | *ARID1A* | *PTEN* | **CRSO Only** | 28.00 | 24.00 | 93 |
| HNSC | *FAT1* | *TP53* | **CRSO Only** | 27.00 | 14.00 | 100 |
| UCEC | *ARID1A* | *PIK3CA* | **CRSO Only** | 24.00 | 17.00 | 58 |
| UCEC | *CTNNB1* | *PTEN* | **CRSO Only** | 23.00 | 19.00 | 93 |
| LUSC | *NFE2L2* | *TP53* | **CRSO Only** | 22.00 | 12.00 | 99 |
| LUSC | *PTEN* | *TP53* | **CRSO Only** | 22.00 | 4.50 | 68 |
| KIRC | *PBRM1* | *VHL* | **CRSO Only** | 20.00 | 18.00 | 100 |
| ESCA | *SMAD4* | *TP53* | **CRSO Only** | 19.00 | 4.90 | 98 |
| LGG | *CIC* | *IDH1* | **CRSO Only** | 19.00 | 11.00 | 100 |
| BLCA | *KDM6A* | *TP53* | **CRSO Only** | 17.00 | 12.00 | 85 |
| UCEC | *CTNNB1* | *PIK3CA* | **CRSO Only** | 17.00 | 14.00 | 99 |
| UCEC | *KRAS* | *PTEN* | **CRSO Only** | 17.00 | 14.00 | 87 |
| LUSC | *NF1* | *TP53* | **CRSO Only** | 16.00 | 2.80 | 99 |
| BLCA | *ARID1A* | *TP53* | **CRSO Only** | 15.00 | 8.10 | 87 |
| HNSC | *NOTCH1* | *TP53* | **CRSO Only** | 15.00 | 5.60 | 65 |
| LUAD | *KRAS* | *TP53* | **CRSO Only** | 15.00 | 9.10 | 100 |
| UCEC | *KRAS* | *PIK3CA* | **CRSO Only** | 15.00 | 13.00 | 82 |
| BLCA | *RB1* | *TP53* | **CRSO Only** | 14.00 | 12.00 | 100 |
| ESCA | *TP53* | *ZNF750* | **CRSO Only** | 14.00 | 2.70 | 86 |
| STAD | *ARID1A* | *TP53* | **CRSO Only** | 14.00 | 7.10 | 96 |
| BLCA | *ERBB2* | *TP53* | **CRSO Only** | 13.00 | 4.70 | 100 |
| BRCA | *PIK3CA* | *TP53* | **CRSO Only** | 13.00 | 7.10 | 73 |
| PRAD | *PTEN* | *TMPRSS2* | **CRSO Only** | 13.00 | 0.00 | 86 |
| UCEC | *PIK3CA* | *TP53* | **CRSO Only** | 13.00 | 10.00 | 100 |
| UCEC | *ARID1A* | *KRAS* | **CRSO Only** | 13.00 | 11.00 | 87 |
| UCEC | *ARID1A* | *CTNNB1* | **CRSO Only** | 13.00 | 10.00 | 86 |
| BLCA | *PIK3CA* | *TP53* | **CRSO Only** | 12.00 | 8.10 | 89 |
| GBM | *PTEN* | *TP53* | **CRSO Only** | 12.00 | 3.40 | 94 |
| LUAD | *EGFR* | *TP53* | **CRSO Only** | 12.00 | 5.20 | 74 |
| LUSC | *RB1* | *TP53* | **CRSO Only** | 12.00 | 4.50 | 99 |

| | | | | | | |
|---|---|---|---|---|---|---|
| STAD | *SMAD4* | *TP53* | **CRSO Only** | 12.00 | 2.40 | 83 |
| GBM | *EGFR* | *PTEN* | **CRSO Only** | 11.00 | 1.90 | 52 |
| HNSC | *PIK3CA* | *TP53* | **CRSO Only** | 11.00 | 8.30 | 75 |
| HNSC | *NFE2L2* | *TP53* | **CRSO Only** | 11.00 | 2.60 | 53 |
| LUSC | *PIK3CA* | *TP53* | **CRSO Only** | 11.00 | 9.60 | 63 |
| SKCM | *BRAF* | *PTEN* | **CRSO Only** | 11.00 | 3.80 | 100 |
| STAD | *ARID1A* | *PIK3CA* | **CRSO Only** | 10.00 | 9.30 | 100 |
| LGG | *FUBP1* | *IDH1* | **CRSO Only** | 8.80 | 4.60 | 59 |
| LUAD | *NF1* | *TP53* | **CRSO Only** | 8.80 | 5.70 | 77 |
| LIHC | *RB1* | *TP53* | **CRSO Only** | 8.50 | 2.10 | 50 |
| BRCA | *ERBB2* | *TP53* | **CRSO Only** | 8.40 | 0.31 | 83 |
| SKCM | *BRAF* | *TP53* | **CRSO Only** | 8.30 | 5.00 | 52 |
| BLCA | *ARID1A* | *PIK3CA* | **CRSO Only** | 8.20 | 4.70 | 97 |
| ESCA | *NFE2L2* | *TP53* | **CRSO Only** | 8.20 | 7.10 | 84 |
| LUAD | *KEAP1* | *TP53* | **CRSO Only** | 8.20 | 0.87 | 61 |
| STAD | *ARID1A* | *KRAS* | **CRSO Only** | 8.20 | 2.60 | 89 |
| STAD | *KRAS* | *TP53* | **CRSO Only** | 8.20 | 2.40 | 68 |
| STAD | *ARID1A* | *SMAD4* | **CRSO Only** | 7.90 | 1.10 | 55 |
| KIRC | *ARID1A* | *VHL* | **CRSO Only** | 7.20 | 0.95 | 100 |
| BRCA | *PTEN* | *TP53* | **CRSO Only** | 7.00 | 1.20 | 99 |
| SKCM | *NRAS* | *TP53* | **CRSO Only** | 6.60 | 4.70 | 100 |
| LIHC | *ARID1A* | *CTNNB1* | **CRSO Only** | 6.30 | 1.00 | 91 |
| LUAD | *ATM* | *KRAS* | **CRSO Only** | 6.10 | 3.50 | 73 |
| LGG | *IDH1* | *PIK3CA* | **CRSO Only** | 6.00 | 5.30 | 84 |
| GBM | *RB1* | *TP53* | **CRSO Only** | 5.90 | 3.70 | 66 |
| BLCA | *ELF3* | *TP53* | **CRSO Only** | 5.60 | 1.30 | 53 |
| KIRC | *BAP1* | *VHL* | **CRSO Only** | 5.50 | 4.00 | 98 |
| BLCA | *ELF3* | *KDM6A* | **CRSO Only** | 5.40 | 1.70 | 53 |
| PRAD | *PTEN* | *TP53* | **CRSO Only** | 5.30 | 0.00 | 69 |
| CESC | *PIK3CA* | *PTEN* | **CRSO Only** | 5.20 | 3.70 | 100 |
| LUAD | *BRAF* | *TP53* | **CRSO Only** | 5.20 | 4.30 | 66 |
| GBM | *IDH1* | *TP53* | **CRSO Only** | 4.80 | 3.40 | 100 |
| CESC | *EP300* | *PIK3CA* | **CRSO Only** | 4.70 | 2.60 | 74 |
| LIHC | *PTEN* | *TP53* | **CRSO Only** | 4.60 | 1.60 | 50 |
| LIHC | *CTNNB1* | *RB1* | **CRSO Only** | 4.40 | 0.00 | 53 |
| GBM | *ATRX* | *TP53* | **CRSO Only** | 4.00 | 0.00 | 63 |
| HNSC | *CASP8* | *HRAS* | **CRSO Only** | 3.80 | 0.99 | 73 |
| BRCA | *MAP3K1* | *PIK3CA* | **CRSO Only** | 3.70 | 3.70 | 100 |
| CESC | *HLA-B* | *PIK3CA* | **CRSO Only** | 3.70 | 1.00 | 72 |
| GBM | *ATRX* | *IDH1* | **CRSO Only** | 3.70 | 0.00 | 63 |

Duos identified by SELECT (i.e., predicted synergies) and high confidence GCDs (conf. $\geq 50$) identified by CRSO among the common mutations for each cancer type.

### S9 Supplemental Figures

#### S9.1 Figure S1: Candidate Driver SMGs and SCNVs per Sample

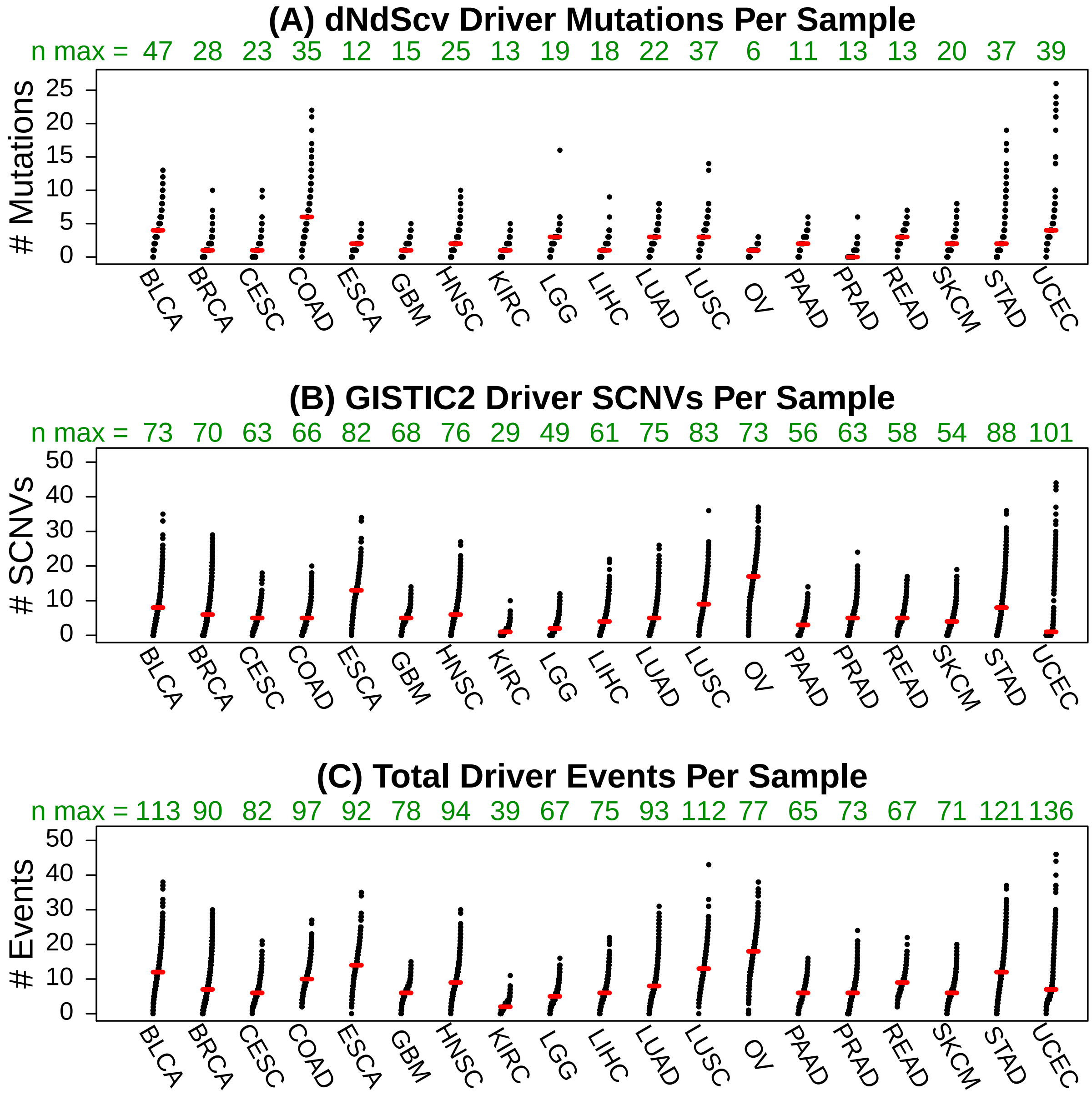

**Figure S1:** Number of driver events per patients across 19 TCGA cancer types. Panel (A) shows SMGs identified by dNdScv, panel (B) shows SCNVs identified by GISTIC2. Green numbers above each plot show the total number of candidate drivers identified by dNdScv (A) and GISTIC2 (B). Panel (C) shows total driver events from both methods. The number of total drivers per cancer type was less than the sum of the SCNVs and SMGs because some SCNVs and SMGs were combined into hybrid events. Red bars indicate median alterations per sample.

##### S9.2 Figure S2: Simulation Example

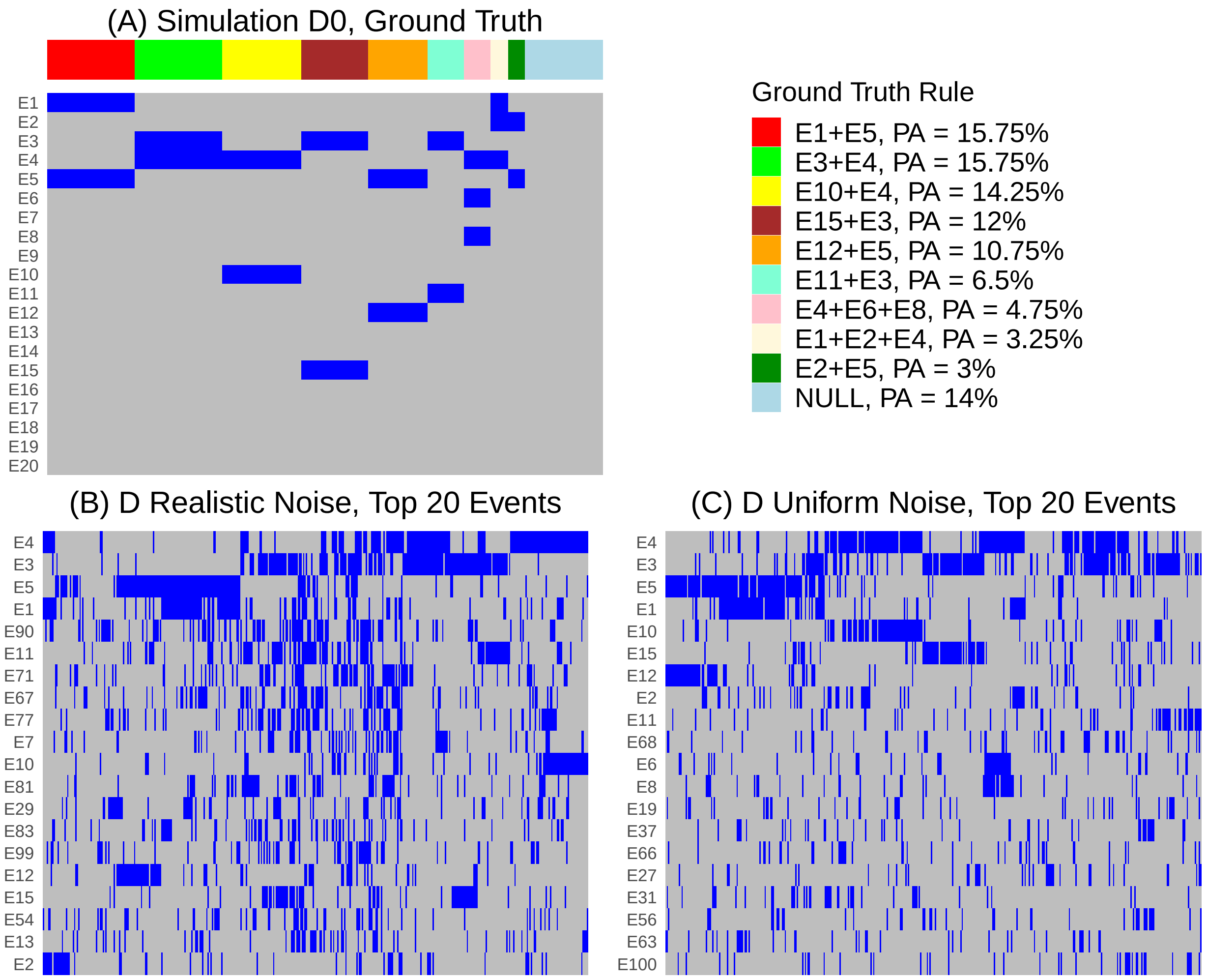

**Figure S2:** Example of randomly generated simulation with known ground truth of 9 rules. The simulation dataset consists of 400 samples and 100 events (top 20 most frequent events shown). (A) D0 matrix of ground truth driver events. . Legend to the right shows the rules along with percentage of samples assigned (PA). (B) Addition of empirical noise based pooled passenger event rates and sample passenger adjustment factors from 19 TCGA tissue types. (C) Addition of uniform noise equal to the mean of the realistic noise, approximately 8% uniform passenger probability.

##### S9.3 Figure S3: Core RS Assignment Agreement with Ground Truth

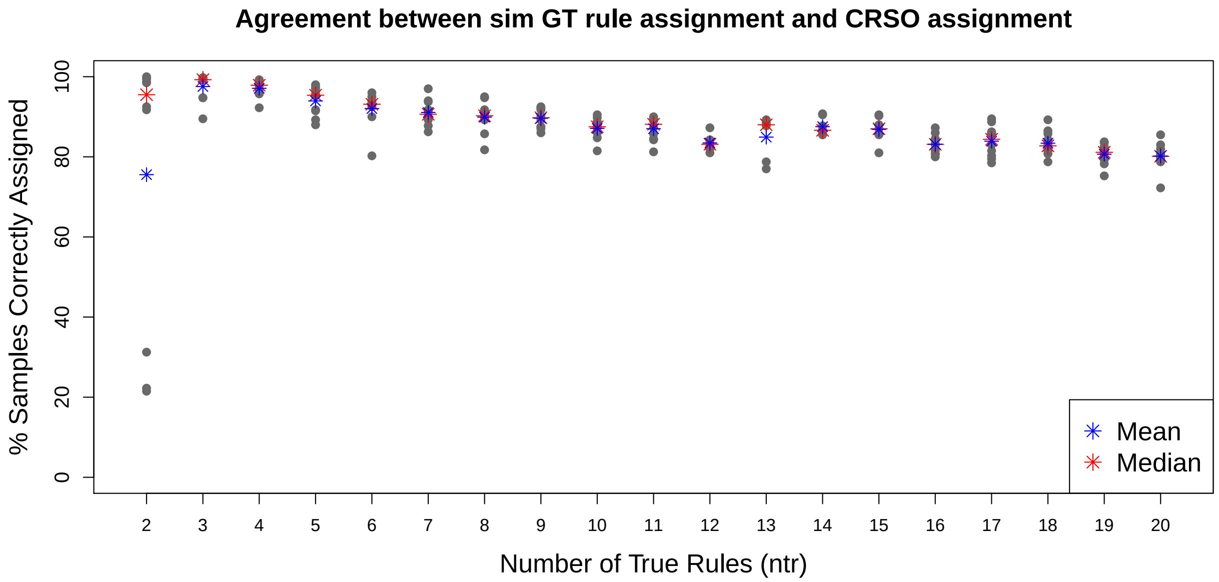

**Figure S3:** Assignment accuracy of each simulation. CRSO assignments were determined according to the core rule set. Blue/red stars show mean/median accuracy for each ntr value.

##### S9.4 Figure S4: Rule Library Sizes for Simulations

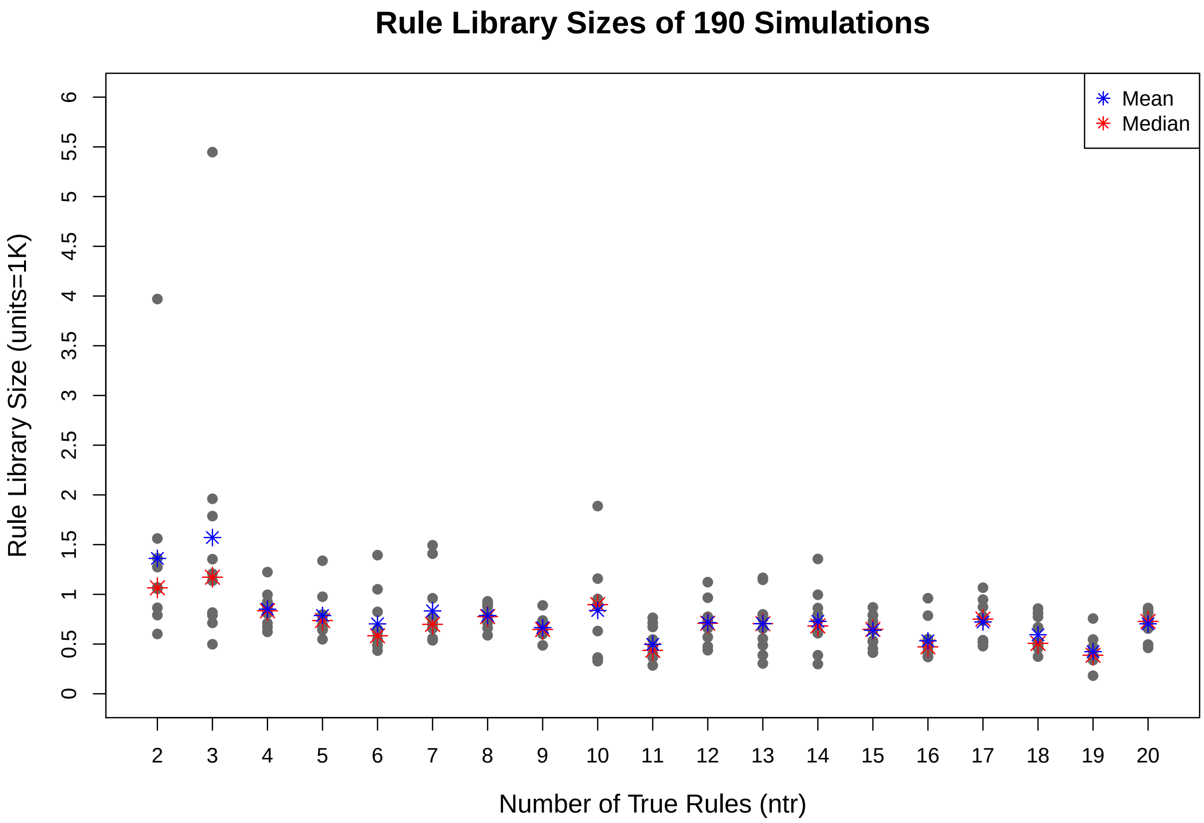

**Figure S4:** Rule library sizes for the 190 simulations, broken down according to number of true rules (ntr). The largest median library sizes were observed for $ntr = 3(1172.5)$ and $ntr = 2(1065.5)$. For $ntr \in \{4 \ldots 20\}$ the median rule library size ranged from 387.5 ($ntr = 19$) to 896.5 ($ntr = 10$).

##### S9.5 Figure S5: Simulation performance of Fisher tests under different passenger probabilities

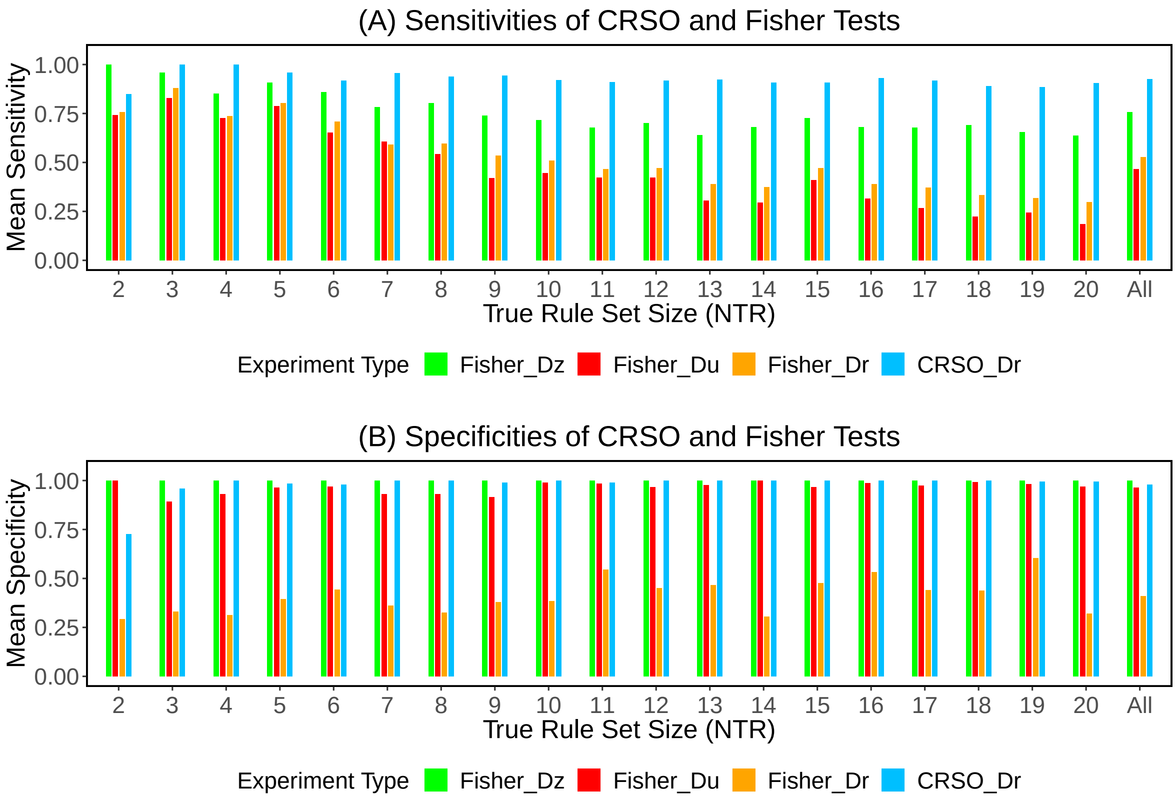

**Figure S5:** Mean Sensitivities and Specificities of consensus GCRs Grouped by True RS Size.

##### S9.6 Figure S6: Melanoma Core RS Identification

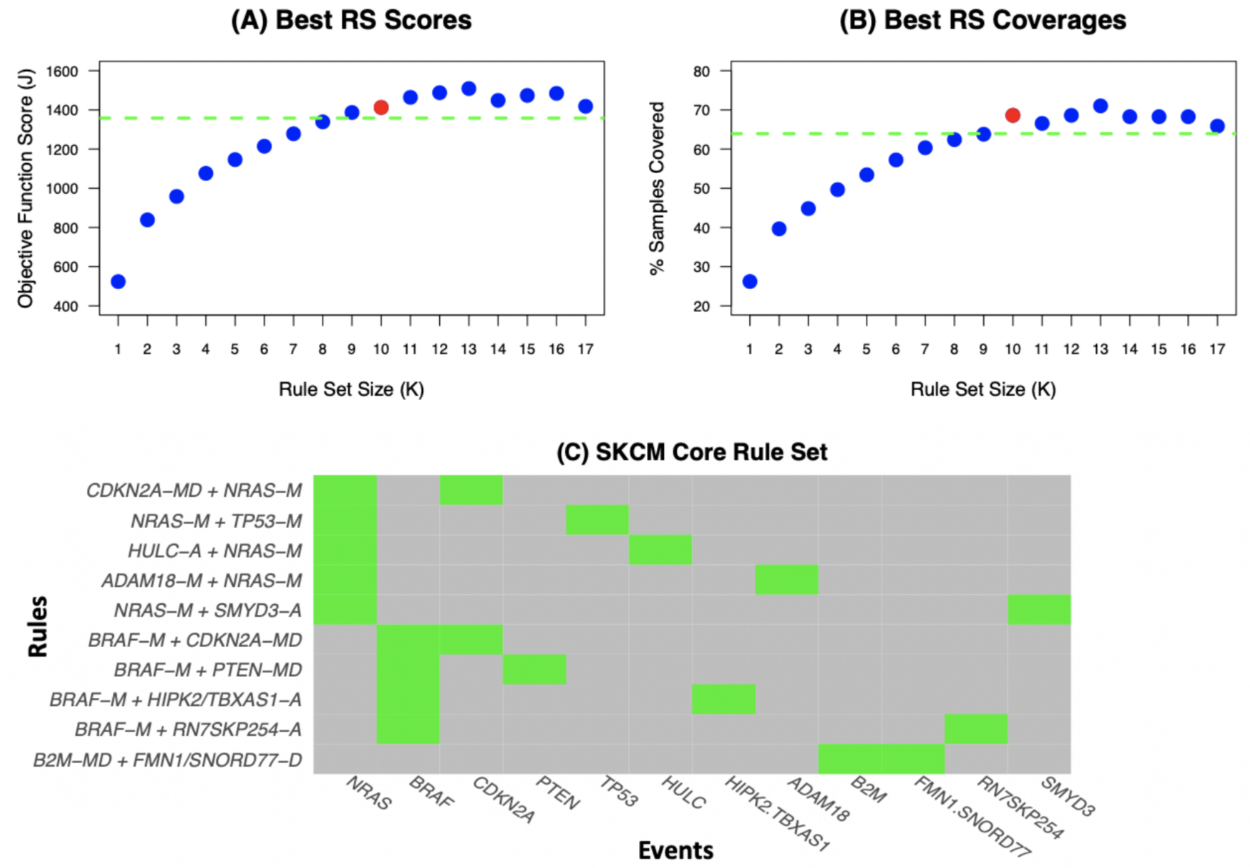

**Figure S6:** Melanoma CRSO results. A–B) The objective function score (A) and coverage (B) of the best filtered rule sets. The core rule set corresponds to $K = 10$ and is shown in red. The dashed green lines are the thresholds for determining the core rule set. The maximum $K$ shown is the largest $K$ for which valid rule sets that satisfy minimum sample assignment of 9 samples (3%). C) Composition of the core rule set.

##### S9.7 Figure S7: Melanoma Generalized Cores

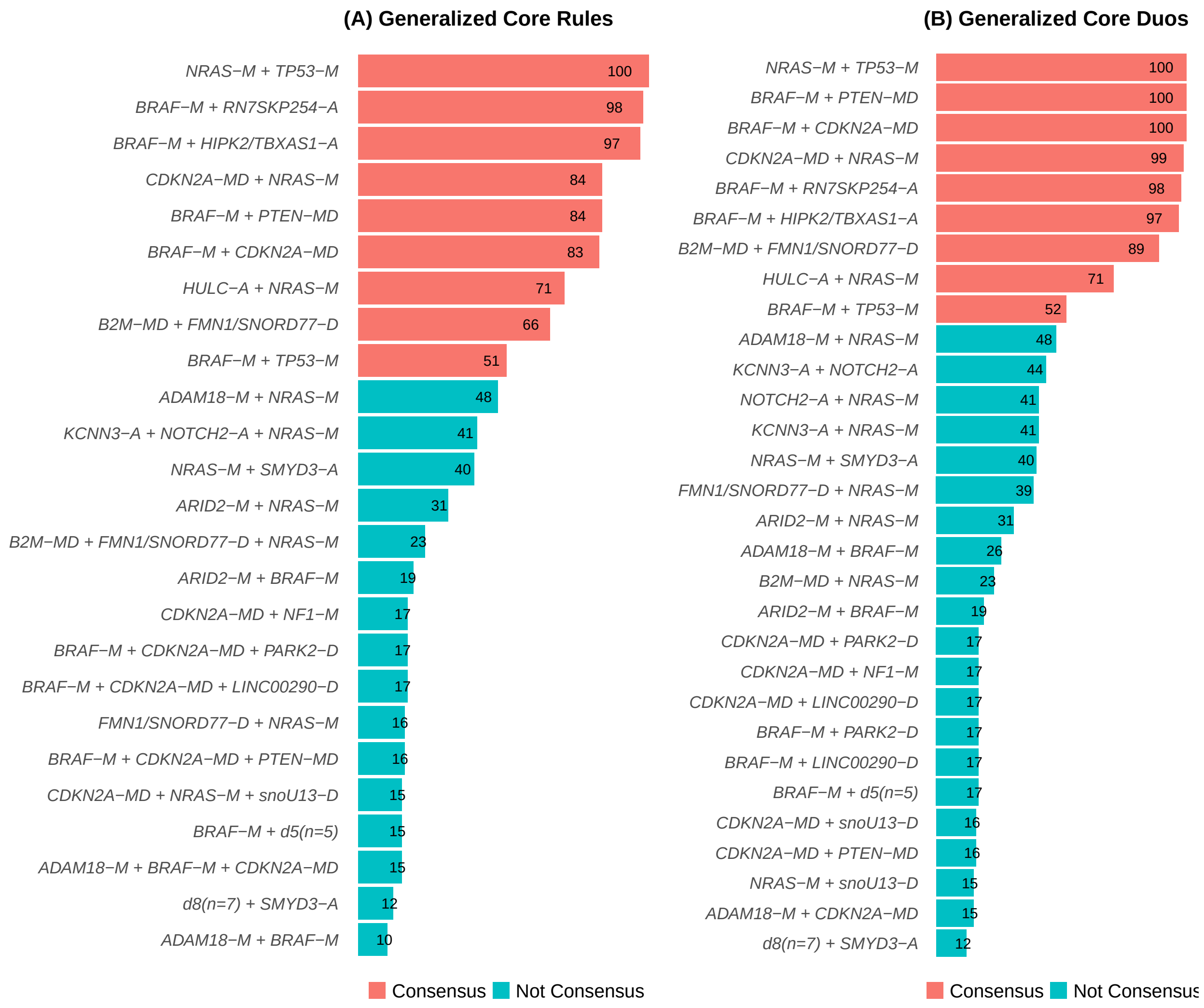

**Figure S7:** Summary of generalized core (GC) results for melanoma. Bars show confidence levels, which are the percentage of sub-sample iterations containing the observation. Generalized core rules (A) and generalized core duos (B) that achieve a minimum confidence level of 5 are shown.

##### S9.8 Figure S8: Performance and Coverage Convergence for 19 TCGA Cancers

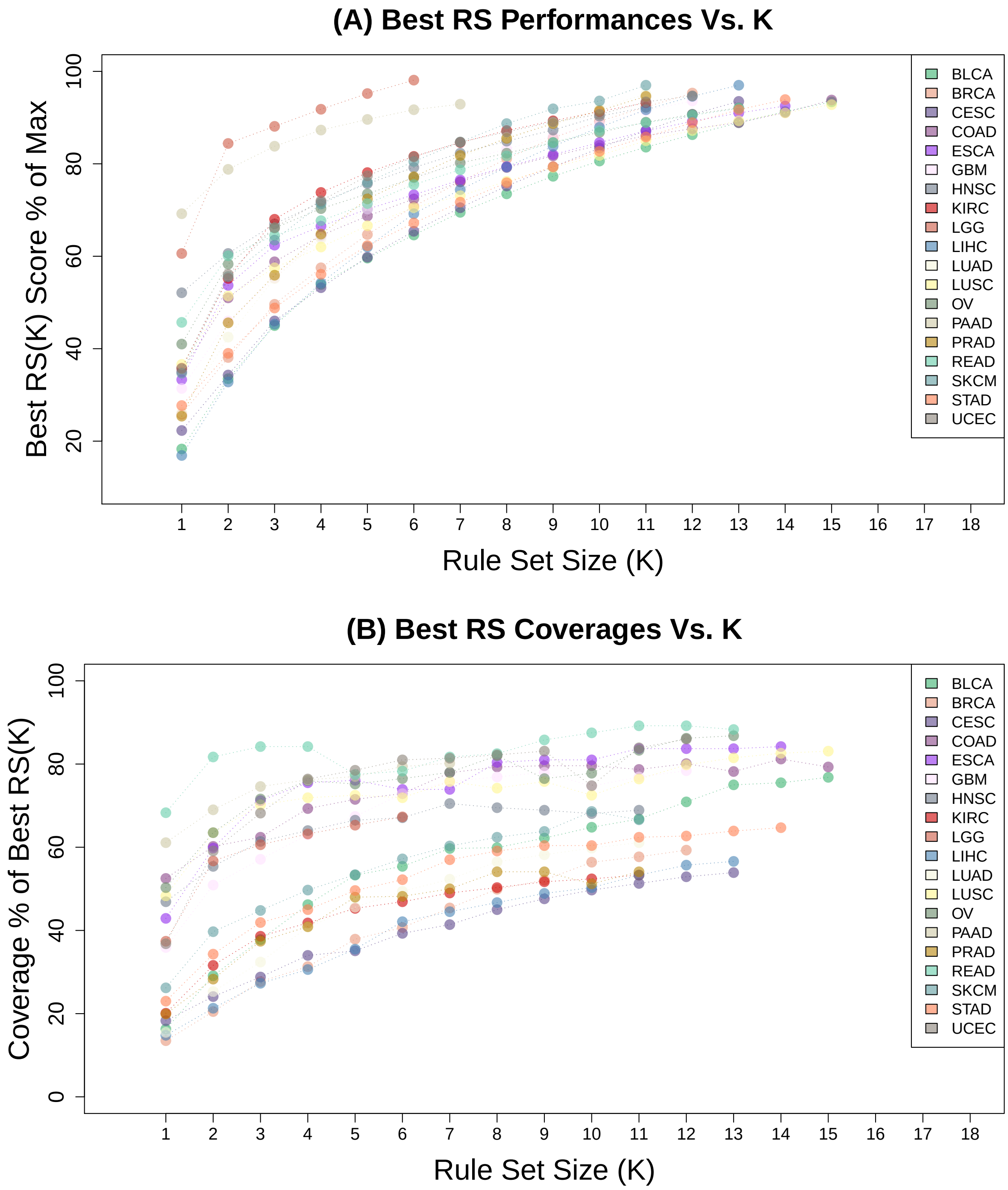

**Figure S8:** Performance and Coverage Convergence for Different Cancers. For each cancer the maximum $K$ plotted is 2 more than $K_{core}$.
