## Supplementary material for "Identifying Modules of Cooperating Cancer Drivers": CRSO reports for 19 TCGA cancer types: CRSO_Report_PRAD.pdf

### CRSO Output Report: PRAD

Michael Klein

Apr 11 2020

**5 Generalized Core Analysis** **9**

**6 Combined Core Table** **13**

**7 Dictionary of Copy Number Events** **14**

### 1 Dataset Overview and Parameters

**Number of samples = 492.**
**Number of events = 73.**
**Rule coverage requirement = 15 samples.**
**Rule assignment requirement = 15 samples.**
**Rule library size = 301 rules.**

#### 1.1 Parameter table

| Parameter Name | Value | Description | Category |
|---|---|---|---|
| n.cores | 20 | Number HPC Cores | Resources |
| msa | 15 | Minimum Samples Assigned | RS Constraint |
| rule.thresh | 0.0305 | Rule Coverage Thresh | Library Definition |
| max.rl | 2000 | Max Rule Library Size | Library Definition |
| p1.ntpr | 40 | P1 Num Trial Per Rule | Phase 1 |
| p1.stop | 24 | P1 Stop Elimination | Phase 1 |
| cut.size | 0.25 | P1 Cut Size | Phase 1 |
| k.max.2 | 10 | P2 Max K | Phase 2 |
| max.nrs.p2 | 200000 | P2 Max RS Evaluated Per K | Phase 2 |
| max.considered.p2 | 1000000 | P2 Max RS Family Check Per K | Phase 2 |
| max.stored.p2 | 10 | P2 Num Top RS Stored Per K | Phase 2 |
| max.nrs.p3 | 200000 | P3 Max RS Evaluated Per K | Phase 3 |
| max.stored.p3 | 100 | P3 Num Top RS Stored Per K | Phase 3 |
| k.max.4 | 40 | P4 Max K | Phase 4 |
| max.nrs.p4 | 100000 | P4 Max RS Evaluated | Phase 4 |
| max.stored.p4 | 100 | P4 Num Top RS Stored Per K | Phase 4 |
| gc.iter | 100 | Num CG Iterations | Generalized Core |
| gc.eval | 100 | Num RS Per GC | Generalized Core |
| Total Time | 54.4 | Total Computational Time | Timing |

#### 1.2 Timing breakdown (in minutes)

| P1 | PS | P2 | P3 | P4 | GC | Total |
|---|---|---|---|---|---|---|
| 3.29 | 6.67 | 6.36 | 13.7 | 4.62 | 19.8 | 54.4 |

### 2 Heatmaps of D and P

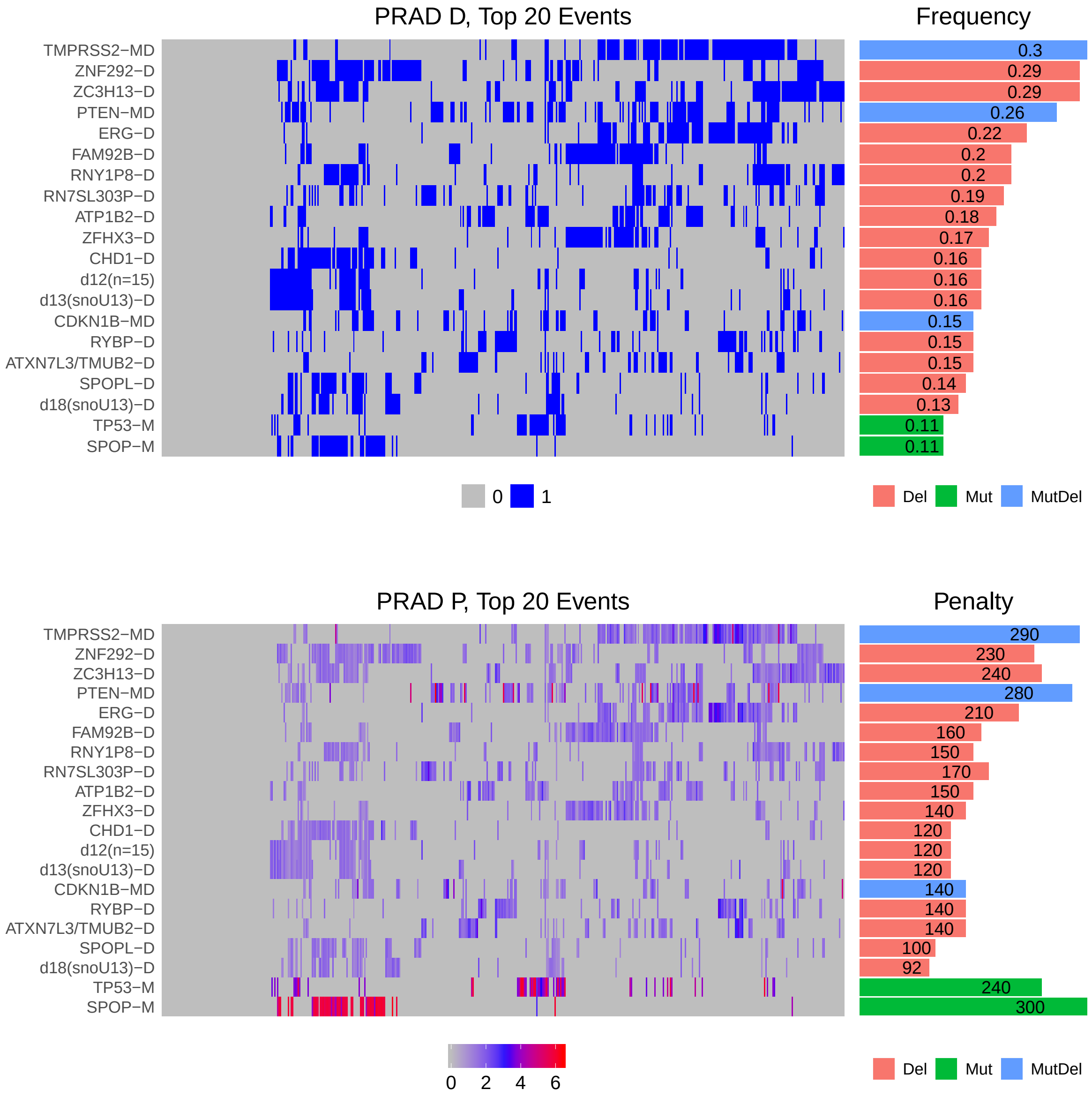

Figure 1: Heatmap of D and P. Top: binary representation of top 20 most frequent events. Top right shows event types and frequencies across the population. Bottom: Penalty matrix for top 20 most frequent events. Bottom right shows total event penalties. Events are ordered by frequency, from top to bottom. Samples are ordered using hierarchical clustering. Wild-type events indicated in grey. Event suffix -M: mutation, -A: amplification, -D: deletion, -MD: mutDel, -MA: mutAmp.

#### 3 Summary of K Best Rule Sets

##### 3.1 Performance and coverage of best rule sets

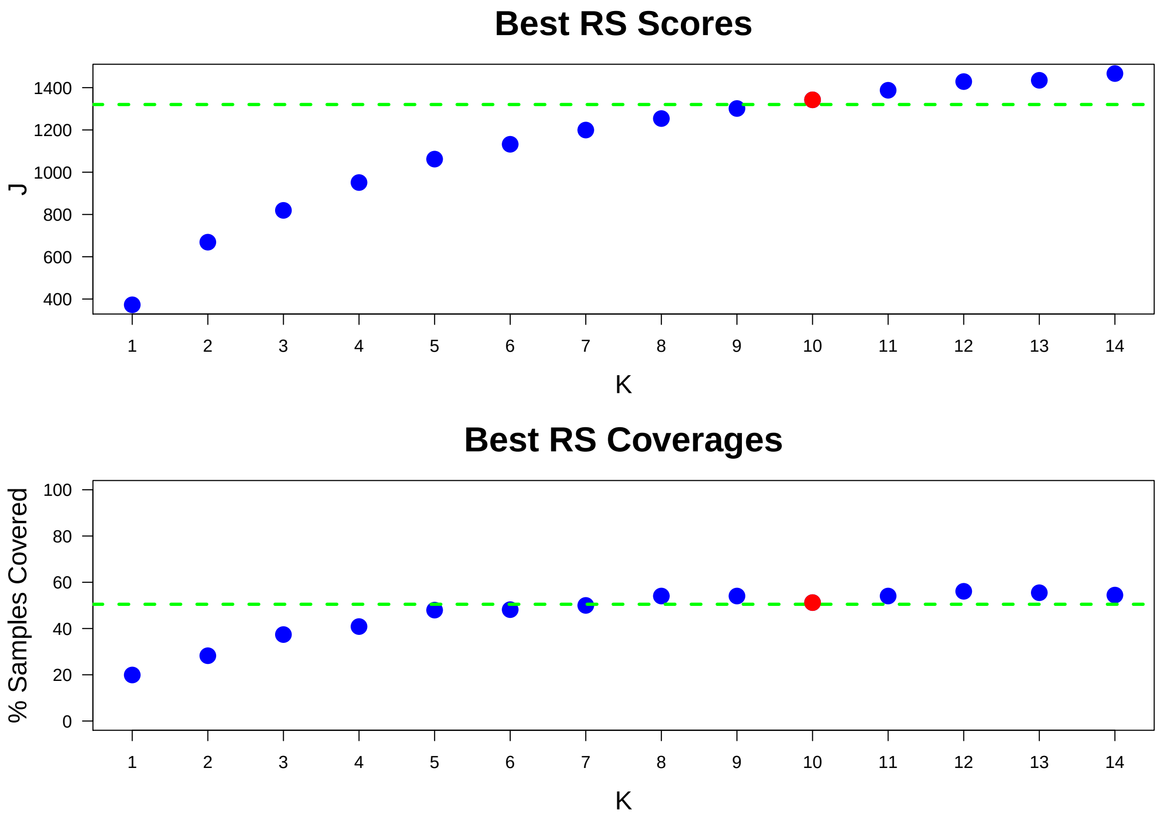

Figure 2: Performance and Coverage of Best Filtered RS for K. The core was determined to be the smallest RS that achieved at least 90% of maximum coverage and at least 90% of maximum performance.

#### 3.2 Table of rules that appear in any best rule set

| IR | Rule | PC | Ks |
|---|---|---|---|
| 2 | FAM92B-D+ZFHX3-D | 14 | 3-14 |
| 1 | ERG-D+TMPRSS2-MD | 20 | 1-9 |
| 3 | RNY1P8-D+ZC3H13-D | 16 | 5-13 |
| 5 | PTEN-MD+TP53-M | 5.3 | 4-11 |
| 9 | FOXA1-M+ZNF292-D | 3.9 | 8-14 |
| 11 | CHD1-D+SPOP-M+ZNF292-D | 6.5 | 8-14 |
| 8 | SPOP-M+ZNF292-D | 8.3 | 2-7 |
| 14 | PTEN-MD+RN7SL303P-D | 7.1 | 10-14 |
| 19 | d12(n=15)+d13(snoU13)-D+SPOP-M+ZNF292-D | 3.3 | 10-14 |
| 4 | ERG-D+PTEN-MD+TMPRSS2-MD | 9.3 | 10-12 |
| 6 | ATP1B2-D+TP53-M | 4.7 | 12-14 |
| 7 | PTEN-MD+TMPRSS2-MD | 13 | 7-9 |
| 10 | ERG-D+RYBP-D+TMPRSS2-MD | 5.9 | 12-14 |
| 20 | ATXN7L3/TMUB2-D+TMPRSS2-MD | 7.9 | 12-14 |
| 21 | CDKN1B-MD+ZNF292-D | 7.9 | 11-13 |
| 30 | PTEN-MD+RYBP-D | 6.5 | 12-14 |
| 68 | CHD1-D+SPOP-M+SPOPL-D+ZC3H13-D | 3.5 | 6-7 |
| 12 | d12(n=15)+d13(snoU13)-D | 12 | 8-9 |
| 17 | RYBP-D+TMPRSS2-MD | 8.5 | 10-11 |
| 18 | ATXN7L3/TMUB2-D+ERG-D+TMPRSS2-MD | 5.5 | 10-11 |
| 43 | ATP1B2-D+ERG-D+PTEN-MD+TMPRSS2-MD | 3.9 | 13-14 |
| 15 | ZC3H13-D+ZNF292-D | 12 | 14 |
| 26 | ERG-D+RNY1P8-D+TMPRSS2-MD+ZC3H13-D | 4.5 | 14 |
| 29 | ERG-D+TMPRSS2-MD+ZC3H13-D | 7.3 | 13 |
| 34 | ERG-D+FAM92B-D+TMPRSS2-MD | 6.3 | 14 |
| 39 | C11orf65-D+NNMT-D | 6.1 | 14 |

**ID** = Rule IDs, rules are numbered according to importance rank determined from phase 1
**PC** = Percent of samples covered **Ks** = Membership in best RS

### 4 Core Rule Set

**Core K = 10.**
**Core rule set coverage = 51.2%.**

#### 4.1 Table of core rule set rules

| IR | Rule | CR | SJR | SJ | NSC | NSA | PC | PA | FracA |
|---|---|---|---|---|---|---|---|---|---|
| 2 | FAM92B-D + ZFHX3-D | 3 | 8 | 224 | 68 | 50 | 14 | 10.0 | 0.74 |
| 3 | RNY1P8-D + ZC3H13-D | 2 | 6 | 250 | 79 | 35 | 16 | 7.1 | 0.44 |
| 4 | ERG-D + PTEN-MD + TMPRSS2-MD | 11.5 | 7 | 249 | 46 | 38 | 9.3 | 7.7 | 0.83 |
| 5 | PTEN-MD + TP53-M | 86 | 35 | 158 | 26 | 26 | 5.3 | 5.3 | 1.00 |
| 9 | FOXA1-M + ZNF292-D | 182.5 | 102.5 | 111 | 19 | 17 | 3.9 | 3.5 | 0.89 |
| 11 | CHD1-D + SPOP-M + ZNF292-D | 47.5 | 3 | 282 | 32 | 20 | 6.5 | 4.1 | 0.62 |
| 14 | PTEN-MD + RN7SL303P-D | 37.5 | 69 | 130 | 35 | 15 | 7.1 | 3.0 | 0.43 |
| 17 | RYBP-D + TMPRSS2-MD | 17.5 | 26 | 168 | 42 | 20 | 8.5 | 4.1 | 0.48 |
| 18 | ATXN7L3/TMUB2-D + ERG-D + TMPRSS2-MD | 78.5 | 41 | 155 | 27 | 15 | 5.5 | 3.0 | 0.56 |
| 19 | d12(n=15) + d13(snoU13)-D + SPOP-M + ZNF292-D | 250 | 29.5 | 163 | 16 | 16 | 3.3 | 3.3 | 1.00 |

**ID** = Rule IDs, rules are numbered according to importance rank determined from phase 1
**CR** = Coverage rank
**SJ/SJR** = Single rule performance / performance rank
**NSC/PC** = Number/percent of samples covered
**NSA/PA** = Number/percent of samples assigned
**FracA** = Fraction of covered samples assigned to rule

##### 4.2 Event breakdown of core rule set

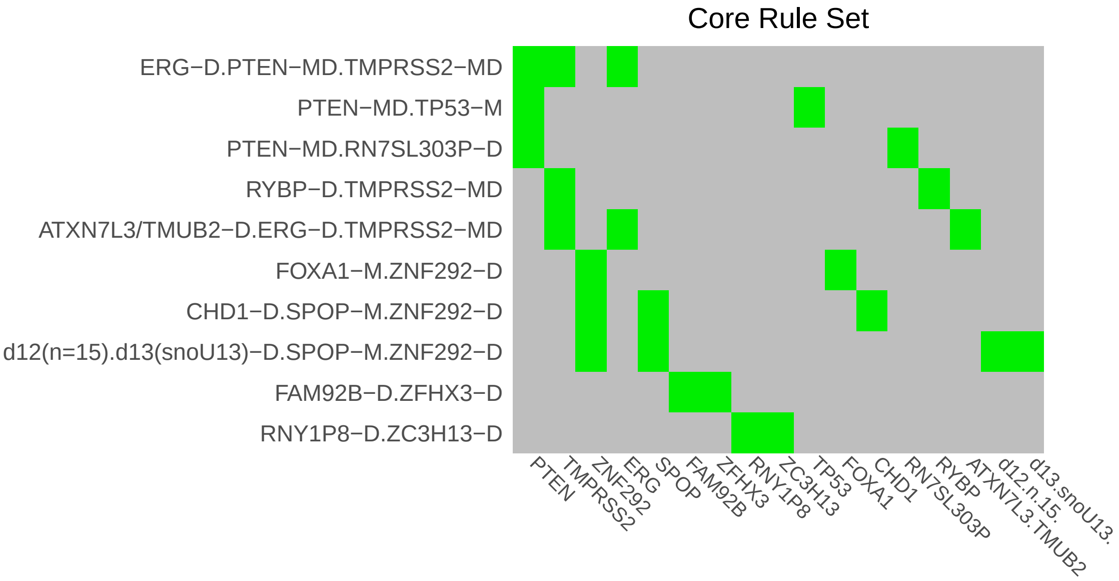

Figure 3: Visualization of core rule set. Rows are rules, columns are events. Events are ordered from left in decreasing rule membership frequency.

##### 4.3 Core Rule Set Penalties: Before and After Assignment

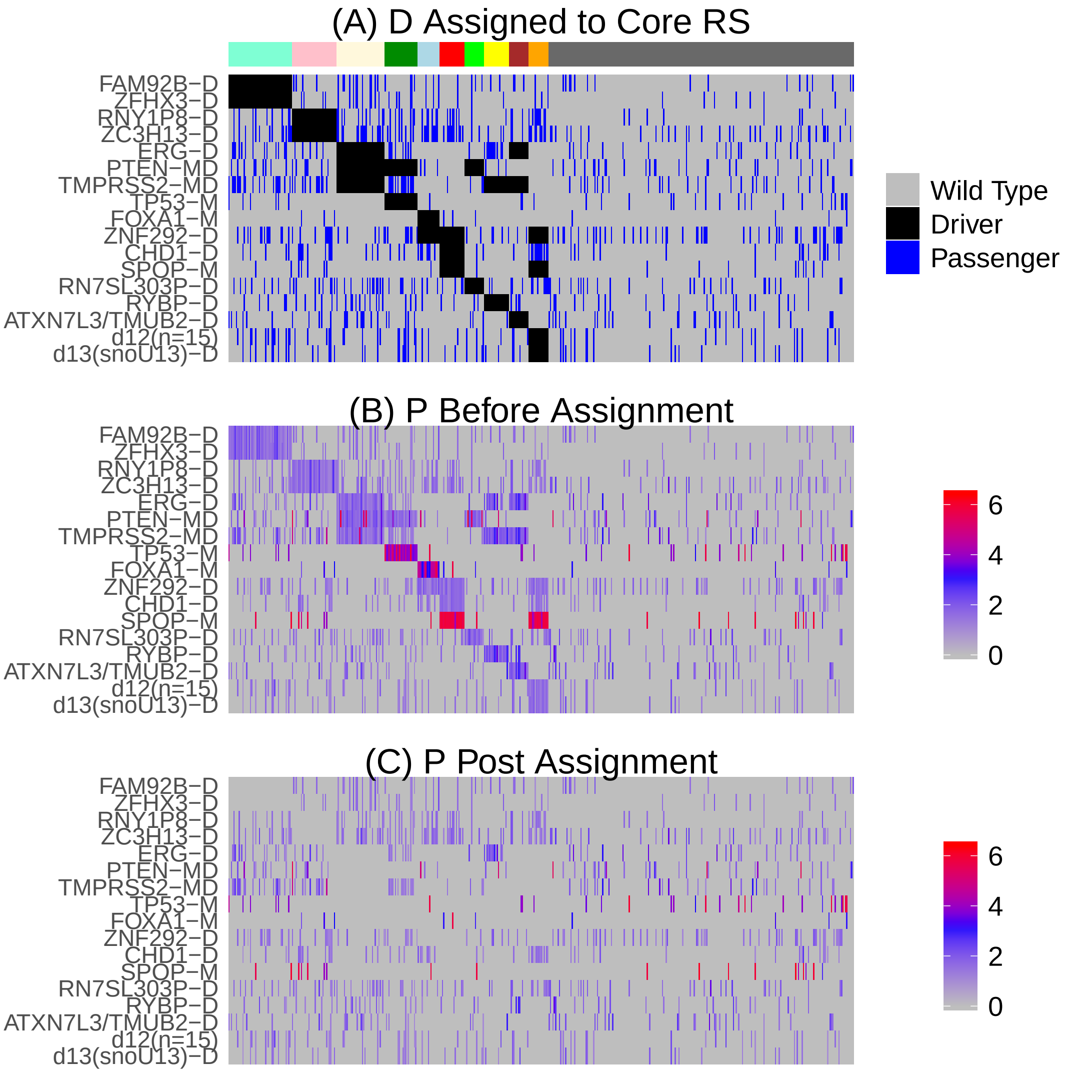

Figure 4: Heatmaps of D and P under core rule set assignment. Events are ordered by frequency. Samples are ordered according rule set membership, as indicated by the color bar. The dark grey group of samples are not assigned to any rule. A) For each sample, assigned events are designated as drivers and are shown in black, unassigned events are shown in blue and are assumed to be passengers. B-C) Heatmap of P before and after assignment to core rule set.

### 5 Generalized Core Analysis

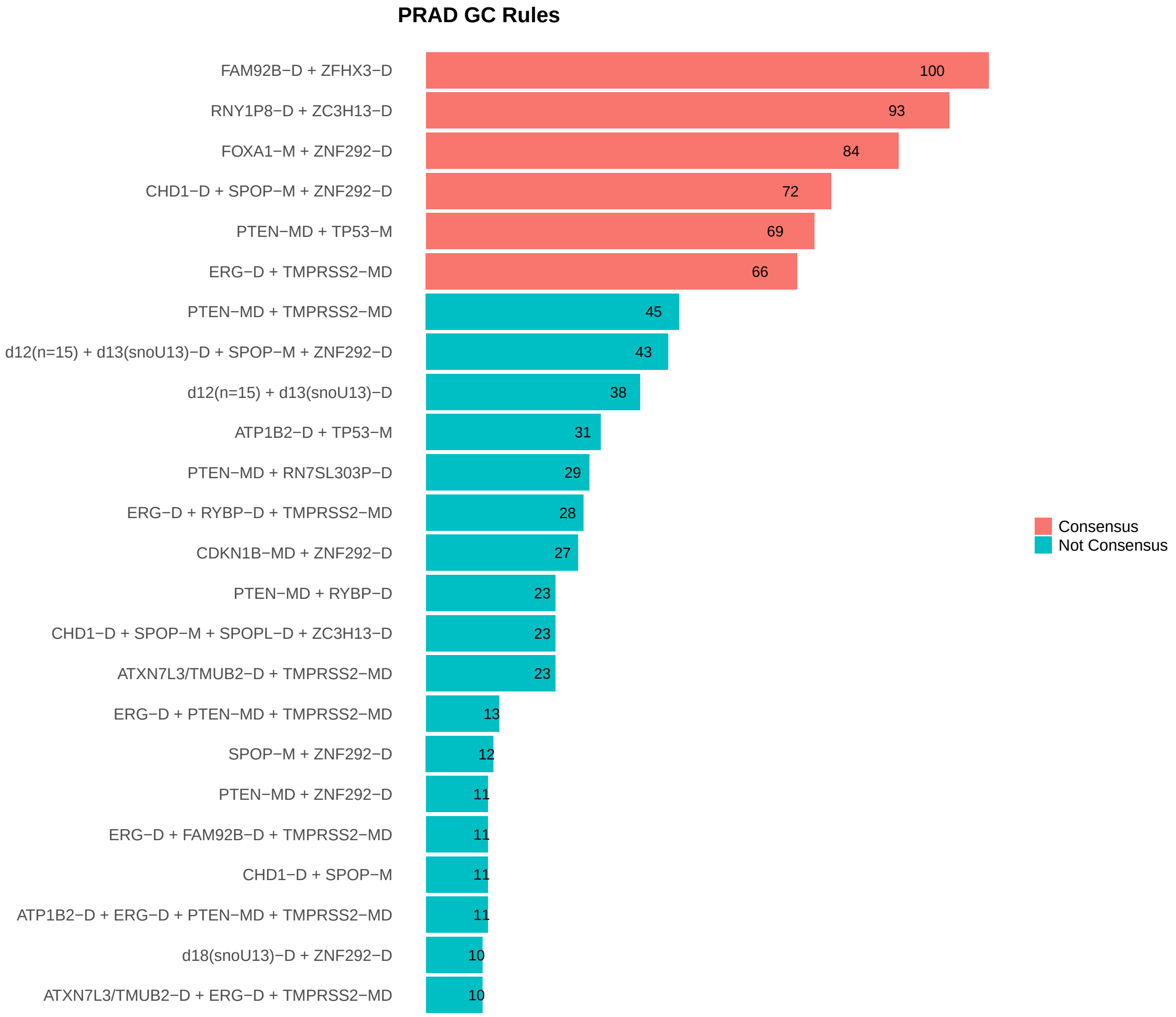

Figure 5: GCRs

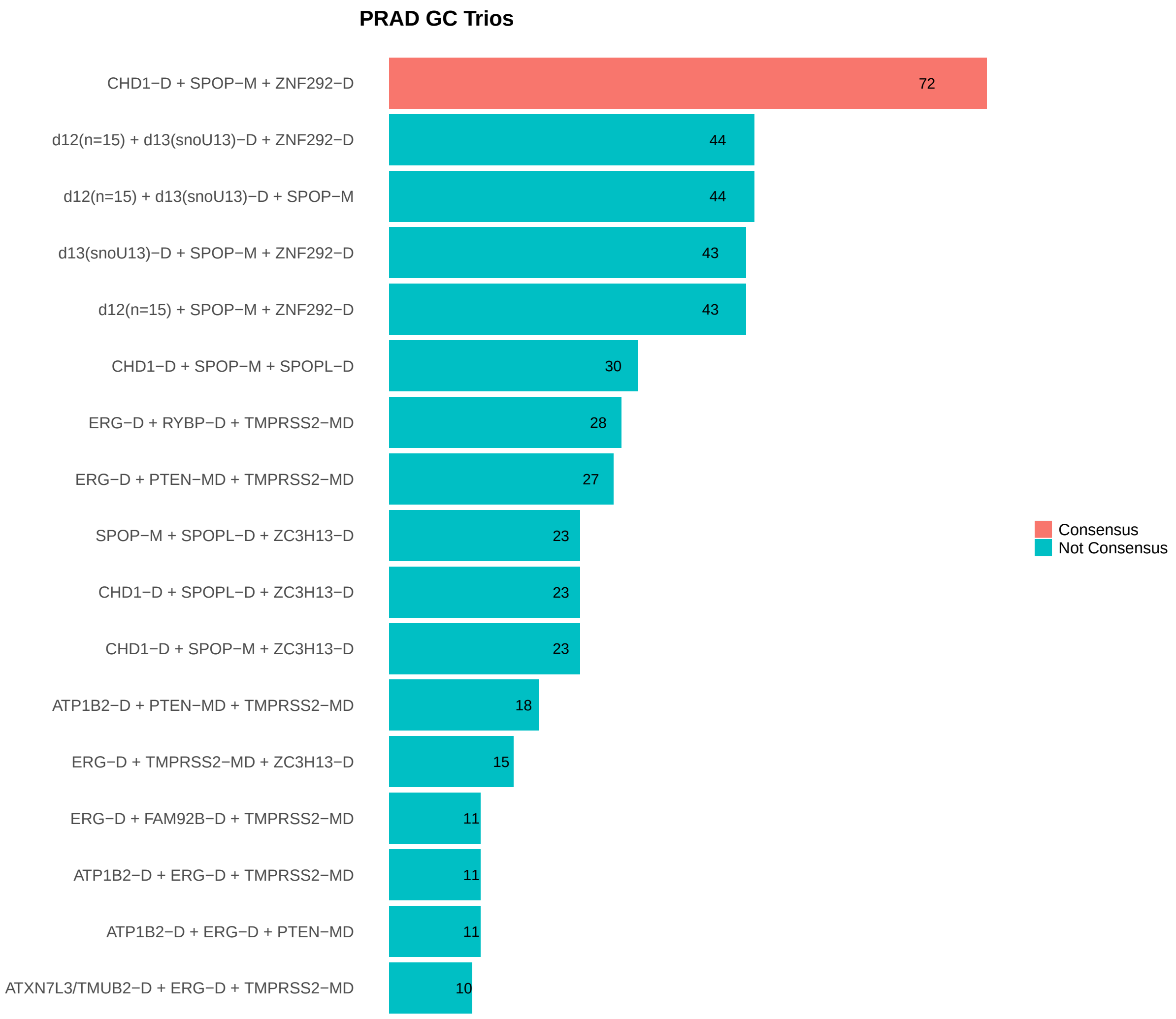

Figure 6: GCTs

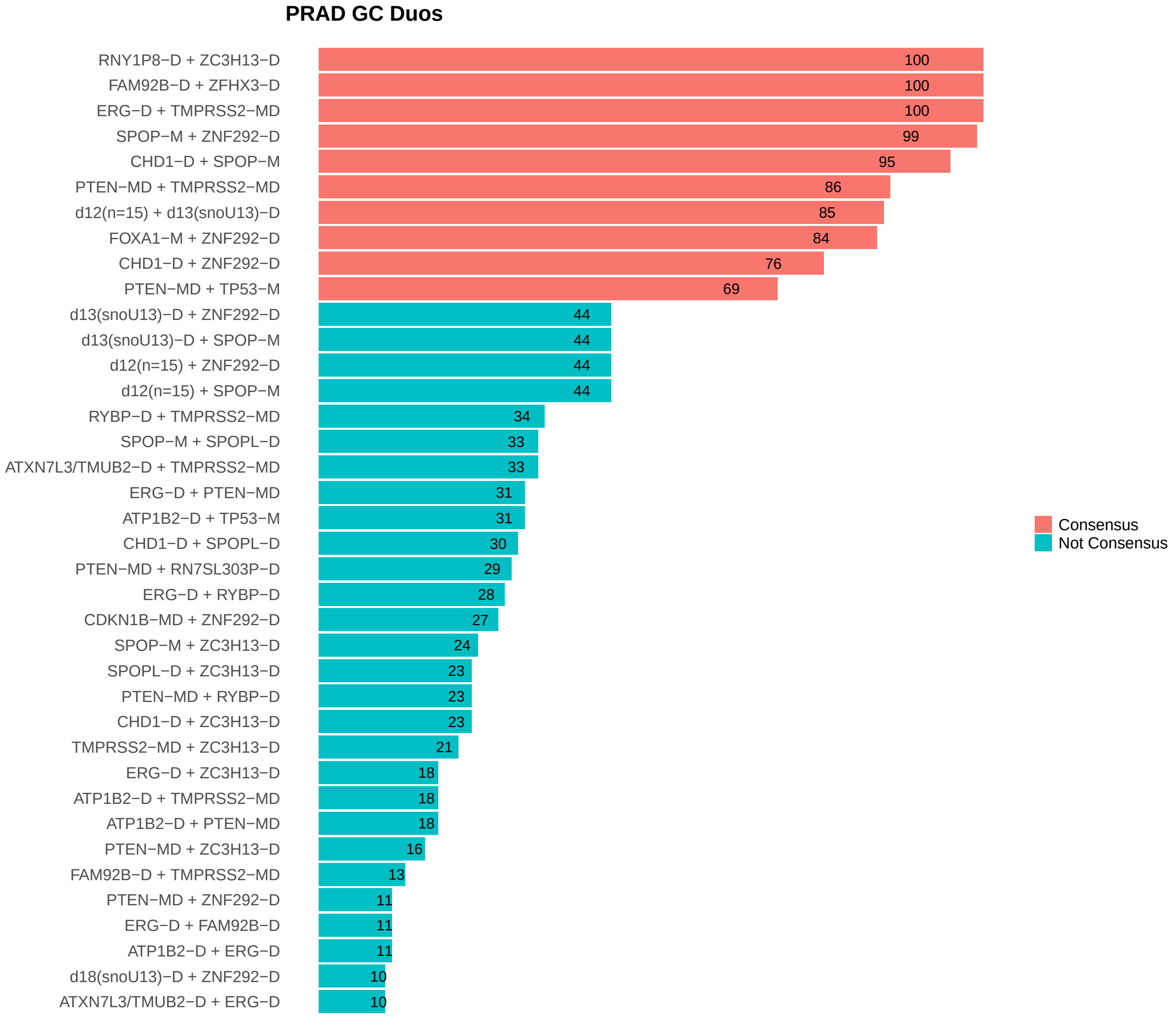

Figure 7: GCDs

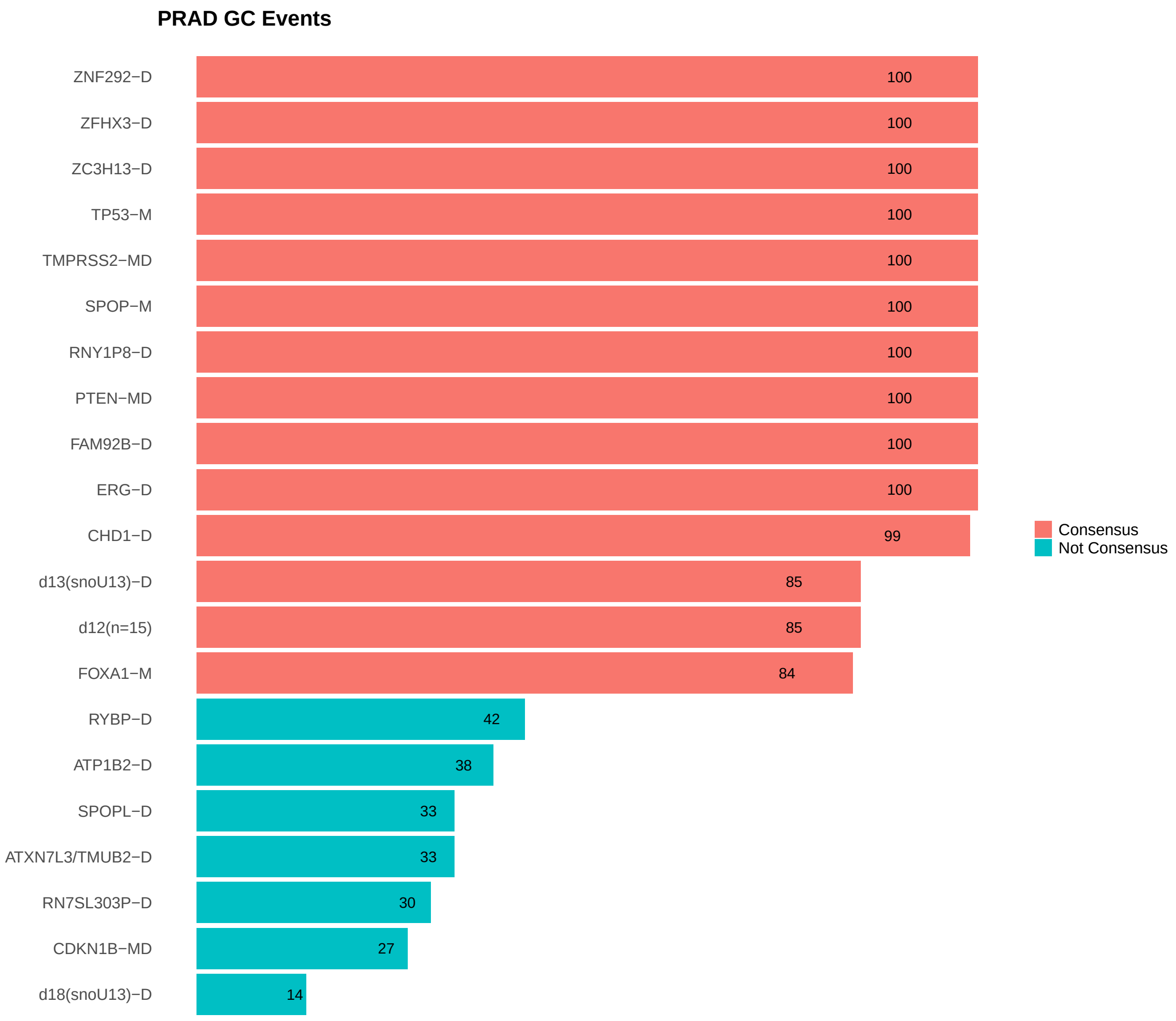

Figure 8: GCEs

### 6 Combined Core Table

| Rule | Core_Type | P1_Rank | Confidence | Coverage | SJ | Fraction_Assigned |
|---|---|---|---|---|---|---|
| FAM92B-D + ZFHX3-D | Both | 2 | 100 | 13.8% (r=3) | 296 (r=8) | 0.74 |
| RNY1P8-D + ZC3H13-D | Both | 3 | 93 | 16.1% (r=2) | 282 (r=6) | 0.44 |
| FOXA1-M + ZNF292-D | Both | 9 | 84 | 3.86% (r=187) | 207 (r=102) | 0.89 |
| CHD1-D + SPOP-M + ZNF292-D | Both | 11 | 72 | 6.5% (r=49) | 195 (r=3) | 0.62 |
| PTEN-MD + TP53-M | Both | 5 | 69 | 5.28% (r=85) | 258 (r=36) | 1 |
| d12(n=15) + d13(snoU13)-D + SPOP-M + ZNF292-D | Core | 19 | 43 | 3.25% (r=256) | 179 (r=29) | 1 |
| PTEN-MD + RN7SL303P-D | Core | 14 | 29 | 7.11% (r=37) | 186 (r=69) | 0.43 |
| ERG-D + PTEN-MD + TMPRSS2-MD | Core | 4 | 13 | 9.35% (r=12) | 275 (r=7) | 0.83 |
| ATXN7L3/TMUB2-D + ERG-D + TMPRSS2-MD | Core | 18 | 10 | 5.49% (r=81) | 182 (r=42) | 0.56 |
| RYBP-D + TMPRSS2-MD | Core | 17 | 6 | 8.54% (r=18) | 183 (r=26) | 0.48 |
| ERG-D + TMPRSS2-MD | conGCR | 1 | 66 | 19.9% (r=1) | 373 (r=1) | - |

### 7 Dictionary of Copy Number Events

| CNV | Genes | Event_Name |
|---|---|---|
| d1 | ZNF292 | ZNF292-D |
| d2 | ZC3H13 | ZC3H13-D |
| d5 | ERG | ERG-D |
| d6 | FAM92B | FAM92B-D |
| d7 | RNY1P8 | RNY1P8-D |
| d8 | RN7SL303P | RN7SL303P-D |
| d9 | ATP1B2 | ATP1B2-D |
| d10 | ZFHX3 | ZFHX3-D |
| d11 | CHD1 | CHD1-D |
| d12 | snoU13\|ENSG00000238451.1, GTF2H2B, RN7SL9P, snoU13\|ENSG00000238740.1, GUSBP3, RN7SL616P, RN7SL476P, GTF2H2, NAIP, SMN1, SMN2, SERF1A, GTF2H2C, SERF1B, OCLN | d12(n=15) |
| d13 | snoU13\|ENSG00000238717.1 | d13(snoU13)-D |
| d14 | RYBP | RYBP-D |
| d16 | ATXN7L3, TMUB2 | ATXN7L3/TMUB2-D |
| d17 | SPOPL | SPOPL-D |
| d18 | snoU13\|ENSG00000238860.1 | d18(snoU13)-D |
| d20 | NNMT | NNMT-D |
| d21 | C11orf65 | C11orf65-D |

Dictionary of important CNVs. Copy number events that are within any of the best K rule sets, or within the generalized core rules, or among the 20 most frequent events.
